# Efficient Design of Affilin^®^ Protein Binders for HER3

**DOI:** 10.1101/2025.04.02.646551

**Authors:** Anna M. Diaz-Rovira, Jonathan Lotze, Gregor Hoffmann, Chiara Pallara, Alexis Molina, Ina Coburger, Manja Gloser-Bräunig, Maren Meysing, Madlen Zwarg, Lucía Díaz, Victor Guallar, Eva Bosse-Doenecke, Sergi Roda

## Abstract

Engineered scaffold-based proteins that bind to concrete targets with high affinity offer significant advantages over traditional antibodies in theranostic applications. Their development often relies on display methods, where large libraries of variants are physically contacted with the desired target protein and pools of binding variants can be selected. Herein, we use a combined artificial intelligence/physics-based computational framework and phage display approach to obtain ubiquitin based Affilin^®^ proteins targeting the HER3 extracellular domain, a relevant tumor target. We demonstrate that the developed *in silico* pipeline can generate *de novo* Affilin^®^ proteins with high experimental success rate using a small training set of sequences (<1000 sequences). The classical phage display yielded primary candidates with low nanomolar affinities. These binders could be further optimized by phage display and computational maturation alike. These combined efforts resulted in four HER3 ligands with high affinity, cell binding, and serum stability that have theranostic potential.

## INTRODUCTION

The dysregulation of membrane-bound receptor tyrosine kinases (RTKs) plays a crucial role in cancer development, leading to enhanced proliferation, survival, migration, and other processes associated with cancer progression. The RTK class consists of the highly homologous epidermal growth factor receptor (EGFR), HER2 (erbB2), HER3 (erbB3), and HER4 (erbB4) (1,2). Expression levels of members of the human epidermal growth factor receptor (HER) family are elevated in multiple cancers (3). Numerous diagnostic and therapeutic compounds have been developed for the first two members leading to broadly applied drugs like Cetuximab and Trastuzumab. More recently HER3-targeting ligands e.g. Seribantumab and Patritumab emerged but none has been approved for tumor therapy yet. HER3 expression is an important factor in the development of resistance to treatment with anti-HER2 agents (4). Ligands against HER3 hold promise to act beside a first-line therapeutic in combination or salvage medication schemes (5).

Affilin^®^ proteins are small scaffold-based compounds with drug-like properties (6,7). Based on the Affilin^®^ technology high affinity tumor marker targeting molecules are selected by advanced phage display protocols (8). Primary hits can be further optimized to meet individual requirements for different applications in the diagnostic or therapeutic field by maturation or site-directed mutagenesis. Scaffold based biologicals overcome limitations of small molecules and antibodies alike (9). The intermediate size is a prerequisite for versatility, engineerability and modularity. It also opens up the route for *in silico* drug design and optimization.

*De novo* design of proteins that bind to a specific site of a target protein is an important challenge in biotechnology, with extensive applications ranging from therapeutics to diagnostics. The recent advances in computational methods, especially due to the surge of deep learning techniques for structural biology, are allowing the characterization of protein-protein interfaces (10), and the following utilization of this information to design binding proteins to specific locations of the receptor (11,12). Moreover, Large Language Models (LLMs), trained on diverse cross-species and cross-family protein sequences, have demonstrated efficient exploration of the protein sequence space and generation of *de novo* plausible protein sequences for a wide variety of protein families (13–16). This generalization capability stems from their training on large datasets, often exceeding 15,000 sequences per protein family even for small-sized language models (17). However, when datasets are smaller—fewer than 1,000 sequences—challenges arise due to the risk of overfitting or memorization rather than meaningful generalization. Fine-tuning state-of-the-art foundation protein language models (PLMs) on such limited datasets may exacerbate these risks, as the dataset size may be insufficient even for fine-tuning tasks (17). In these limited data scenarios, approaches that combine less data demanding artificial intelligence (AI) methods with more consolidated physic-based Molecular Modeling (MM) techniques can boost the chances of successfully designing binders for the target protein (18).

On their side, MM techniques are fundamental in understanding the complex nature of protein-protein interactions (PPIs). These techniques, such as molecular dynamics (MD) simulations (19), Monte Carlo (MC) methods (20,21), and protein-protein docking (PPD) algorithms (22), provide a detailed atomic-level view of their interactions and dynamics. These techniques can estimate the binding affinity between the protein partners (23), identify critical residues at the binding interface (23), and predict the conformational changes upon binding (24). Consequently, MM techniques provide a framework to validate (and even reinforce) AI-designed sequences, ensuring their stability and functionality (25). However, to scale the usage of these classical techniques, it is essential to streamline the prediction of protein-protein complexes by finding a tradeoff between accuracy and computational cost.

Experimental methods, such as phage display, serve as a critical complement to the protein designs coming from the AI-MM pipeline, addressing challenges that computational approaches alone cannot solve. Phage display libraries enable the empirical generation and *in vitro* screening of billions of protein variants, providing an unparalleled ability to identify high-affinity binders. This approach is especially advantageous as a starting point when computational methods face limitations, such as difficulties in effectively scanning large or highly complex targets. By offering experimentally validated leads, phage display bridges gaps left by AI and MM techniques and provides a foundation for further computational refinement. However, this method comes with several drawbacks, including that it can be labor-intensive, requiring extensive experimental optimization, and the binders identified *in vitro* may not always translate to functional performance under *in vivo* conditions. Despite these challenges, integrating experimental methods with AI and MM ensures finding new binders, where empirical insights validate and enhance theoretical predictions, facilitating the design of proteins with robust binding capabilities and practical applications.

In this paper, we employ AI-MM computational design methods alongside classical phage display to identify HER3-binding Affilin^®^ proteins. The promising candidates discovered through these approaches were then computationally optimized and experimentally validated for their binding efficacy towards HER3. On the one hand, for the *in silico* approach we trained a Variational Autoencoder (VAE) with a small-sized sequence dataset (<1,000 sequences) of Affilin^®^ proteins. Since the AI-generated sequences were not target-specific, we set up a robust MM pipeline to screen the *de novo* sequences against our target of interest. This pipeline strategically reduces the number of candidates by progressively integrating more accurate MM techniques, transitioning from PPD to MC and MD simulations for accurate binding predictions, leading to distinct HER3 binding Affilin^®^ proteins with a dissociation constant (KD) below 100 nM. On the other hand, for the phage display campaigns we used a panel of nine different library designs and two target formats in the presence and absence of serum yielding several promising candidates with up to single-digit nanomolar affinities which could be further optimized by phage display and computational maturation.

## RESULTS

### Combining AI and MM to design de novo Affilin® proteins for HER3

We explored combining AI algorithms with our MM pipeline (see benchmark in Supplementary Note 1) to enhance the success rate of generating new Affilin^®^ proteins. Specifically, we used a VAE algorithm to generate *de novo* Affilin^®^ protein sequences, which were then ranked using the herein-described MM pipeline to prioritize sequences for the experimental assays (Figure 1). Despite the constrained size of the training set, the integration of the VAE model with the MM pipeline facilitated the identification of a batch of 12 promising candidate sequences, among which 2 binders showed experimental KD < 100 nM. This highlights the efficiency of combining AI + MM approaches in increasing success-rate.

**Figure 1.**
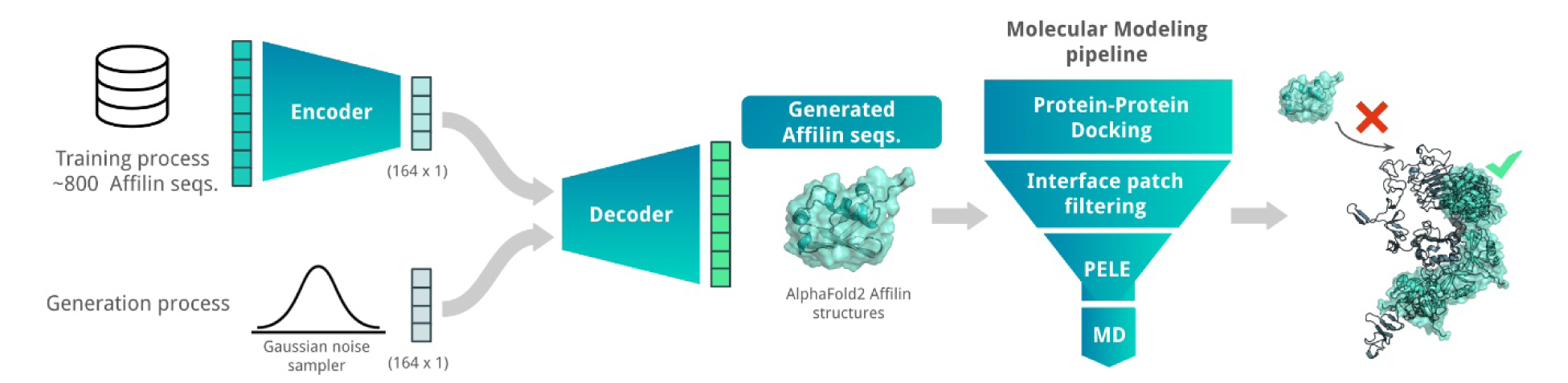
Scheme of the AI + MM pipeline utilized for generating and identifying strong Affilin^®^ proteins targeted towards a specific protein, in this case HER3. The AI block consisted of a VAE that was trained on 852 non-identical Affilin^®^ protein sequences. *de novo* Affilin^®^ protein sequences were generated by using Gaussian random noise as input for the decoder. The generated sequences were filtered based on sequence length (between 72 and 82 amino acid length) and sequence identity with the ubiquitin wild type (between 75 and 90% sequence identity). The resulting sequences were modeled with AlphaFold2 and sent to our MM pipeline (Figure SN1a). This pipeline strategically reduces the number of binding modes and progressively integrates more accurate MM techniques, transitioning from PPD to MC and MD simulations for accurate binding predictions. Its performance is assessed with two retrospective studies (see Supplementary Note 1).

### Selection of the machine learning algorithm for de novo sequence generation using a small-sized dataset

Despite the success of various Machine Learning (ML) algorithms in protein sequence generation (16,17,26), the constraints arising from a small-sized dataset make a rational selection of the ML algorithm imperative. In this context, VAEs emerged as the preferred choice over more complex architectures like transformers. VAEs are particularly advantageous in this context for their efficiency in handling sparse data and their ability to capture essential distributions of highly similar sequences through a compressed latent space. This efficiency is particularly valuable, enabling the generation of unique biologically plausible sequences within the constraints of our dataset. The decision to adopt VAEs is further justified by their intrinsic regularization mechanism, addressing the crucial concern of overfitting given the dataset’s size and similarity which mitigates overfitting, a critical consideration given our dataset’s size and similarity (27). More in detail the regularization enabled generating versatile sequences showing promise for real-world applicability beyond the sequences from the training set.

In contrast, transformers, while powerful in modeling complex sequence dependencies, demand larger datasets and significant computational resources, and their effectiveness in avoiding overfitting and in generating original sequences from such a specialized dataset is less certain (28). Consequently, transformer architectures were considered less suitable for our specific scenario than VAEs.

### Generation of de novo Affilin^®^ protein sequences for HER3

The trained VAE effectively encodes the known protein data into a compressed latent space. This encoding effectively distilled the inherent patterns and intricacies of our initial sequences, offering a richer, denser representation of the limited data we had. From this optimized latent space, we generated 500 distinct Affilin^®^ protein sequences. While these sequences were influenced by the patterns of the initial data, the VAE’s sampling method using Gaussian noise enabled us to design *de novo* protein sequences. One of the most notable capabilities we observed was the VAE’s ability to interpolate between distinct data points from our initial starting point (Figure 2A). This led to the creation of sequences that incorporated features from multiple proteins, giving us a varied and diverse output.

**Figure 2.**
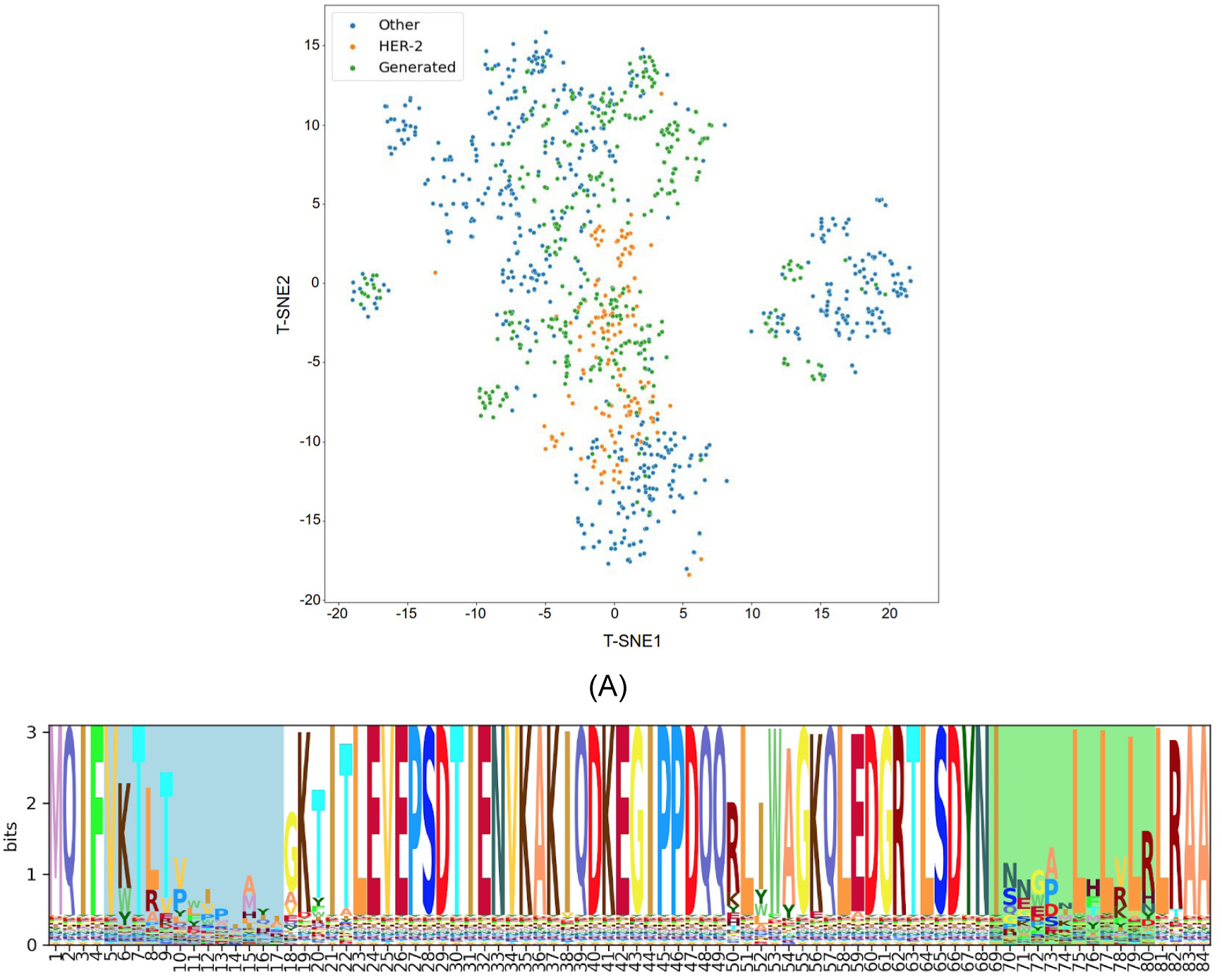
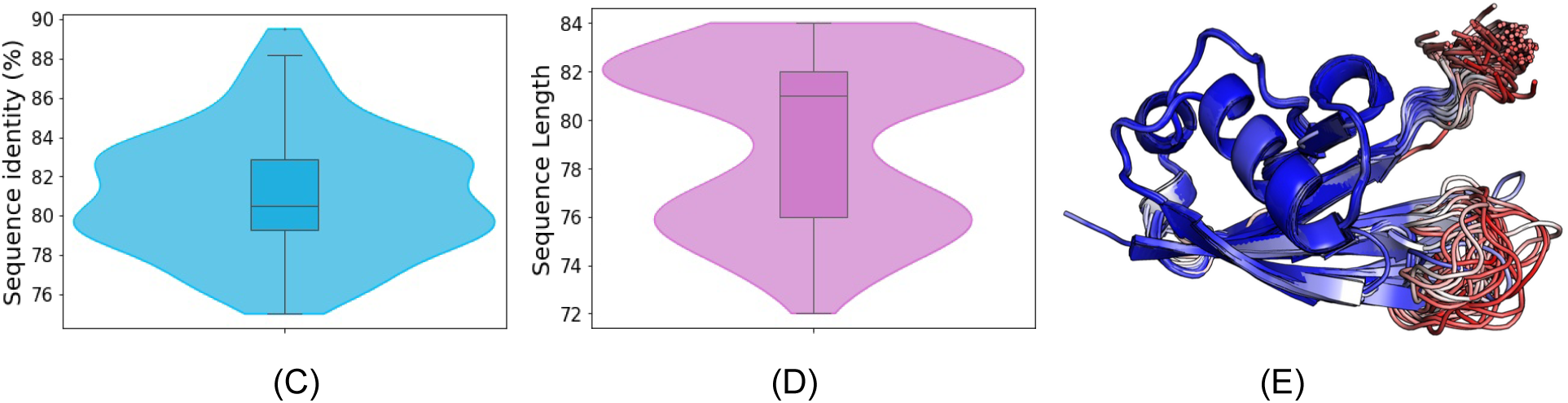
Analysis of de novo Affilin^®^ binder candidates targeting HER3. (A) t-distributed stochastic neighbor embedding (t-SNE) representation of the available sequence space of the Affilin^®^ proteins with highlighted HER2 Affilin^®^ binders and the 500 generated sequences. (B) Sequence logo of the 112 Affilin^®^ proteins selected to target HER3 with the 2 variable loops highlighted (N-terminal loop in blue and the C-terminal in green). (C) Sequence identity distribution to ubiquitin of the 112 selected sequences. (D) Length distribution of the selected sequences. (E) Superposition of the AlphaFold2 structures of the 101 selected Affilin^®^ proteins that pass the pLDDT filter (no region of the protein with a pLDDT < 50) colored by pLDDT. Regions in blue are highly confident regions, while regions in red indicate low confidence.

The resulting Affilin^®^ protein candidates underwent further filtration based on two criteria: i) a sequence identity to the ubiquitin wild type (WT) ranging between 75-90%, and ii) a sequence length of 72-84 residues (with the ubiquitin WT having a length of 76 residues). This filtration process yielded 112 designs, exhibiting an average identity of 81.15% to the ubiquitin WT and an average length of 79 amino acids (refer to Figures 2C and 2D, respectively).

Subsequently, all the 112 selected sequences were subjected to modeling using AlphaFold2. The overall confidence in the generated sequences models was high (pLDDT > 90), as indicated by the pLDDT values with an average of 90.46. However, out of these 112 sequences, 11 had some regions of the protein with a pLDDT below 50, so they were discarded to aim for overall good-quality models according to the confidence score. As illustrated in Figure 2E, the 101 AlphaFold2 models of the generated Affilin^®^ proteins clearly showed that the structural variability was coming from the first loop (N-terminal) and the last loop at the C-terminus, consistently with the sequence variability (Figure 2B) and the pLDDT scores in those regions (Figure 2E).

### Identification of promising de novo sequences with our computational pipeline

Having confident AlphaFold2 models for the 101 selected Affilin^®^ protein sequences, we employed our MM pipeline (Figure SN1a) to estimate their binding strength towards HER3.

In the PPD analysis, 73 sequences were excluded based on the mean of the top 50 PyDock binding energies of conformationally distinct poses (L-RMSD > 2Å). Specifically, Affilin^®^ protein candidates with a mean binding energy towards HER3 higher than the average binding energy across all systems were discarded (Figure S1). The remaining 39 Affilin^®^ protein sequences, which passed this criterion, underwent refinement with PELE for more precise binding affinity estimation. Employing the same rationale as in the PPD analysis, an additional 13 sequences were rejected due to their mean energy of the top 10 PELE poses being above the average binding energy across all systems (Figure S2).

The remaining 26 Affilin^®^ protein sequences underwent MD simulation starting from PELE’s best binding energy pose to assess their binder affinity towards HER3. The computational results suggested that 6 Affilin^®^ sequences were no binders (NB) to weak binders (WB), 9 were medium binders (MB), and 11 were strong binders (SB) (Figure S3).

### Experimental validation of the de novo selected sequences

Among the 11 SBs and 1 WB selected for experimental validation, 10 were effectively purified. Even within this cohort, the expression varied drastically (Figure S4).

Within this group of 10 proteins, 7 demonstrated binding activity with KD ∼10 μM. Notably, 2 out of these 7 proteins exhibited medium to strong binding, with one achieving a KD below 50 nM (Table 1). One drawback was the lack of single-digit nanomolar affinity and cell binding of the in silico identified ligands.

**Table 1.** Summary of binding affinity towards HER3-Fc data for in silico designed Affilin ^®^ proteins obtained with different methods. Lab scale purified Affilin^®^ proteins were used in a concentration-dependent manner for SPR measurement, which was performed on a Sierra SPR-32 (Bruker) with HER3-Fc immobilized on a Protein A-coated chip. Cell binding was measured on a Guava easyCyte 5HT FACS using HEK293-HER3-sc87 cells and Affilin^®^ proteins in a concentration-dependent manner. Sequences with hyphens were not expressed. The best Affilin^®^ designs are indicated in bold. Table abbreviations: NB – no binding was detected.

| Affilin<sup>®</sup> | SPR (lab scale)<br>[nM] | Cell binding to HEK293-HER3-sc87<br>[nM] |
| --- | --- | --- |
| Comp-1 | - | NB |
| Comp-2 | 2061 | NB |
| Comp-3 | - | NB |
| Comp-4 | NB | NB |
| Comp-5 | NB | NB |
| Comp-6 | 10000 | NB |
| Comp-7 | 462 | NB |
| Comp-8 | 5381 | NB |
| **Comp-9** | **29.8** | NB |
| Comp-10 | 1488 | NB |
| **Comp-11** | **63.2** | NB |
| Comp-12 | NB | NB |

### In vitro identification of de novo HER3-binding Affilin^®^ proteins using phage display and high-throughput screening

In parallel to the *in silico* generated HER3 targeted Affilin^®^ proteins, we performed an *in vitro* selection using phage display technology. We obtained HER3 binding phage pools after three selection rounds and transferred them to a high-throughput screening to identify specific binding Affilin^®^ proteins. Proteins fulfilling the required specifications were nominated as hit and expressed, purified and analyzed using the extra- cellular domain (ECD) of the HER3 protein or by using HER3-expressing cell lines. The general workflow consists of four main steps (Figure 3).

**Figure 3.**
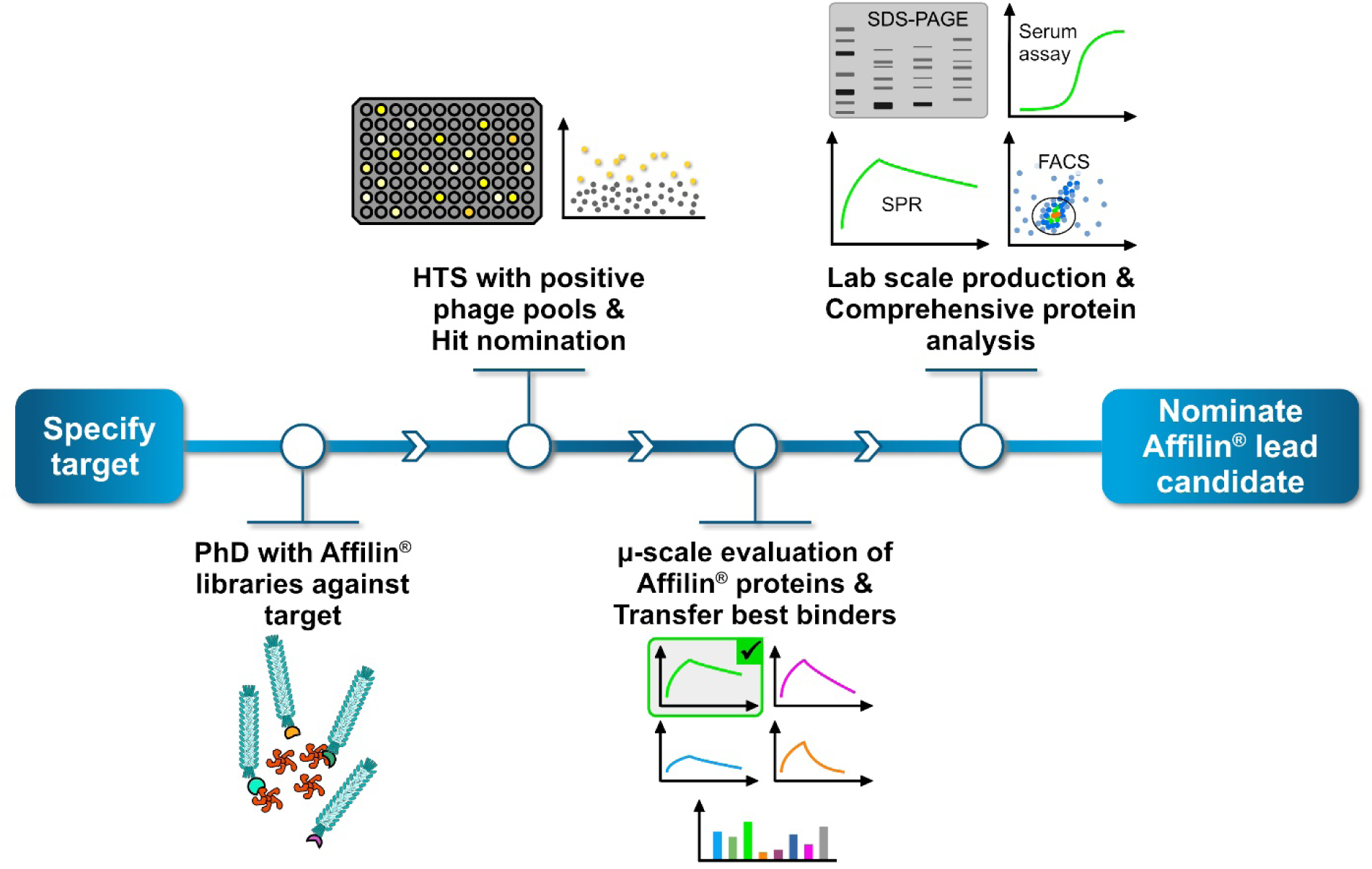
General workflow to generate and identify specific Affilin^®^ proteins versus a given target. New projects start with validating and specifying a target. The first step consists of a selection process using phage display (PhD) with different high diverse Affilin^®^ libraries against the target. Positive binding phage pools will be transferred to a high-throughput screening (HTS), the second step in the workflow. Up to 15.000 single clones can be analyzed, each representing a unique Affilin^®^ protein. Specific binding variants are nominated as hit. The third step consists of a first evaluation of each hit applying a µ-scale expression and purification in a 96-well format including a fast small-scale purification by using PhyNexus columns. Binding affinity is tested with surface plasmon resonance (SPR) und flow cytometry (FACS) measuring techniques. The most promising Affilin^®^ variants are transferred to step four, which includes an upscaling process in the protein expression and purification using *e.g.* gel filtration and affinity chromatography systems. The highly purified samples undergo a comprehensive protein analysis including concentration- dependent SPR, FACS, serum stability assays, DSF (differential scanning fluorimetry) and many more measurements to identify the best Affilin^®^ lead candidate.

### Primary selection with Affilin^®^ libraries using phage display technology

We performed a phage display using highly complex Affilin^®^ libraries with a diversity of more than 10^10^ variants, which were fused to the pIII phage protein using a derivative of the pCD87SA phagemid. The setup of the primary selection consisted of nine different Affilin^®^ libraries that were incubated with the HER3 target protein over three rounds with reducing amount of target, starting from 180 nM to 8 nM. Additionally, a serum incubation of the phage pools preceding round two and three was performed. The obtained phage pools from each round were tested for recognizing the target in an ELISA setup. Eight libraries revealed specific HER3 binding phage pools (Figure S5).

The HER3 positive phage pools were subcloned into expression vectors for the high-throughput screening.

### Identification of HER3-binding Affilin^®^ proteins via high-throughput screening

The high-throughput screening (HTS) is divided into three steps – a primary screen, a secondary screen and a µ-scale protein analysis. Relevant pools were subcloned into an expression vector, generating Affilin ^®^ proteins with an N-terminally fusion to green fluorescent protein (GFP) and expressed in *E. coli* BL21 (DE3).

Single clones were picked, 8930 in total distributed over eight pools, for the primary screen. The ELISA-based high-throughput screening was performed with biotinylated HER3 as target protein and the respective expression supernatant of each clone as analyte. The binding of the Affilin^®^ proteins to the target protein was measured via GFP fluorescence. Overall, 3378 out of 8930 Affilin^®^ proteins could be nominated as primary hit and were transferred to a secondary screening (Figure S6).

The secondary screen utilized two target proteins to further evaluate the primary hits in an ELISA-based setup. The expression supernatants of each hit were incubated in mouse serum for 24 h at 37 °C and binding to biotinylated HER2 and biotinylated HER3 was tested. A specific binding to HER3 could be detected for 285 clones (Figure S7). To eliminate identical clones due to sequence enrichment during the selection, sequence analysis for all 285 clones was conducted and led to 87 unique Affilin^®^ proteins, which were finally selected for a µ-scale protein analysis.

The µ-scale analysis consists of a fast protein purification step by using a small-scale affinity matrix after which the binding behaviour of the samples can be evaluated by surface plasmon resonance (SPR) and cell binding. During this analysis, the 87 Affilin® proteins were additionally cloned into another expression vector containing only a C-terminal StrepTag. Four Affilin® proteins were nominated as primary lead variants with Affilin® Exp-1 as the best one, which already appeared to be as a strong binder with a KD < 10 nM (Table 2, Figure S8).

**Table 2.** Summary of binding affinity data towards HER3-Fc for in vitro selected Affilin^®^ proteins obtained with different methods. Either µ-scale or lab scale purified Affilin^®^ proteins were used in a concentration-dependent manner for SPR measurement, which was performed on a Sierra SPR-32 (Bruker) with HER3-Fc immobilized on a Protein A-coated chip (see also Figure 4). Serum stability assay in mouse or human serum was performed by an ELISA with HER3-Fc immobilized on 384-well plates and lab scale purified Affilin^®^ proteins (see also Figure 5). Table abbreviations: nd – not determined

| Affilin® | SPR (μ-<br>scale)<br>[nM] | SPR (lab<br>scale)<br>[nM] | Binding in mouse serum [nM] | | Binding in human serum [nM] | |
| --- | --- | --- | --- | --- | --- | --- |
|  |  |  | 0h | 24h | 0h | 24h |
| Exp-1 | 2.5 | 5.4 | 0.8 | 0.7 | 1.0 | 0.7 |
| Exp-2 | 8.6 | 19.4 | 11.1 | 10.3 | 21.1 | 12.4 |
| Exp-3 | 8.7 | 26.4 | nd | nd | nd | nd |
| Exp-4 | 5.9 | 17.1 | nd | nd | nd | nd |

### Validation of in vitro selected Affilin^®^ proteins

After the initial µ-scale analysis Exp-1, Exp-2, Exp-3 and Exp-4 were chosen for a more detailed analysis of lab scale samples. All variants showed good expression and solubility in *E. coli* BL21 (DE3) (Figure S9) and purification via StrepTag and polishing via size-exclusion chromatography resulted in abundant and pure protein. The chosen variants showed specific binding to HER3-Fc below 30 nM and were classified as SBs (Table 2, Figure 4) and Exp-1 had the highest affinity to HER3-Fc.

**Figure 4.**
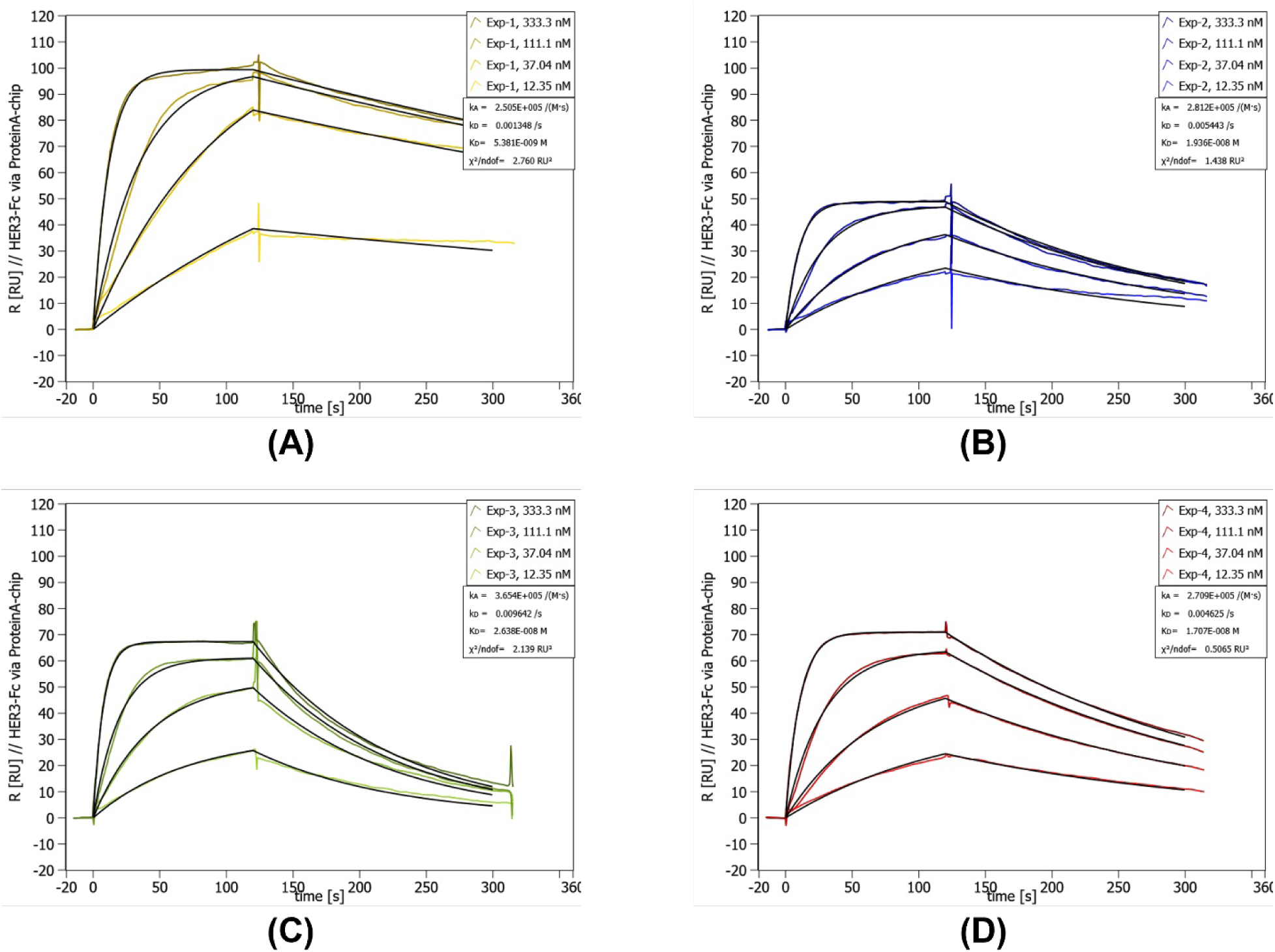
Concentration-dependent SPR data of lab scale purified Exp-1 to Exp-4. SPR was performed on a Sierra SPR-32 (Bruker) with a Protein A-coated chip on which the HER3-Fc was immobilized. The concentration of the curves is given in the upper right corner of each individual graph. Below the concentrations, the measured kinetic values are given. (A) SPR data for Exp-1. (B) SPR data for Exp-2. (C) SPR data for Exp-3. (D) SPR data for Exp-4.

The target affinity of Affilin^®^ proteins was further evaluated in a more biological environment using HER3- expressing cell lines. Binding to the cells was identified by FACS measurement with an AlexaFlour488- labeled anti-StrepTag antibody, which recognized the StrepTag-fused Affilin^®^ proteins. All four tested variants revealed HER3-specific binding. The interaction with HER3-overexpressing HEK293-cells resulted in a higher signal-to-noise ratio (x-fold) compared to SK-BR-3-cells expressing the HER3 at a lower level (Table S1). Furthermore, serum stability of Exp-1 and Exp-2 was tested by an ELISA in a concentration-dependent manner determining the affinity after incubation in mouse serum up to 24 h. Both variants showed a high affinity in the low nM range, which was not significantly influenced after serum incubation over 24 h at 37 °C (Table 2, Figure 5). Therefore, these variants were rated as serum stable.

**Figure 5.**
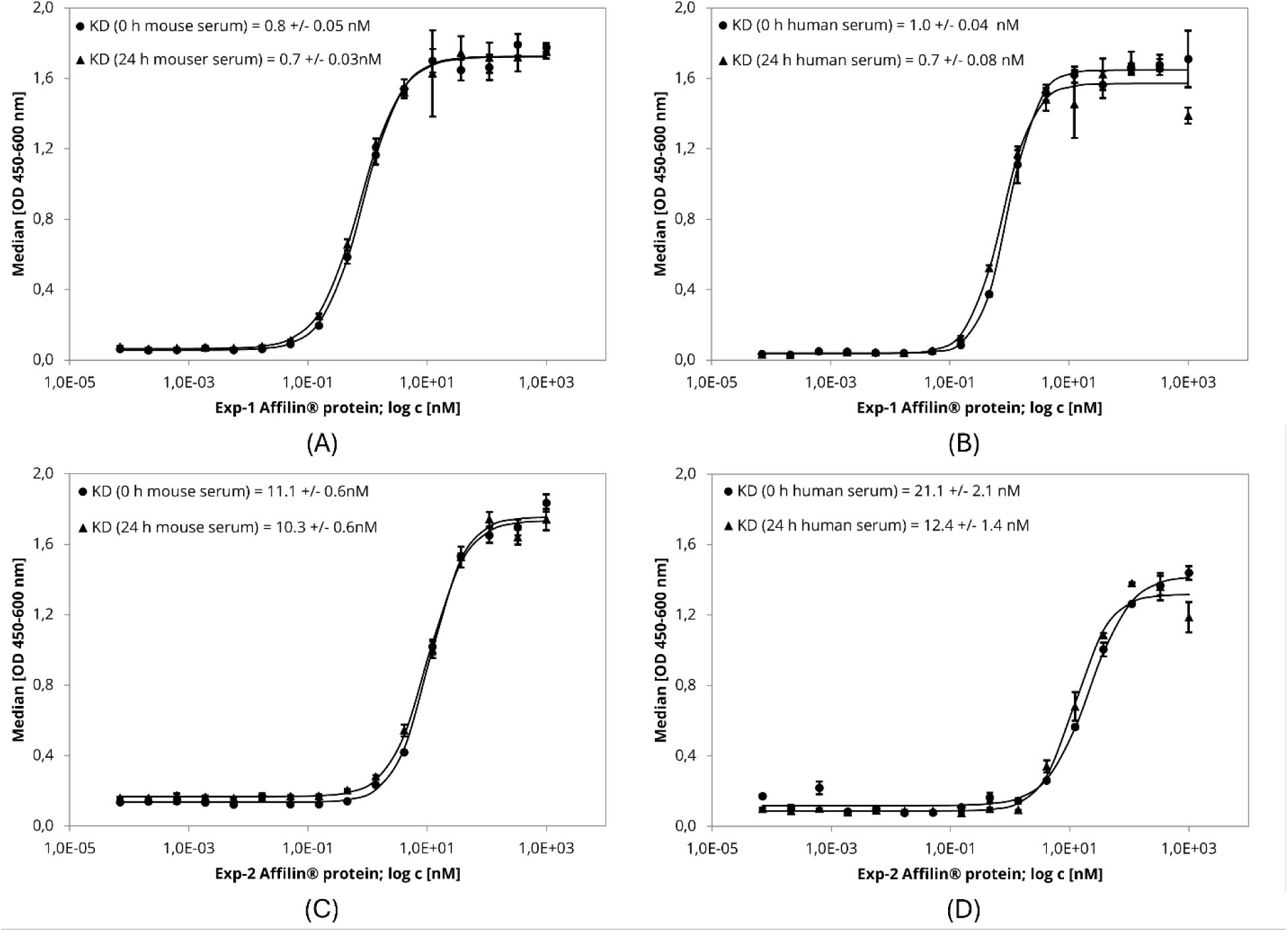
ELISA-based serum stability assay with lab scale purified Affilin^®^ proteins Exp-1 and Exp- 2. 384-well plates were immobilized with HER3-Fc as target protein. Affilin^®^ proteins were prepared in a concentration-dependent manner starting at 1 µM. Binding was detected using a 30x-bioinylated anti- ubiquitin antibody and a peroxidase-coupled Streptavidin. Readout was done using absorption at 450 nm and 600 nm. (A) Exp-1 after incubation in mouse serum for 0 h and 24 h @ 37 °C. (B) Exp-1 after incubation in human serum for 0 h and 24 h @ 37 °C. (C) Exp-2 after incubation in mouse serum for 0 h and 24 h @ 37 °C. (D) Exp-2 after incubation in human serum for 0 h and 24 h @ 37 °C.

The sequences of the *in vitro* identified Affilin^®^ proteins contain identical residues at some amino acid positions, but have also positions with different residues (Figure S10, Table S2).

### In silico maturation of HER3-binding Affilin^®^ proteins to improve performance using ProteinMPNN

After identifying 7 hits computationally and 4 experimentally, we aimed to boost the binding affinity of the best hits through site-directed mutagenesis, namely of the computationally designed Affilin^®^ protein with the best binding (Comp-9 with KD = 29.8 nM) and the top two experimental HER3 hits (Exp-1 with KD = 5.4 nM, and Exp-2 with KD = 20.1 nM). Binding the target in a natural context on cells is a key feature for therapeutic applications. To achieve this, a general improvement of affinity was the next step. We capitalized on recent advancements in AI-driven protein engineering, specifically exploring the capabilities of ProteinMPNN (29) to propose original plausible mutants. This deep learning algorithm addresses the inverse folding problem ー predicting protein sequences capable of adopting a specific fold. ProteinMPNN has proven effective in redesigning binder sequences, as exemplified by its success in enhancing affinity to the GRB2’s Src homology 3 (SH3) domain by testing approximately 100 mutants (29). Therefore, our objective was to integrate this tool with our MM pipeline (Figure SN1a) to strategically reduce the number of mutants required for testing to improve the binding affinity of an initial hit.

In this context, we first identified the interface residues that were not significantly contributing to the binding as potential positions to be engineered by ProteinMPNN. Those residues were identified as the ones with an energy contribution above -2 Kcal/mol when decomposing the binding energy of each HER3 binding Affilin protein with MM-GB(PB)SA into the residues on the protein-protein interface. Allowing mutagenesis in the identified residues, we generated 100 sequences per Affilin^®^ protein with ProteinMPNN. Each sequence pool was clustered at 97.5 % of sequence identity with MMseqs2 (30). The centroid of each cluster (10 clusters for Comp-9, 12 for Exp-1, and 8 for Exp-2) was selected to be modeled with our MM pipeline. Specifically, we computationally introduced the mutations to the Affilin^®^ proteins and ran an MD simulation from the best binding mode found by PELE for the original Affilin^®^ protein. Using the MD simulations, we measured the binding energy of each variant (see Methods) and compared them to the initial hit (i.e. a ΔΔG computed as ΔG_NewVariant_ - ΔG_OriginalHit_) to identify the variants that could potentially improve the original hit binding affinity.

None of the nine ProteinMPNN variants created from the computationally designed Comp-9 Affilin^®^ protein showed a negative ΔΔG (Table S3), indicating that the variants did not improve the binding affinity with respect to the original hits. On the other hand, the top 2 Affilin^®^ proteins from the experimental pipeline had 6 and 5 variants with better predicted binding energies (Table S3), which also showed good AlphaFold2’s pLDDT and RMSD values (Figures S11 and S12). Since the predictive power of this strategy is limited by the effects of multiple mutations, which are typically not properly modeled (31,32), we came up with a list of 27 single mutants (Table S4) based on the multiple mutants from ProteinMPNN and the available experimental information from other assayed Affilin^®^ hits (Table 1 and 2). Twelve of these single mutants showed negative ΔΔG (Table S4), but 3 of those variants showed slightly higher RMSD values along the MD simulation (Figure S13) compared to the original variant, which may be caused by a change in the binding mode. Lastly, we generated a list of double and triple mutants (hereafter referred to as “consensus mutants’’) that combined the changes of the ProteinMPNN mutants that contribute more to the improvement in the binding affinity according to the MM-GB(PB)SA calculations. In this case, we generated the AlphaFold2 models for the 7 consensus mutants and launched the MD simulations to estimate the binding energy, like in the previous mutants. These mutants showed confident AlphaFold2 structures (Figure S14), as well as good RMSD values along the MD (Figure S15). Four out of the seven assayed mutants were predicted to have a better binding affinity (a negative ΔΔG) (Table S5).

### In vitro maturation of HER3-binding Affilin^®^ proteins using phage display and HTS

In parallel to the *in silico* maturation, we performed an *in vitro* maturation selection using libraries based on the identified HER3-binding Affilin^®^ proteins. Specially designed maturation libraries based on Comp-9, Comp-11, Exp-1, Exp-2, Exp-3 and Exp-4, in total nine libraries, were applied in a phage display consisting of two selection rounds with strong reduction of target amount and including mouse serum incubation of phages before each round to eliminate binders with instability in serum. Many pools revealed a specific and increased binding to HER3 in the phage pool ELISA (example see Figure S16).

The phage pools were subcloned into an expression vector generating C-terminal StrepTag fused Affilin^®^ proteins that were identified using StrepTactin-HRP during an ELISA-based maturation screening. The screening setup included 360 clones per pool and either biotinylated HER3 or HER3-Fc as target protein. To evaluate the specificity for each variant Sigmablocker or hIgG1-Fc was applied as negative control. The thresholds for nominating hits were derived from the binding signals obtained from the parental Affilin^®^ proteins of the respective maturation libraries. In total, 564 out of 3240 tested clones were labeled as hit (example see Figure S17) and validated in a µ-scale format.

### Validation of optimized Affilin^®^ proteins

The most promising mutants from the *in silico* maturation underwent a first round of experimental validation, namely 17 mutants from ProteinMPNN, 9 single mutants, and 4 consensus mutants. The genes for the mutants were synthesized, expressed and the proteins purified as µ-scale samples and the experimental results showed that the 17 ProteinMPNN mutants could be expressed but lost their binding affinity to the HER3-Fc target although being computationally predicted as favorable. The disagreement between the computational and experimental results is probably due to the high number of mutations (between 8 to 16 mutations per mutant) introduced in each mutant, which significantly diminishes the computational predicting power (31,32). On the other hand, both the single and the consensus mutants gave more promising results, with all being expressed and 6 showing similar or even improved binding affinity in the sub-nanomolar range (Table S7 – refer to *in silico* round 1). The best two variants, Exp-MatComp-4 and Exp-MatComp-12, with improved binding affinity were selected for further analysis as lab scale samples by SPR and cell binding (Table S8 – refer to Exp-MatComp-4 and Exp-MatComp-12).

The µ-scale purified Affilin^®^ proteins from the *in vitro* maturation were evaluated by SPR measurement against HER3-Fc. The SPR data confirmed an improved affinity for 100 maturated Affilin^®^ proteins, which were evaluated in a cell-based assay. The selected variants showed good expression and solubility in *E. coli* BL21 (DE3) (Figure S18). The variants were tested on HER3-overexpressing HEK293 cells and native HER3-expressing SK-BR-3 cells. The cell-based assay revealed 8 out of the 100 variants with an improved affinity, which were nominated as hit (Table S6). Two of the best binders were the Exp-MatExp-5 and Exp- MatExp-8 Affilin^®^ (Figures S19 and S20).

The expression of the computational and experimentally optimized Affilin^®^ proteins in the lab scale format showed well behaved soluble proteins (Figure S18, Figure S21). The purification via StrepTag and the polishing via size exclusion chromatography produced mainly highly pure protein, however some variants showed partial dimerization or partial proteolysis. The SPR analysis confirmed an improved binding affinity for two of the computational and two of the experimentally optimized Affilin^®^ variants (Table 3, Table S7 – refer to *in silico* round 1, Figure 6). The affinity could be improved to even sub-nanomolar affinities. Binding to HER2-His could not be detected, which verifies the specificity of the binding.

**Figure 6.**
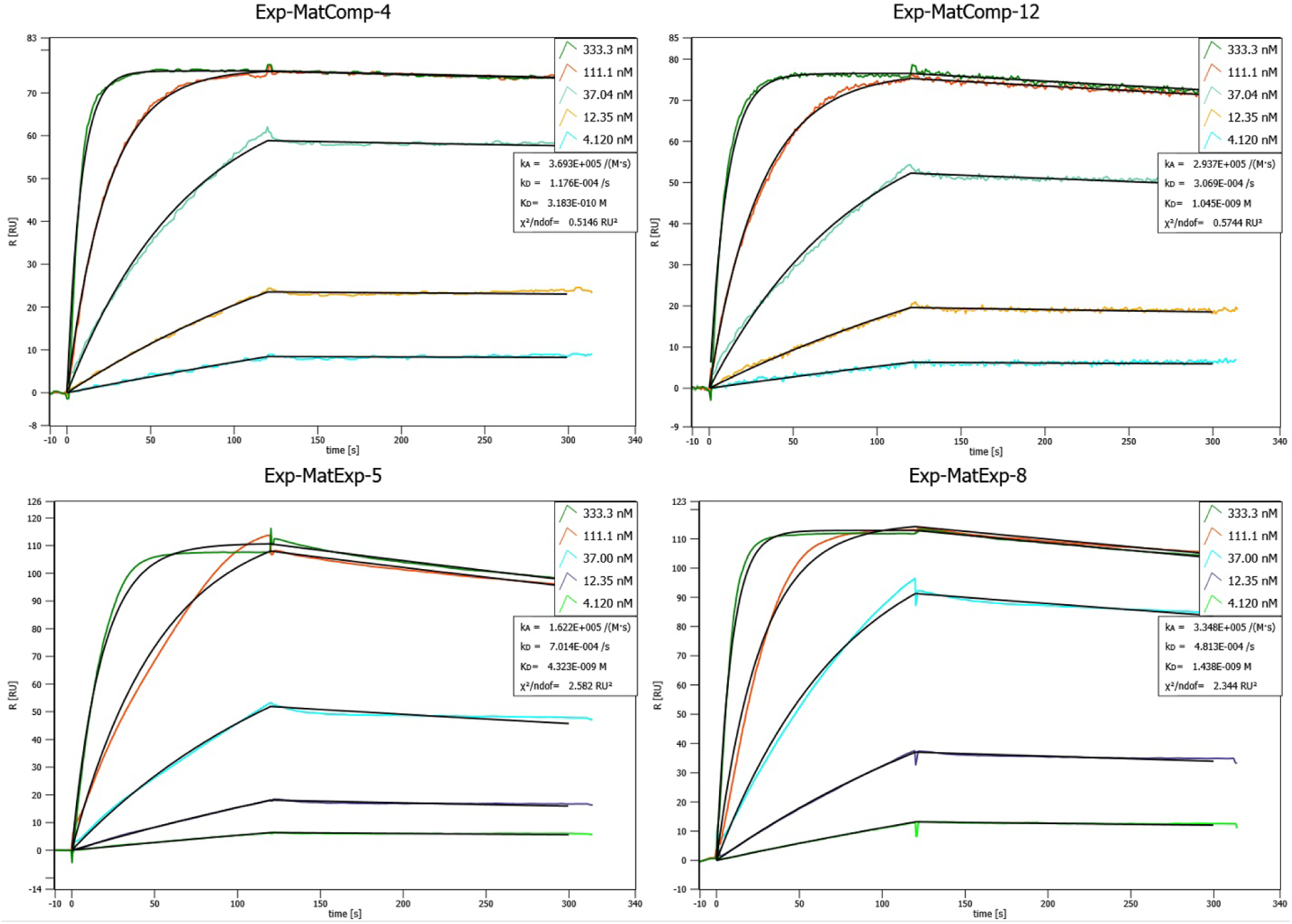
Concentration-dependent SPR binding curves of different matured, lab scale purified Affilin^®^ proteins. Mat-ExpComp variants were maturated computationally and show the best binding affinities against HER3- Fc. The Exp-MatExp variants were experimentally maturated and show very similar affinities towards HER3-Fc. SPR was performed on a Sierra SPR-32 (Bruker) with a Protein A-coated chip on which the HER3-Fc was immobilized.

**Table 3.** Summary of binding affinity towards HER3-Fc for in vitro selected Affilin^®^ proteins obtained with different methods. Summary of binding affinity data for *in vitro* and *in silico* maturated Affilin^®^ proteins, lab scale purified and analyzed with different methods. SPR measurement was performed on a Sierra SPR-32 (Bruker) with HER3-Fc immobilized on a Protein A-coated chip (Figure 6). Serum stability assay in mouse or human serum was performed by an ELISA with HER3-Fc immobilized on 384-well plates. Cell binding was measured on a Guava easyCyte 5HT FACS using HEK293-HER3-sc87 cells and Affilin^®^ proteins in a concentration dependent manner. Table abbreviations: nd – not determined

| Affilin® | SPR<br>(lab scale)<br>[nM] | Binding in mouse<br>serum [nM] |  | Binding in human<br>serum [nM] |  | Cell binding to<br>HEK293-hHER3-sc87<br>[nM] |
| --- | --- | --- | --- | --- | --- | --- |
|  |  | 0h | 24h | 0h | 24h |  |
| Exp-MatExp-5 | 4.2 | 0.8 | 0.8 | 0.8 | 0.9 | 0.2 |
| Exp-MatExp-8 | 1.5 | 1.1 | 1.1 | 1 | 0.9 | 0.03 |
| Exp-MatComp-4 | 0.2 | 0.7 | 0.6 | 0.7 | 1 | 0.6 |
| Exp-MatComp-12 | 1.1 | 0.8 | 0.7 | 1.3 | 0.9 | 0.6 |
| Exp-1 | 5.4 | 0.8 | 0.7 | 1 | 0.7 | nd |

Furthermore, the cell binding was evaluated. A strong binding to HER3-overexpressing HEK293 cells and native HER3-expressing SK-BR-3 cells could be seen using FACS (Table S8). The absolute binding signals on HER3-HEK293 cells were higher in comparison to SK-BR-3 cells. A concentration-dependent measurement with the Affilin^®^ variants on cells revealed affinities in the sub-nanomolar range, with Exp- MatExp-8 as the most affine variant (Table 3, Table S7 – refer to *in silico* round 1). Finally, the stability test in human and mouse serum showed no loss of affinity after 24 h incubation time for the maturated Affilin^®^ proteins (Table 3). The sequences of the most promising maturated Affilin^®^ proteins show mutations especially, at positions 6 and 7 (Figure S22, Table S2).

In view of the first *in silico* round results, we tried to further improve the binding affinity in a last round of single and double mutants by mutating with ProteinMPNN positions that were not assayed in the first round of mutagenesis. In this case, we computationally tested the new mutants using the initial and last MD frames of the best hit (Exp-1). With this analysis and a visual inspection of the complex interface, we came up with a list of 4 single and 1 double promising mutants (Table S9, Figures S23 and S24). We experimentally tested the 5 second-round promising mutants, as well as 2 mutants from the first round which could be promising according to the experimental results of the first round. All the mutants could be expressed in lab scale format (Figure S21) and 4 out of the 7 tested showed better binding affinity measured than the initial hit, in particular two had an improvement of 8-fold or higher and showed better cell binding than the original Affilin^®^ protein hit Exp-1 (Table S7 – refer to *in silico* round 2). However, the binding was not better than the above presented variants.

## DISCUSSION

Herein, we describe the design of high affinity ligands for the challenging RTK HER3 protein using a combined wet chemistry and AI + MM computational pipeline. Our approach enabled us to obtain these promising HER3 binders while optimizing laboratory resources and reducing the number of experimental assays needed, demonstrating a highly efficient workflow for binder discovery that can be adapted to other scaffolds and laboratory settings.

A set of primary binders was obtained by a classical phage display selection campaign reaching affinities up to the single-digit nanomolar range. Notably, two of these binders displayed cell binding and serum stability, which is mandatory for *in vivo* applications. By only using a small-sized dataset of 759 Affilin^®^ sequences against different targets, the *in silico* work yielded two *de novo* binders with a remarkable affinity against the biochemical target (KD < 100 nM) only requiring 12 experimental assays. This harnesses the power of combining VAE algorithms, with MM techniques to make the most out of the available data and strategically prioritize molecules for experimental validation. However, unlike the experimentally derived binders, the computationally designed binders did not demonstrate cell binding. This limitation highlights two key challenges. On the one hand, it points out the inherent difficulties of purely structural computational methods to fully capture the complexity of serum interactions and cellular environments (33). On the other hand, generating *de novo* sequences with cell binding properties might be a different challenge altogether. A primary obstacle is the lack of relevant sequence data annotated with and spanning all possible environmental and contextual in-cell parameters. This limitation underscores that, to the best of our knowledge, no state-of-the-art model has proven capable of embedding such intricate contextual knowledge from solely sequence data. Nevertheless, the flexibility of the AI+MM approach broadens the accessibility of achieving *in vitro* successful outcomes with limited data, thereby empowering a wider array of laboratories to participate and succeed in binder design endeavours.

Starting from the hit sequences obtained via phage display it could be demonstrated that the MM pipeline can also be used to suggest and prioritize mutants (having a higher success rate for single mutants) for enhancement of binding affinity from Affilin^®^ proteins, an improvement of up to 16-fold with respect to the parental variant could be achieved. A maturation selection of the phage display hits yielded optimized binders in that range as well, underscoring the value of computationally guided mutant prioritization. Interestingly the same key position for the interaction, K6 was identified in both approaches. The combined use of wet chemistry and computational methods revealed the best outcome in the endeavour to identify HER3 ligands with high affinity, cell binding and serum stability for theranostic applications, including imaging and treatment of multiple cancers due to HER3’s implication.

Ongoing developments and new routes using these methods will pave the way for even better outcomes in ligand identification and optimization. A complex design integrating specifications like the binding of selected epitopes, species cross reactivity and isoform specificity while preserving compound stability and manufacturability requires the application of complex tools.

### Limitations of the study

Despite the promising results of our combined wet chemistry and AI + MM computational pipeline, several limitations warrant consideration. First, while the computational approach successfully identified *de novo* binders with high biochemical affinity, these binders did not exhibit cell binding, which is a critical factor for their potential therapeutic application. This discrepancy underscores a key limitation of purely structural computational methods (33), and the lack of sequence data annotated with environmental and contextual in-cell parameters to generate embeddings that can account for the cellular context, as mentioned in the Discussion.

Another limitation stems from the relatively small dataset used in the *in silico* design phase (759 Affilin^®^ sequences). Although this approach demonstrated success with this dataset, the generalizability of the results to larger or more diverse datasets remains uncertain. The ability of the VAE algorithms to extract meaningful insights from limited data is a notable strength, but the performance of this method may vary when applied to other scaffolds or targets that are more structurally complex or poorly characterized.

In the context of *in silico* affinity maturation, it was crucial to introduce only a small number of mutations to the Affilin^®^ hits to use our MM pipeline to effectively identify mutants with improved binding affinity. When many mutations are introduced, MM tools often struggle to accurately predict the resulting behavior. In such cases, the MM predictions can fail, potentially leading to the loss of binding to the target protein, as discussed in the Results section.

The *in vitro* approach is limited by the number of variants, which can be presented and identified by the selection and screening process. An Affilin^®^ library may compromise a theoretical diversity of up to 10^17^ variants, depending on the number of randomized positions and permitted amino acids. However, the preparation of the phage library, including transformation and culturing of *E. coli* leads to a representation of just ∼10^10^ variants per library. Target specific phage pools after the selection may still contain up to 10^5^ different clones from which the high-throughput screening is able to cover ∼15.000 clones per screen. The consequences of this limitation are compensated by the enrichment of potential binding variants during the selection process and can therefore be covered by the high-throughput screening.

## RESOURCE AVAILABILITY

### Materials availability

*De novo* Affilin^®^ sequences generated in this study have been deposited to the supplementary information (Document S2).

### Data and code availability

All original code is deposited in our GitHub repository: https://github.com/annadiarov/ProtVAE under MIT license.

## ACKNOWLEDGMENTS

We thank Dr. Ulrich Haupts for compiling the Affilin^®^ training data set and Jaydeep Belapure for critical review of the final draft. We also thank the Spanish Ministry of Science and Innovation for funding A.M.D-R. predoctoral fellowship (FPU/03921).

## AUTHOR CONTRIBUTIONS

Conceptualization, A.M., L.D., V.G., E.B.-D. and S.R.; methodology, A.M.D.-R., A.M., M.G.-B., M.M., J.L., G.H. and S.R.; software, A.M.D.-R., A.M. and S.R.; validation, G.H. and J.L.; formal analysis, A.M.D-R., C.P., G.H., I.C., J.L. and S.R.; investigation, A.M.D.-R., M.Z., M.G.-B., I.C., G.H., C.P., A.M., J.L. and S.R.; resources, M.M., A.M., S.R.; data curation, A.M.D.-R., C.P. and S.R.; writing—original draft preparation, A.M.D.-R., E.B.-D., M.Z., G.H., J.L. and S.R.; writing—review and editing, A.M.D.-R., V.G., E.B.-D. and S.R.; visualization, A.M.D.-R., G.H., M.G.-B., J.L. and S.R.; supervision, V.G., E.B.-D. and S.R.; project administration, M.Z., E.B.-D., and S.R.

## SUPPLEMENTAL INFORMATION

Document S1. Figures S1–S24, Table S1-S9, and Supplementary Note 1 (with Figures SN1-SN13 and Table SN1) describing the benchmark for the Molecular Modeling pipeline.

Document S2. List of *de novo* generated sequences with the metrics obtained from the AI+MM pipeline.

## MATERIALS & METHODS

### Available Affilin® protein data

We disposed of a total of 759 Affilin^®^ protein sequences, 329 of them monomeric, with experimental KD against 14 targets (Table 4), and one crystal (PDB: 8PEQ) between a strong dimeric Affilin^®^ binder towards the fibronectin (FINC) extra-domain B (ED-B). To establish the modeling pipeline (see Supplementary Note 1), we concentrated on the monomeric proteins to prevent inaccurate results due to the uncertainties in dimerization. However, we trained the ML model with all available data since discarding any data would be detrimental in this small dataset. Namely, for each dimeric sequence, we obtained two monomeric sequences cutting by the Affilin^®^ N-terminal sequence motifs (MQI, MRI, MAS, MTI). This led to 852 non- identical (at 99% identity) monomeric Affilin^®^ protein sequences.

**Table 4.** Summary of Affilin^®^ protein sequences available used for the training set. Number of monomeric Affilin^®^ protein sequences and the number of weak binders (WB; KD ≥ 100 nM), medium ones (MB; 100 nM < KD ≤ 10 nM), and strong ones (SB; KD < 10 nM) by target.

| Affilin® | SB | MB | WB |
| --- | --- | --- | --- |
| HER2 | 72 | 59 | 16 |
| FINC ED-B | 36 | 32 | 18 |
| Others | 127 | 208 | 191 |

### Protein structure prediction and system preparation

The initial structure of the target proteins, FINC and HER2 for the retrospective study and HER3 for the prospective one, were obtained from the PDB database (FINC: 8PEQ, HER2: 7MN6, HER3: 7MN6). The Affilin^®^ proteins were generated with AlphaFold2(34) (version 2.1.0), but we also used MODELLER(35,36) in the FINC retrospective study since we had as reference the crystal of the Affilin^®^ protein in complex with FINC (PDB: 8PEQ) (see Supplementary Note 1). All structures were prepared using the Protein Preparation Wizard (PPW) tool (37) from Schrödinger. More in detail, PPW was employed to add missing hydrogen atoms, reconstruct incomplete side chains and short missing loops(38), assign the ionization states at pH 7 and optimize the hydrogen bond network. Finally, we performed a restrained minimization allowing hydrogen atoms to be freely minimized using the OPLS2005 force field(39).

In the FINC retrospective study, we also computed each monomer’s alpha carbon root mean square distance (Cɑ-RMSD) against the crystalized Affilin^®^ protein. To compute the Cɑ-RMSD between different proteins, we used the Biopython library(40) (version 1.80) to align the sequences (with gap_penalty -10.0, extend_penalty -0.5, max_gap 50, substitution matrix BLOSUM62) and compute the Cɑ-RMSD of the sequence pairs.

### Rigid Protein-Protein Docking

Rigid PPDs were performed with PyDock(22). For each docking pair, we generated 92.400 poses using FTDock sampling(41), keeping the target protein fixed (i.e., receptor protein) while rotating and translating the Affilin^®^ protein (i.e., ligand-protein). The docking parameters were set to default.

In the retrospective analysis of the FINC study (see Supplementary Note 1), we determined the ligand- protein root mean square distance (L-RMSD) compared to the Affilin^®^-FINC crystal. This metric is calculated as the Cɑ-RMSD of the ligand-protein (i.e., Affilin^®^ protein) upon aligning it with the receptor protein (i.e., the target protein).

### Protein-protein interface filtering

To predict the interacting regions of both the target protein and the Affilin^®^ proteins, we used both the Optimal Docking Area (ODA)(42) from PyDock and the MaSIF-site(10) using default parameters. The predicted protein-protein interfaces were used to filter all the poses obtained from the PPD to ensure they were close enough to interact. Namely, we kept poses where at least one residue (using Cɑ atoms) in the interacting patch of one protein is within 10 Å of any other residue in the partner protein’s interacting patch. For HER2 and HER3 systems, we applied an extra filter to discard the poses in which the Affilin^®^ protein was interacting with the HER2-HER3 dimerization area.

### Clustering filtered poses

We employed an iterative clustering method designed to consider both pose diversity and energetic relevance. Namely, for each system, we kept the best energy pose, and we then compared it with the second-best one. If the L-RMSD between them was greater than 2 Å, we also kept the second too; otherwise, we discarded it for being too similar to a previously seen structure with better energy. We repeated this process iteratively until we obtained 50 poses—a computationally manageable quantity of poses for performing more resource-intensive calculations like PELE. Through this approach, we ensured pose diversity and the most energetically representative poses.

### PELE refinement simulations

PELE was used to assess the predicted binding mode(s) from PyDock and estimate the binding affinity of the Affilin^®^ protein to the target (20). PELE is a heuristic MC-based algorithm coupled with protein structure prediction methods. The software begins by sampling the different microstates of the ligand through small rotations and translations. Applying normal modes through the anisotropic network model (ANM) approach (43), the protein’s flexibility is also considered. Once the whole system has been perturbed, side chains of the residues at the protein-protein interface are sampled to avoid steric clashes. Last, a truncated Newton minimization with the OPLS2005 force field is performed (44), and the new microstate is accepted or rejected based on the Metropolis criterion. The Variable Dielectric Generalized Born Non-Polar (VDGBNP) implicit solvent model (45) was used to mimic the effect of water molecules around the protein. Each PELE simulation per system utilized 200 computing cores for 2h.

### Molecular Dynamics

Classical MD simulations with Amber (46) were performed to estimate the binding affinity of the Affilin^®^ proteins towards the studied targets (the simulation times change depending on the system, they are specified along the results). MD simulations were launched for each system using the best pose obtained in the PELE refinement. A water cubic box (distance of 12 Å between the closest protein atom and the edge of the box) was created around the system using the TIP3P water model. The charge of the system was stabilized using monovalent ions (Na^+^ and Cl^−^), and the optimal number of counter ions (Na^+^ and Cl^−^) was added to reach a salt concentration of 0.1 M. The protein system was parameterized with the AMBERff14SB (47) force field. Berendsen thermostat and barostats were applied for the NPT ensemble (constant pressure and temperature, 1 bar and 300 K, respectively). The Verlet integrator with a 2 fs time step was used with a 12 Å nonbonded long-range interactions cutoff. MD simulations were analyzed calculating the backbone L- RMSD of the Affilin^®^ proteins along the simulation.

### Binding affinity estimation and identification of key interaction residues

MMPBSA.py script (48) was applied to some long MD simulations to compute the binding energy calculations and per-residue energy decomposition of the protein complexes. Using 500 snapshots extracted from the last 500 ns of each MD trajectory, we applied the MM-GB(PB)SA (23) method to calculate the binding energy (ΔG). Conformational entropy contribution to binding was not included, given the difficulty of computing it for large systems like protein-protein complexes, and its small effect when calculating relative (and similar) system free energies. Computed ΔGs were used to determine the Affilin^®^ variants that improved the binding by calculating the relative binding energy (ΔΔG) between variants. On the other hand, we used the per-residue binding energy (ΔGres) to identify hotspots, that is, residues that highly contribute to the binding (ΔGres <-2 Kcal/mol) (49). The energetic contribution of each single residue in the interface was calculated by adding its interactions over all the residues in the system.

### Variational Autoencoder

To model the latent space of our data and enable the generation of novel Affilin^®^ proteins, we employed a VAE architecture (27). The VAE was fed with a 164 vector containing the sequence one-hot-encoded using 20 amino acids plus a padding token for shorter sequences. The encoder comprised 2 neural network layers. An LSTM layer with 256 neurons followed by a fully connected layer of 128 neurons that transforms the input data into the latent space. The decoder network consisted of 3 layers. The decoder was built with two layers to decode hidden and current states with 256 neurons each one followed by a 256 neuron LSTM layer that progressively transforms the representations of the latent vector back into the input sequence. The entire VAE was trained end-to-end by optimizing the evidence lower bound (ELBO) loss function which includes a reconstruction term and a regularization term, encouraging the learned latent space to follow a standard normal distribution (27).

The model was implemented in PyTorch (50) and trained using 852 monomeric Affilin^®^ sequences for 42 epochs with an early stopping criterion upon convergence of the validation loss for 5 epochs. The Adam optimizer was used with a learning rate of 0.001. The architecture of the VAE, including the number of layers and neurons in each layer, was determined through a process of experimentation to achieve a balance between model complexity and performance on the task at hand.

New Affilin^®^ sequences were generated using the optimized decoder network. While these sequences were influenced by the patterns of the initial data, the VAE’s sampling approach facilitated the creation of novel protein sequences. Specifically, to ensure both broad exploration and biological plausibility, our method autoregressively decodes random latent vectors drawn from a standard normal distribution (N(0,1)) to generate novel protein sequences. Controlled variability introduced via random latent vectors allows for the discovery of unique sequences. To balance novelty with accuracy, we utilize temperature-controlled sampling, modulating the stochasticity of sequence generation—higher temperatures increase diversity, while lower temperatures yield more predictable outcomes.

### Materials for experimental assays

HER3 extracellular domain containing His-Avi-tag was purchased from AcroBiosystems (Cat. No. ER3- H82E6; Ser20-Thr643). When HER3-Fc or HER3-Avi-His-tag is mentioned in the text, it refers to the extracellular domain of the protein, with the respective tag.

HER2/ErbB2 extracellular domain containing His-Avi-tag was purchased from AcroBiosystems (Cat. No. HE2-H82E2). HER2-His was purchased from R&D Systems (Cat. No. 10126-ER). When HER2-Avi-His-tag or HER2-His is mentioned in the text, it refers to the extracellular domain of the protein, with the respective tag.

Cell culture: Native HER3-expressing SK-BR-3-cells were cultured in McCoy’s 5A -medium with 10 % FCS and HEK293 cells were cultured in DMEM with 10 % FCS at 37 °C, 5 % CO2 and 90 % humidity. To establish overexpressing cell lines, plasmid-DNA of ErbB 3 (ERBB3) (NM_001982) Human Tagged ORF Clone (Origene; RC209954) was transfected into HEK293-cells with FuGENE^®^ HD Reagenz (Promega; E231A). Single-cell clones were selected with 1 mg/ml G418 and analysed with Patritumab for HER3- expression. The control cell line was established with pCMV6-Entry Mammalian Expression Vector (Origene; PS100001).

### Target protein production

Expression: For target expression, Expi293-cells were cultured in Expi293™ Expression Medium (Gibco; 13489756) at 37 °C, 8 % CO2 and 95 % humidity and the extracellular domain of HER3 (ECD-HER3; Ser20- Thr643) in fusion with human Fc-domain and the leader sequence of EGFR were cloned in the mammalian expression vector pCEP4. Expi293-cells were transfected with pCEP4-ECD-HER3-Fc according to the manufacturer’s instructions.

Purification of HER3-Fc: Cell culture supernatant containing HER3-Fc was purified via affinity chromatography using a HiTrap Protein A column (Cytiva). Protein was eluted with 100 mM citrate pH 4.0. After adjusting pH to 7.3 and adding 150 mM NaCl, the protein was further purified by a Superdex™ 200 HiLoad 16/600 gel filtration column equilibrated with PBS, pH 7.3. Protein-containing fractions were analyzed by SDS-PAGE and protein concentration was determined via UV280 nm absorption.

### Primary selection by phage display

The selection via phage display was performed with nine different phage libraries based on ubiquitin scaffold. The highly complex libraries contain more than 10^10^ Affilin^®^ proteins, which are cloned into a pCD12 phagemid, a derivative of pCD87SA (51). Three selection rounds were performed with decreasing amounts of target protein starting from 90 pmol to 4 pmol. Each selection round started with pre-incubation of the phage libraries with Sigmablocker-blocked (Sigma-Aldrich; cat. no. B6429-500ML) M-270 Streptavidin Dynabeads (Thermo Fisher; cat. no. 65306) for 1h. Sigmablocker is further designated as an OFF target. Parallel to the pre-selection a biotinylated human HER3-His-Avi construct (AcroBiosystems; cat. no. ER3- H82E6), was immobilized to M-270 Streptavidin Dynabeads. The biotinylated human HER3-His-Avi is further designated as an ON target. After removing the beads from the pre-selected phage-pool, the ON target containing beads were added and incubated for 1.5h with the phages. The whole selection process was performed by using the automated KingFisher Duo system (Thermo Scientific), which contains magnetic tips to focus the beads in the samples. Each selection round was finished by an increasing number of washing steps with 1xPBS-containing buffer and elution of the bound phages to the bead-immobilized target by adding trypsin for 30 min at 37 °C to the samples. The eluted phages were then used for a re- infection into *E. coli* ER2738 and cultured overnight. Before the next selection round started, the phages were prepped out of the *E. coli* cultures using 20 % PEG-6000 sodium chloride solution and already-known protocols. A pre-incubation of the prepped phages was performed with mouse serum for 21h at 37 °C before rounds two and three were started.

The selection was evaluated by performing a phage-pool-ELISA by using a reserved sample of prepped phages from each selection round. The phage samples were normalized to 10^12^ phages /ml. The samples were applied to ON and OFF targets, which were indirectly immobilized to Streptavidin-coated 384-well plates. The number of phages was normalized to allow a comparison between the pools. The ELISA was run by standard protocol by using an anti-M13-HRP (B62-FE2) antibody for detection.

Specific binding pools were subcloned first into pNP-013 vector generating N-terminal GFP-fused Affilin^®^ proteins and secondly into pNP-002 vector generating C-terminal Strep-tagged Affilin^®^ proteins.

### High-throughput screening of primary selection pools

The high-throughput screening is based on an ELISA setup. The workflow contains a pre-screen with a small number of variants to find the best setup, a primary screen which uses only to ON target, and a secondary screen which consists of the ON and OFF targets and a pre-incubation in mouse serum.

The selected variants were provided by transforming the subcloned pools into electro- or chemically competent *E. coli* BL21 (DE3). The *E. coli* cultures were plated onto Q-trays with LB agar containing Kanamycin to obtain single clones. These single clones were automatically picked using the QPix system and cultured in 384-deepwell plates. After overnight cultivation in a 2xYT medium containing Kanamycin, the cells were lysated and the supernatant of the lysate was used as a sample in the screening process. To identify serum stable variants, the lysis step during the secondary screen was performed directly in mouse serum containing lysis additives for 24 h at 37 °C.

The setup for the pre-screen consisted of indirectly immobilized biot. HER3-His-Avi via Streptavidin-coated 384-well plates (ON target) or just Streptavidin-coated 384-well plates blocked with Casein (OFF target). The GFP-fused variants were used as samples, with 90 variants per pool. Positive binding variants were detected via GFP fluorescence. Each pool contained binders, so the primary screen was followed by using this setup.

The primary screen was performed with 14245 clones and only the ON target. Positive binding variants detected via GFP fluorescence were nominated for the secondary screen.

The secondary screen was set up with an indirectly immobilized ON target on Streptavidin-coated 384-well plates and additionally with indirect immobilized biotinylated human HER2 (AcroBiosystems; Cat. No. HE2- H82E2). GFP fluorescence was used for detection. The samples, nominated variants from the primary screen, were lysated in mouse serum. HER3 binding variants were nominated for µ-scale analysis.

The analysis in µ-scale format was done by small-scale chromatographic purification of *E. coli* lysate via StrepTactin-containing matrices in PhyNexus columns (Custom PhyTip by Biotage with resin from IBA). The samples were normalized to a concentration of 500 nM and used to measure binding kinetics against human HER3-Fc target protein immobilized to Protein A-coated SPR-chips on a Bruker-SPR32.

A specific number of nominated variants was subcloned into the pNP-002 vector generating C-terminal Strep-tagged Affilin^®^ proteins and used for additional µ-scale analysis on the Bruker-SPR32. The purified variants were measured in a concentration-dependent manner.

### Construction of maturation libraries

Hits from the primary selection and two *de novo* designed variants were used to generate maturation libraries. One maturation strategy was module shuffling where the paratope in the Affilin^®^ protein contributing to binding to HER3 was partly rerandomized. Another maturation technique used was epPCR, which randomly inserts exchanges into the cDNA. The cloning of maturation libraries and preparation of phage libraries was performed as described in (8). The maturation was either done on single variants or a variant pool.

### Maturation selection by phage display

The maturation libraries were applied to the phage display like in the primary selection. Biotinylated HER3- His-Avi was again used as the ON target and Sigmablocker-blocked beads were used as the OFF target. The target immobilization of M270-Streptavidin Dynabeads was done in the same way. The maturation selected consisted of only two rounds with a strongly reduced amount of target – in the first round 40 pmol and in the second round 0.5 pmol. The prepped phages were incubated in mouse serum before each round for 21 h at 37 °C.

The evaluation of the selection was done by phage-pool-ELISA identically to the primary selection but with a lower number of phages. Specific binding phage pools were subcloned into the pNP-002 vector to generate C-terminal Strep-tagged Affilin^®^ proteins.

### High-throughput screening of maturated pools

The screen was performed with 360 clones per pool. Subcloned pools were transformed into *E. coli* BL21 (DE3), plated onto Q-Trays, single clones were picked and cultured in 384-well plates. The expression cultures were lysated and the supernatant was used as a sample for the screen, no incubation in mouse serum was performed. The first ELISA setup contained biotinylated HER3-His-Avi on Streptavidin-coated 384-well plates as ON target or Casein-blocked Streptavidin-coated plates as an OFF target. The second ELISA used HER3-Fc on Protein A-coated 384-well plates or hIgG1-Fc on Protein A-coated 384-well plates as OFF target. The nomination of specific binding clones (hits) was done by setting a threshold above the binding signal of the parent variants, which were used to generate the maturation libraries. The nominated hits were analyzed in µ-scale format by SPR measurement, in the same way as earlier described, but with a sample concentration of just 100 nM.

### Cloning of Affilin^®^ protein constructs

Genes for Affilin^®^ proteins Exp-MatComp-5 - Exp-MatComp-21 were generated by PCR amplification with appropriate oligo nucleotides (see **Supplementary Information Tables S10 & S11**) introducing single amino acid mutations. PCR products were ligated via Type II restriction sites into pNP-002 (= pET28aS derivative), providing variants with a C-terminal StrepTagII for further purification steps.

Genes for Affilin^®^ proteins Comp-1 - Comp-12 & Exp-MatComp-1 - Exp-MatComp-4 containing more than one amino acid mutation were synthesized as string fragments (GeneArt, Regensburg, Germany) and cloned into pNP-002 as described above. Sequence identities were confirmed by external sequencing services (Microsynth AG, Balgach, Switzerland).

### Expression and purification of Affilin® proteins

Affilin^®^ proteins were expressed in *Escherichia coli* BL21 (DE3). Cells were grown in ZYM-5052 medium at 37 °C and harvested after 5 h. Cells were resuspended in 100 mM Tris, 150 mM NaCl, 1 mM EDTA, pH 8.0 and lysed by sonification. After centrifugation 20 µg Avidin per ml was added to the clear cell lysate and incubated for 30 min at room temperature. The solution was applied to a StrepTactin affinity column (GE Healthcare) and proteins were eluted with 100 mM Tris, 150 mM NaCl, 1 mM EDTA, 2.5 mM Desthiobiotin, pH 8.0. The eluted proteins were applied to a size exclusion chromatography column (Superdex™ 75 HiLoad 16/600) equilibrated with PBS pH 7.3. Protein-containing fractions were analyzed by SDS-PAGE and protein concentration was determined via UV280 nm absorption.

### Protein purity and apparent size

The apparent size for the proteins was determined by analytical size exclusion employing an UltiMate 3000 HPLC (Thermo Fisher Scientific) with the Superdex^TM^ 75 increase 5/150 GL_SE. The proteins were eluted using 1x PBS pH 7.3 with a flow rate of 0.3 ml/min at room temperature. For the approximation of the size, a linear calibration curve with the Gel Filtration Standard (BioRad) was used.

Purity was determined using a Vanquish Core HPLC (Thermo Fisher Scientific) with a PLRP-S 300A 5µm 250x4.6 mm. For elution a gradient form from 30% acetonitrile with 0.1 % TFA to 50% acetonitrile with 0.1% TFA at 55 °C with 1.1 ml/min flowrate over 40 ml was used.

### SPR

For SPR analysis the Sierra SPR^®^ 32-Pro (Bruker) with corresponding high-capacity amine sensors (Bruker) was used. HER3-Fc was used and bound on chips with immobilized Protein A (Sino Biological). The Fc-fusions were bound with around 500 RUs on the chips. For HER2-His-Avi (R&D Systems; Cat. No. 10126-ER) the protein was directly immobilized on Amine Sensors with 1000 RUs. The coupling reactions were performed according to the manufacturer’s protocol. For the determination of binding curves, the proteins were applied in concentrations ranging from 1000 nM to 0.46 nM in 1x PBS pH7.3 + 0.05 % Tween-20. The contact time of the proteins was 120s with 180s dissociation time and a flow rate of 30. For the calculation of the affinities, a 1:1 Langmuir fit was assumed.

### Melting point determination

Temperature stability was determined using DSF (differential scanning fluorimetry). Using the ViiA7 in combination with the Protein Thermal Shift Software 1.4 (Thermo Fisher Scientific), the protein solutions were analyzed in 1x PBS pH 7.3 with SYPRO^TM^ Orange (Sigma Aldrich) in a 1:200 dilution. The solutions were heated from 25 °C-95 °C with a ramp of 1 °C/min. From the resulting melting curves, the Boltzmann melting point was determined and given as the melting point of the protein.

### Flow cytometry analysis

Transfected HEK293-hHER3-cells, empty vector control HEK293-pEntry-cells, SKBR3-cells and HEK293- cells were trypsinized and resuspended in a medium containing FCS and washed in pre-cooled FACS blocking buffer (3% FCS; 0.1 % NaAc; PBS) and filled with 1x 10^6^ cells/ml into a 96 well plate. Cells were treated with a dilution series of StrepTag-fused Affilin^®^ proteins or 1 µg/ml Patritumab (Proteogenix) as positive control for 45 min at 4 °C. Affilin^®^ proteins were detected via rabbit anti-StrepTag antibody (GenScript; A00626), 1:300 diluted in FACS blocking buffer and 2 µg/ml of goat anti-rabbit-IgG- AlexaFluor488 antibody (Thermo Fisher Scientific; A11008). Patritumab was analyzed with 1 µg/ml of anti-human-IgG-AlexaFlour488 (Thermo Fisher Scientific; A-11013). Flow cytometry measurement was conducted on the Guava easyCyte 5HT device (Merck-Millipore) at excitation wavelength 488 nm and emission wavelength filter 525/30 nm.

### ELISA

Dilution series of Affilin^®^ proteins were incubated in 100 % mouse serum or 100 % human serum for 24 h at 37 °C. 384-well- plates (high binding; Greiner) were coated with 2.5 µg/ml HER3-Fc over-night at 4 °C, washed 3 times with 1xPBST (1xPBS, 0.1 % Tween) and blocked with 3 % BSA/ 0.5 % Tween/ PBS for 2 h at RT. Affilin^®^ proteins diluted in serum were incubated on ELISA plates for 1 h at RT. Binding to HER3-Fc was proven with biotinylated anti-Ubiquitin-antibody with a dilution of 1: 300 and Streptavidin-HRP with a dilution of 1: 5000 in 1xPBST.

