## Supplementary Information for "Efficient Design of Affilin^®^ Protein Binders for HER3"

### Part I

### Supplementary Information Figures

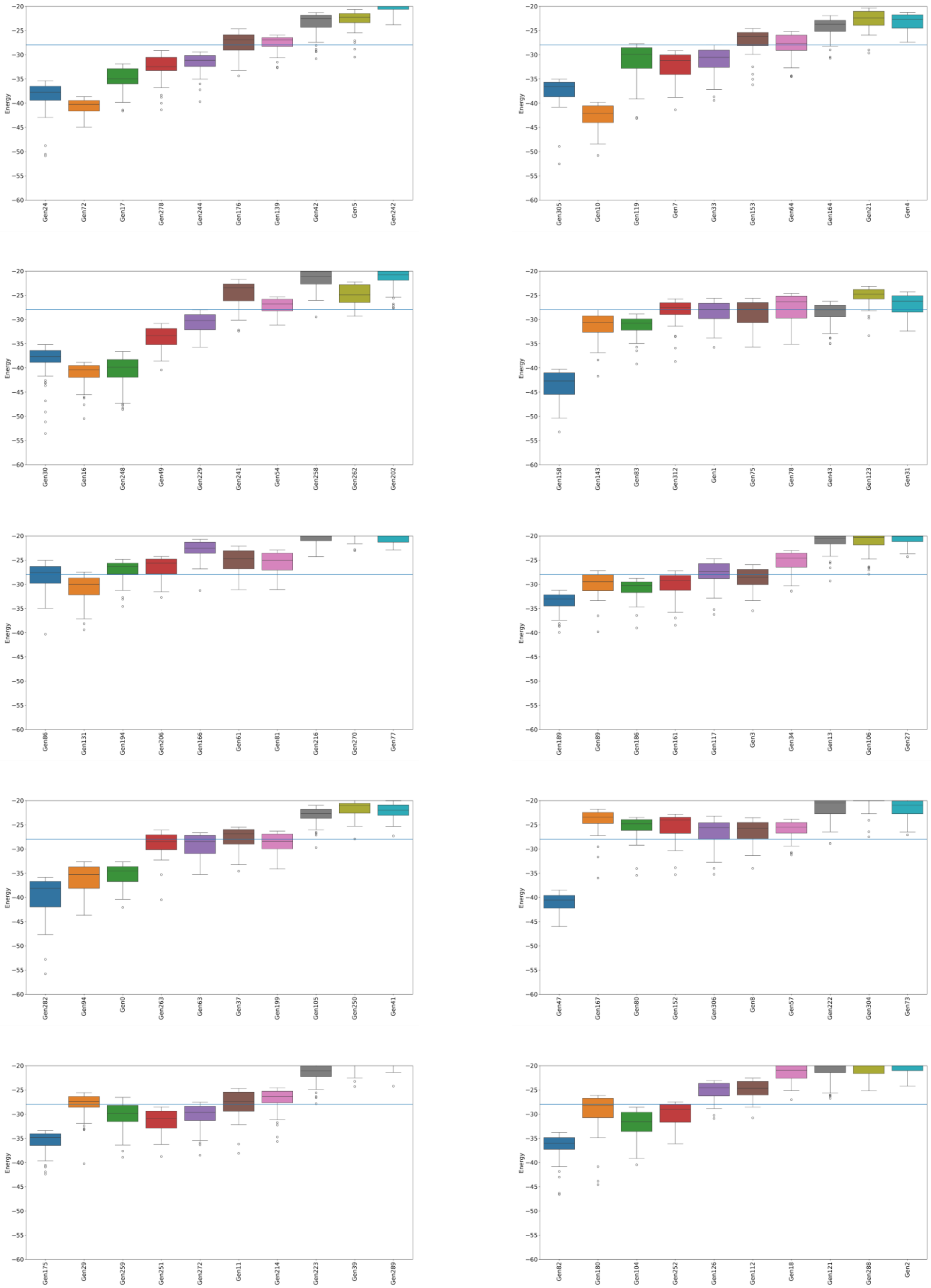
Energy
Gen24 Gen72 Gen17 Gen278 Gen244 Gen176 Gen139 Gen42 Gen5 Gen242
Gen305 Gen10 Gen119 Gen7 Gen33 Gen153 Gen64 Gen164 Gen21 Gen4
Gen30 Gen16 Gen248 Gen49 Gen229 Gen241 Gen54 Gen258 Gen262 Gen202
Gen158 Gen143 Gen83 Gen312 Gen1 Gen75 Gen78 Gen43 Gen123 Gen31
Gen86 Gen131 Gen194 Gen206 Gen166 Gen61 Gen81 Gen216 Gen270 Gen77
Gen189 Gen89 Gen186 Gen161 Gen117 Gen3 Gen34 Gen13 Gen106 Gen27
Gen282 Gen94 Gen0 Gen263 Gen63 Gen37 Gen199 Gen105 Gen250 Gen41
Gen47 Gen167 Gen80 Gen152 Gen306 Gen8 Gen57 Gen222 Gen304 Gen73
Gen175 Gen29 Gen259 Gen251 Gen272 Gen11 Gen214 Gen223 Gen39 Gen289
Gen82 Gen180 Gen104 Gen252 Gen126 Gen112 Gen18 Gen121 Gen288 Gen2

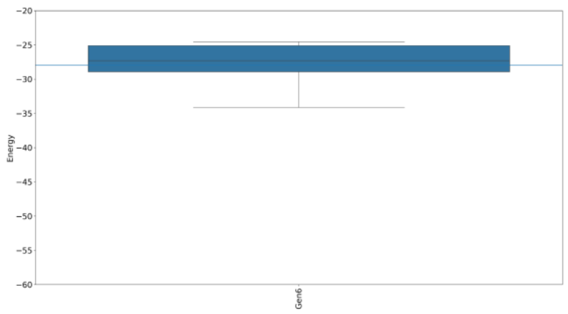


**Figure S1.** PyDock binding energy distribution of the top 50 different PPD poses (L-RMSD > 2Å) for the 102 selected Affilin® sequences targeting HER3. The blue line indicates the mean binding energy value among all the systems.

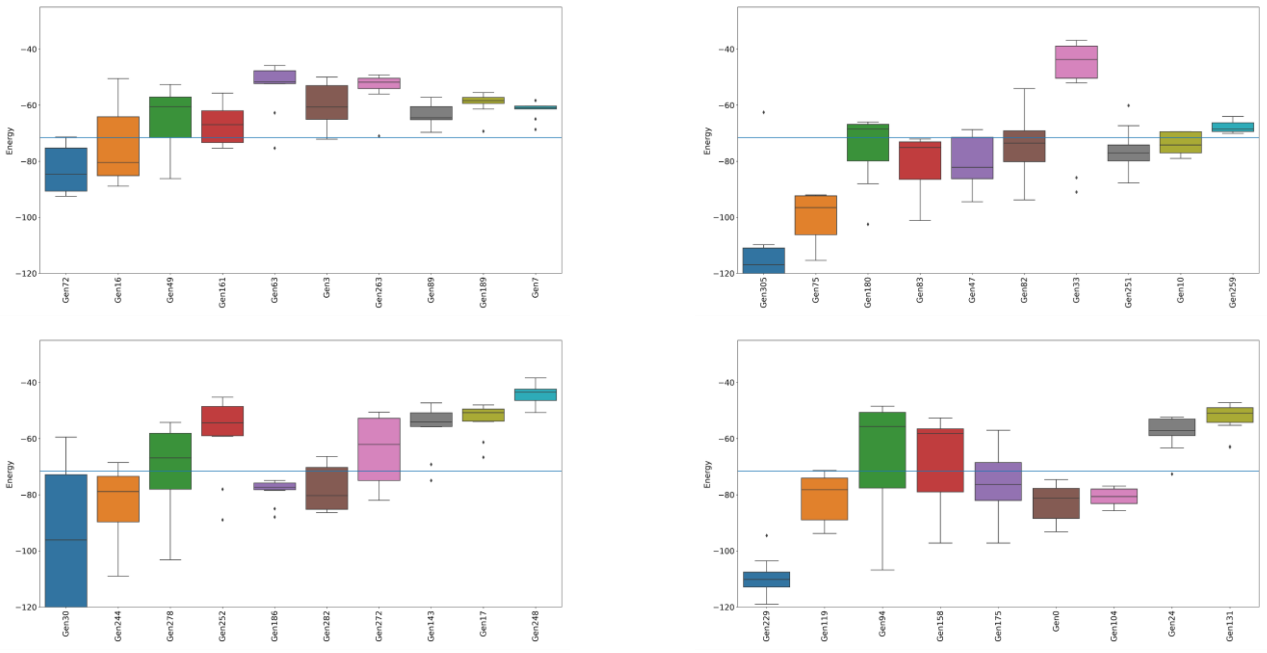


**Figure S2.** Top 10 PELE binding energy distributions of the 39 generated Affilin® sequences targeting HER3 that were selected from the PyDock energy analysis. The blue line indicates the mean binding energy value among all the systems.

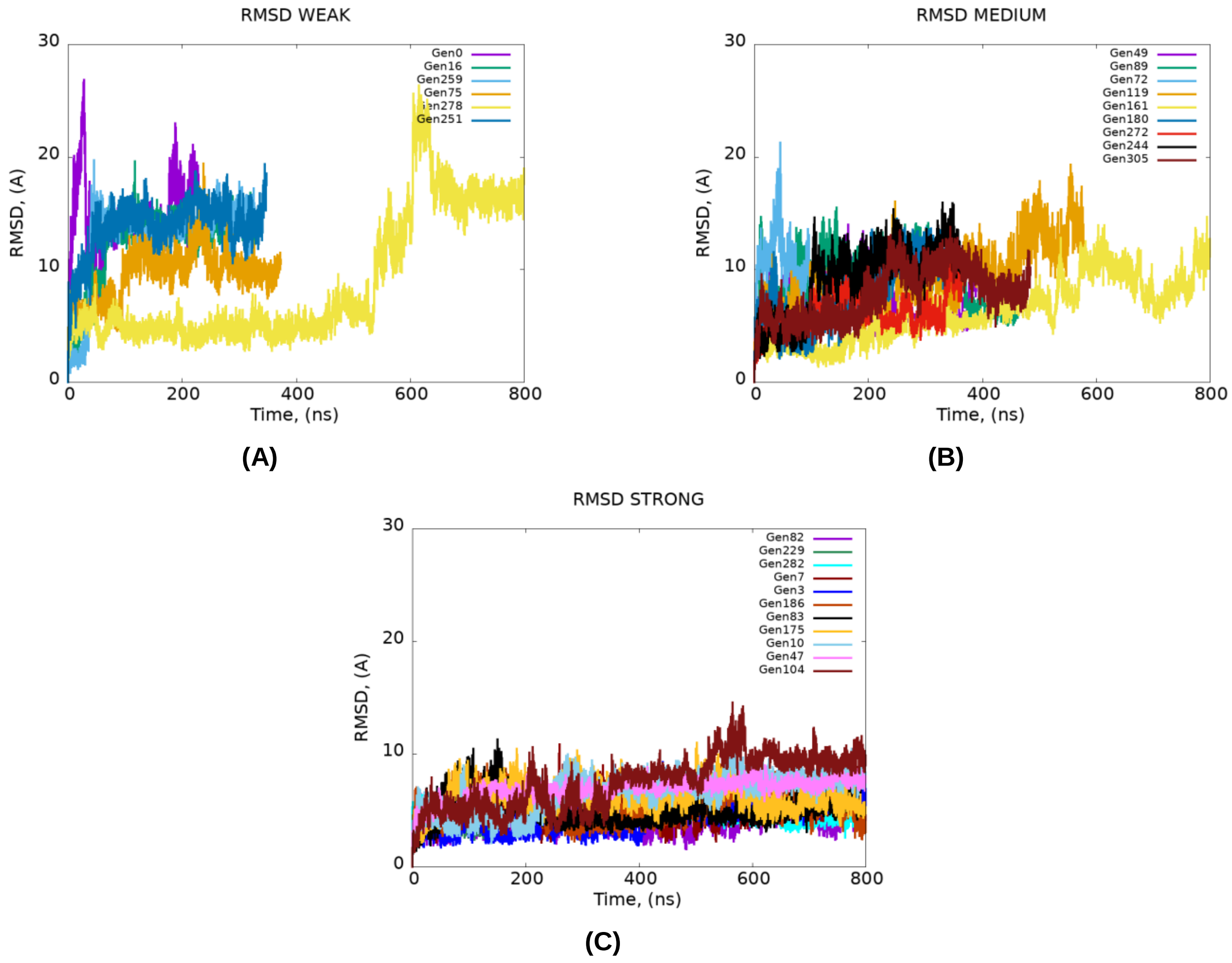


**Figure S3.** Affilin® backbone L-RMSD evolution over time in the MD simulations of the 26 *de novo* Affilin® proteins selected from the PELE analysis. The designs were classified into (**A**) WB, (**B**) MB, or (**C**) SB according to the fluctuation of the RMSD. To reduce the computational cost, the simulations that had high RMSD fluctuations were stopped and classified as WB or MB, explaining why some simulations are shorter. Consequently, the binders that fluctuated around low RMSD values over 800 ns were classified as SB.

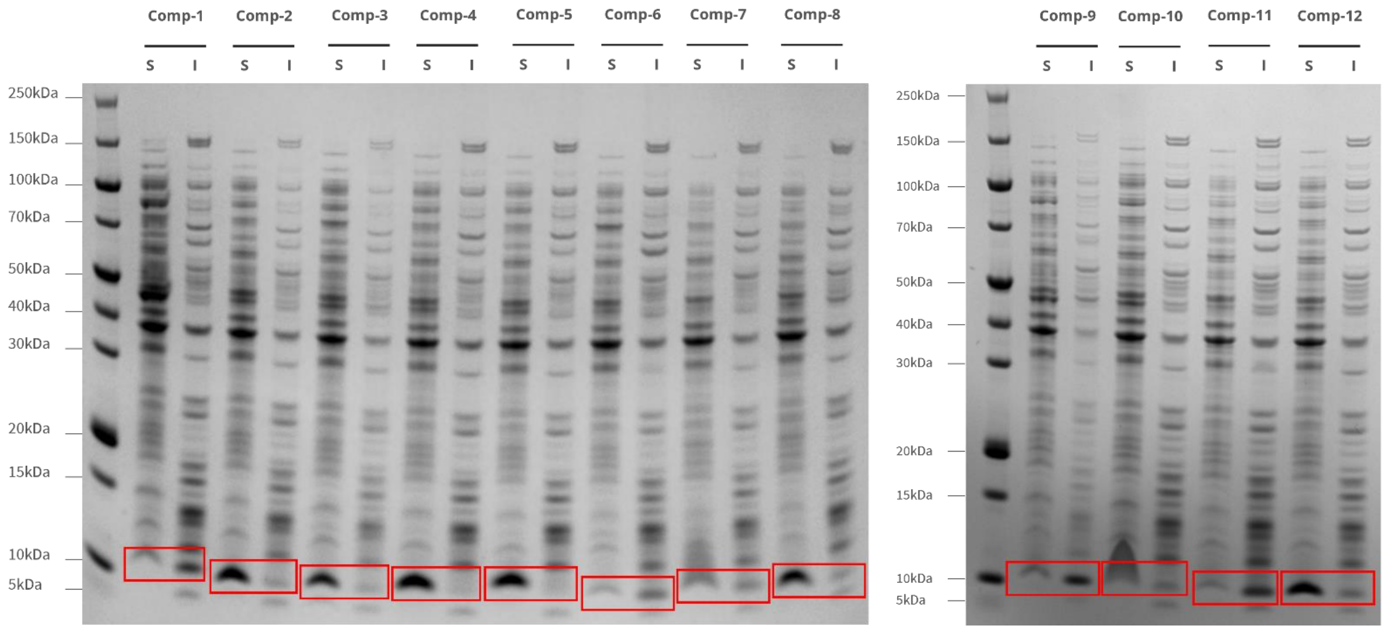


**Figure S4:** Expression of 12 computational identified Affilin® proteins – 11 SBs and 1 WB. SDS-PAGE containing soluble (S) and insoluble (I) fractions for Comp-1 to Comp-12. Pairs of soluble and insoluble fractions of individual Affilin® proteins are highlighted by red boxes.

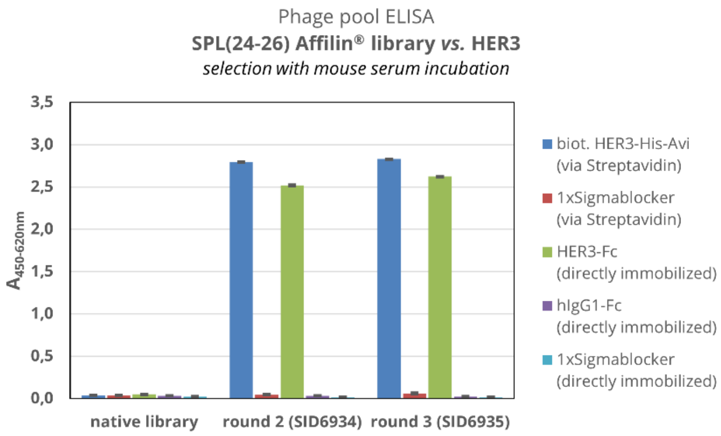


**Figure S5.** Phage pool ELISA for SPL(24-26) Affilin® library after three rounds of selection using phage display technology. The phage pools of the native library (prior to selection), round two and three were tested against HER3 (biotinylated or Fc-tagged) and controls (Sigmablocker and IgG1-Fc) in a 384-well plate. Phage samples were normalized to $5x10^{11}$ phages per well. The readout was taken by using a peroxidase-coupled anti-M13 antibody and an absorption at 450 nm and 620 nm.

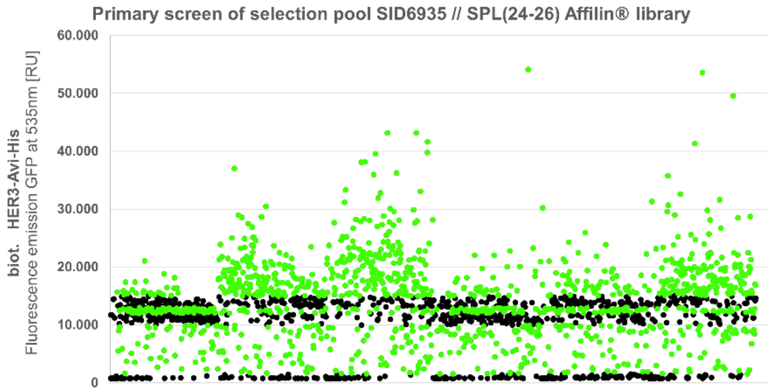


**Figure S6.** Data of ELISA-based high-throughput screen for the selection pool SID6935. Expression lysates of *E. coli* BL21 (DE3) single clones containing GFP-fused Affilin® proteins were analyzed for binding biotinylated HER3-His-Avi immobilized on 384-well plates via Streptavidin. Readout was taken by excitation at 485 nm and emission at 535 nm. 2160 clones were tested and 1192 could be nominated as hit using a manually defined threshold between 1.500-10.000 and 12.000-13.000 and >15.000 RU.

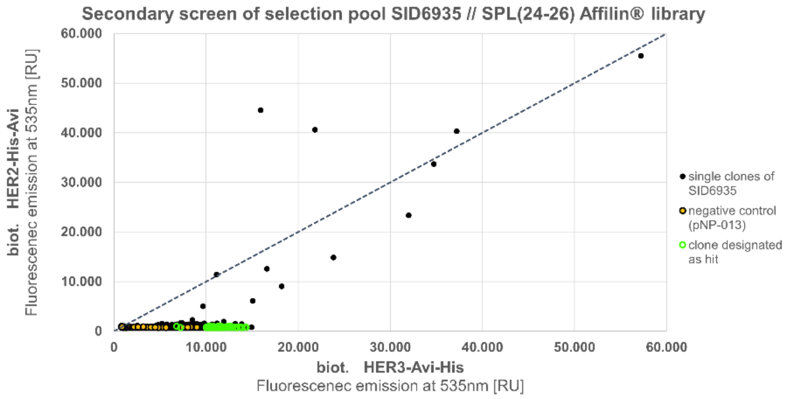


**Figure S7.** Data of ELISA-based high-throughput screen with hits derived from the selection pool SID6935 after the primary screen. Expression lysate of *E. coli* BL21 (DE3) single clones containing GFP-fused Affilin® proteins was analyzed for binding to either biotinylated HER2-His-Avi or biotinylated HER3-His-Avi, both immobilized on 384-well plated via Streptavidin. Readout was taken by excitation at 485 nm and emission at 535 nm. 112 out of 1192 clones could be nominated as hit (signal on HER3 >10.000 RU and signal ratio HER3:HER2 >=10; additionally, two distinct clones < 10.000 RU).

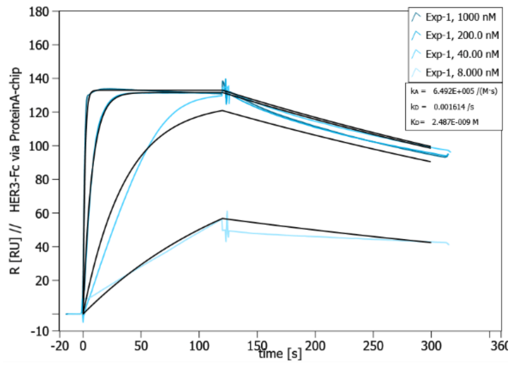


**(A)**

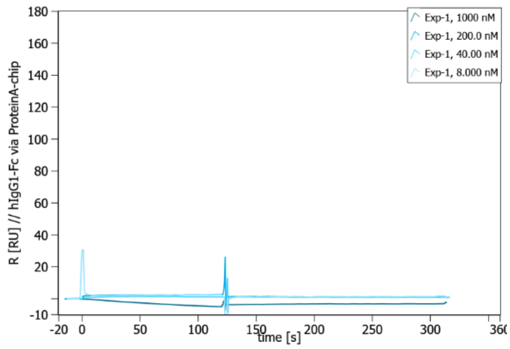


**(B)**

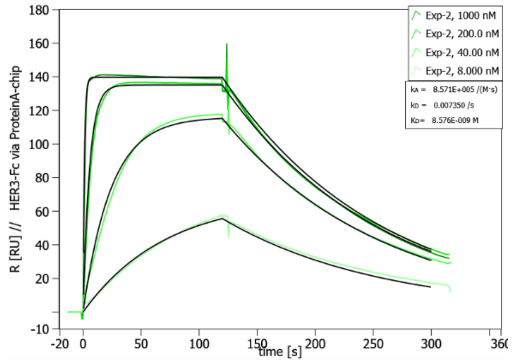


**(C)**

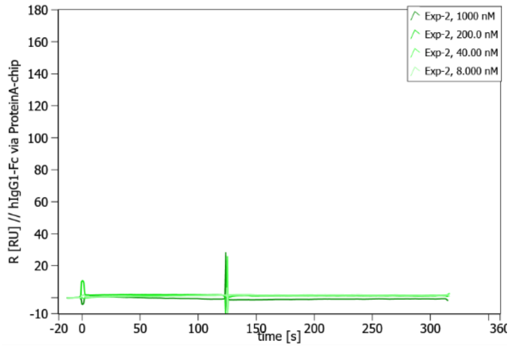


**(D)**

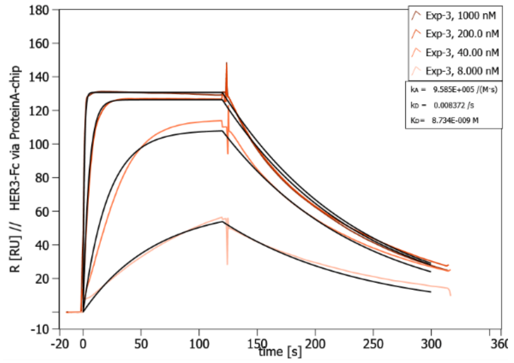


**(E)**

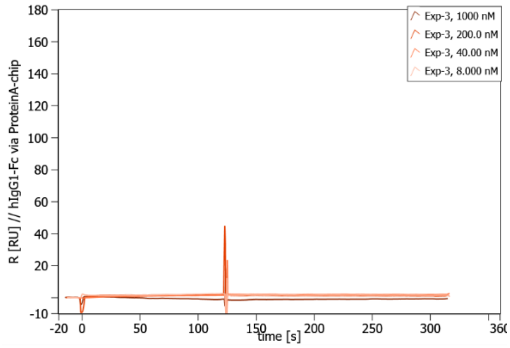


**(F)**

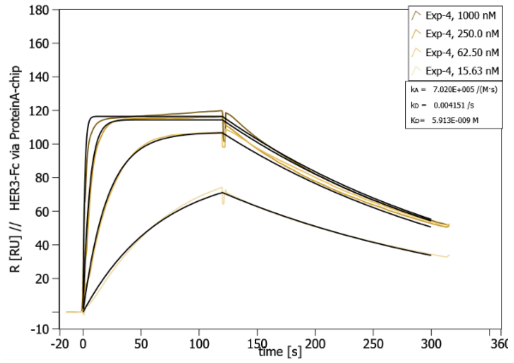


**(G)**

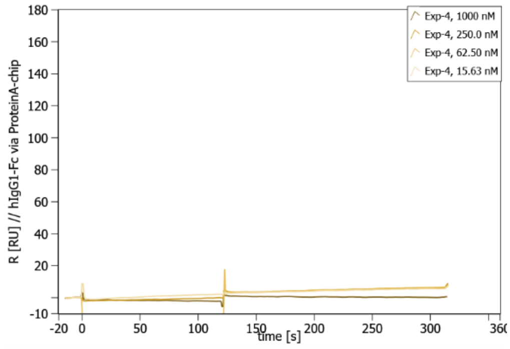


**(H)**

**Figure S8.** Concentration-dependent SPR measurements with calculated affinity shows specific binding to hHER3-Fc of µ-scale purified Exp-1, Exp-2, Exp-3 and Exp-4 Affilin® variant. The measurement was performed on a Sierra SPR-32 (Bruker). **(A), (C), (E), (G)** Fc-tagged HER3 (HER3-Fc) was used as target protein, immobilized via a Protein A-coated chip. **(B), (D), (F), (H)** Human IgG1-Fc protein was used as negative control, immobilized via a Protein A-coated chip.

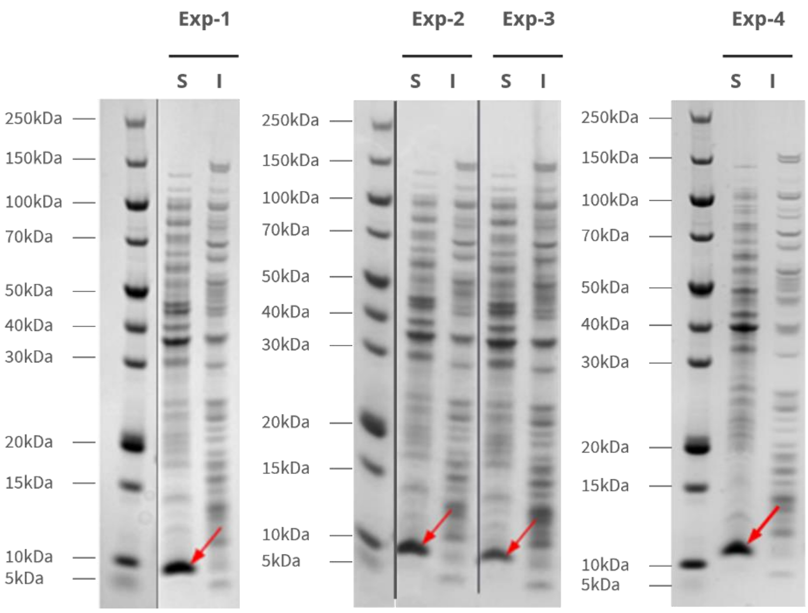


**Figure S9.** Expression of Exp-1 to Exp-4. SDS-PAGE containing soluble (S) and insoluble (I) fractions. Soluble fractions of Affilin® proteins are marked by a red arrow.

```
Affilin® library: MQIFVKTLT------GKTITLEVEPSDTIENVKAKIQDKEGIPPDQQRLIWAGKQLEDGRTLSDYNIxxxxxLxLxLxLRAA
           Exp-1: MQIFVKTLTTTIAPVGKTITLEVEPSDTIENVKAKIQDKEGIPPDQQRLIWAGKQLEDGRTLSDYNIEKDSVLALLLNLRAA
           Exp-2: MQIFVKTLTSNEQYKGKTITLEVEPSDTIENVKAKIQDKEGIPPDQQRLIWAGKQLEDGRTLSDYNISDESILALLLNLRAA
           Exp-3: MQIFVKTLTTQVSPPGKTITLEVEPSDTIENVKAKIQDKEGIPPDQQRLIWAGKQLEDGRTLSDYNIRAHDQLALLLELRAA
           Exp-4: MQIFVKTLTTKYDIPGKTITLEVEPSDTIENVKAKIQDKEGIPPDQQRLIWAGKQLEDGRTLSDYNIVQNSMLALLLNLRAA
```

**Figure S10.** Sequence Alignment of *in vitro* identified Affilin® sequences.

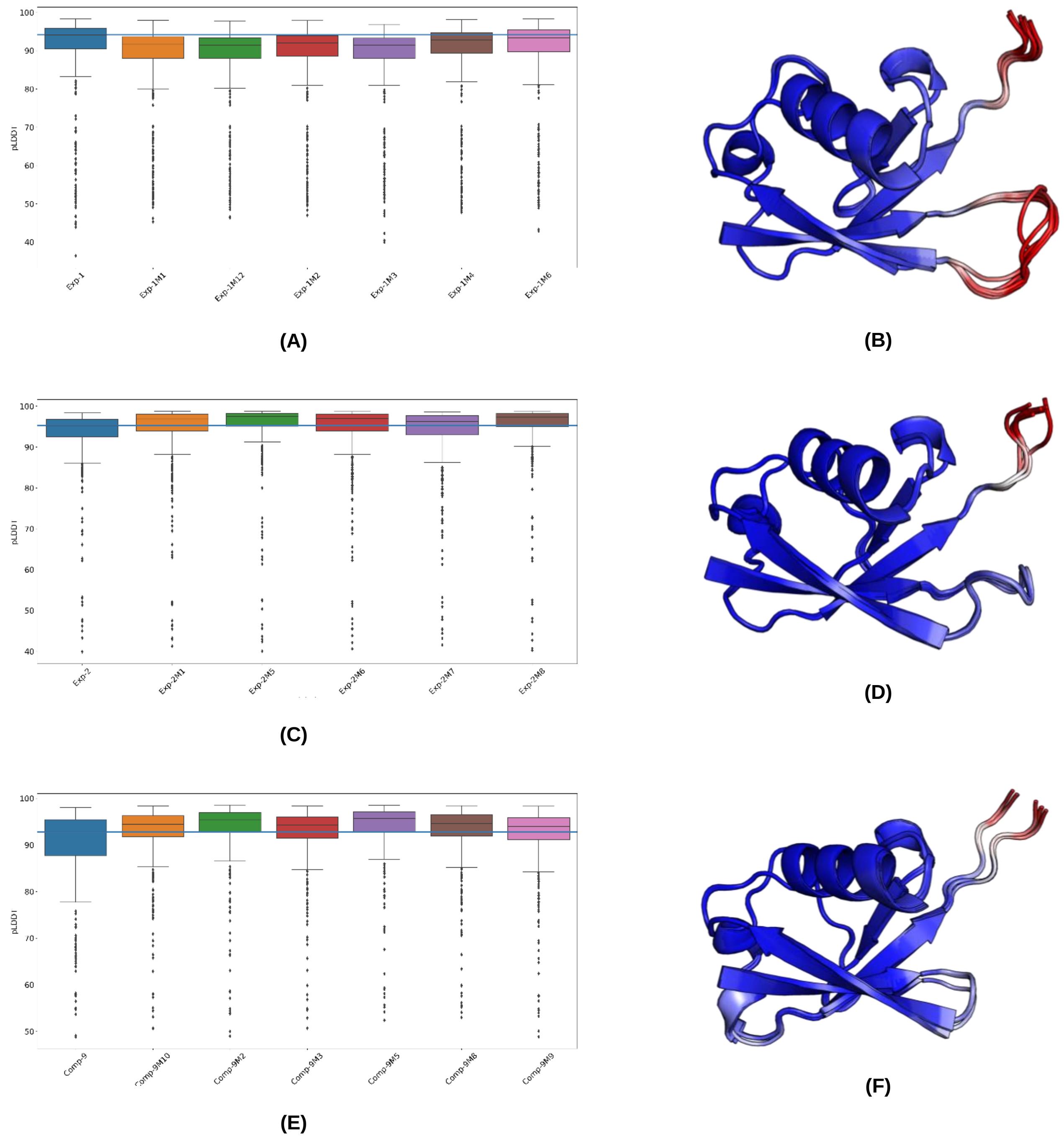


**Figure S11.** AlphaFold2 models confidence scores for the promising ProteinMPNN multiple mutants. (**A, C, E**) Box plots show the pLDDT distribution for all models of Exp-1, Exp-2, and Comp-9 respectively. The blue line indicates the median pLDDT score of the original Affilin® hit. (**B, D, F**) Superposition of the monomeric Affilin® variants of Exp-1, Exp-2, and Comp-9, respectively. Regions in blue are highly confident regions, while regions in red indicate low confidence.

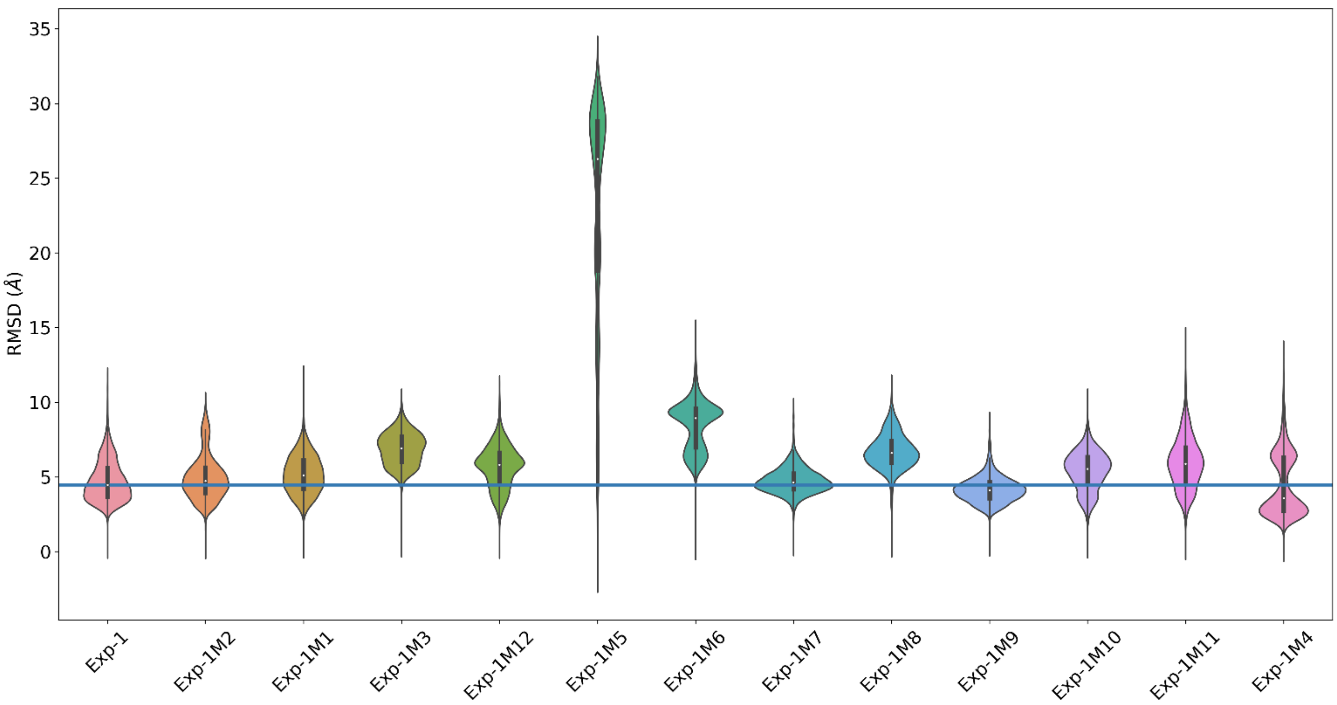

RMSD (Å)
35
30
25
20
15
10
5
0
Exp-1
Exp-1M2
Exp-1M1
Exp-1M3
Exp-1M12
Exp-1M5
Exp-1M6
Exp-1M7
Exp-1M8
Exp-1M9
Exp-1M10
Exp-1M11
Exp-1M4


**(A)**

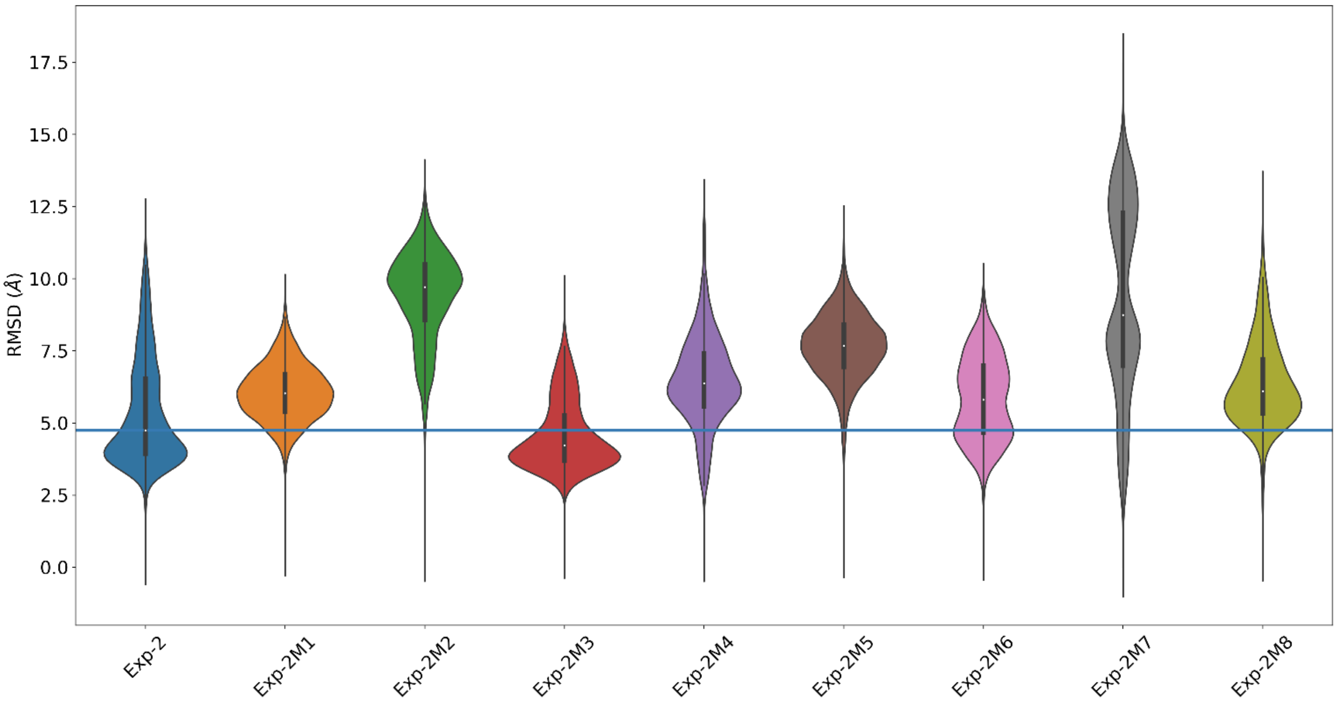

RMSD (Å)
17.5
15.0
12.5
10.0
7.5
5.0
2.5
0.0
Exp-2
Exp-2M1
Exp-2M2
Exp-2M3
Exp-2M4
Exp-2M5
Exp-2M6
Exp-2M7
Exp-2M8


**(B)**

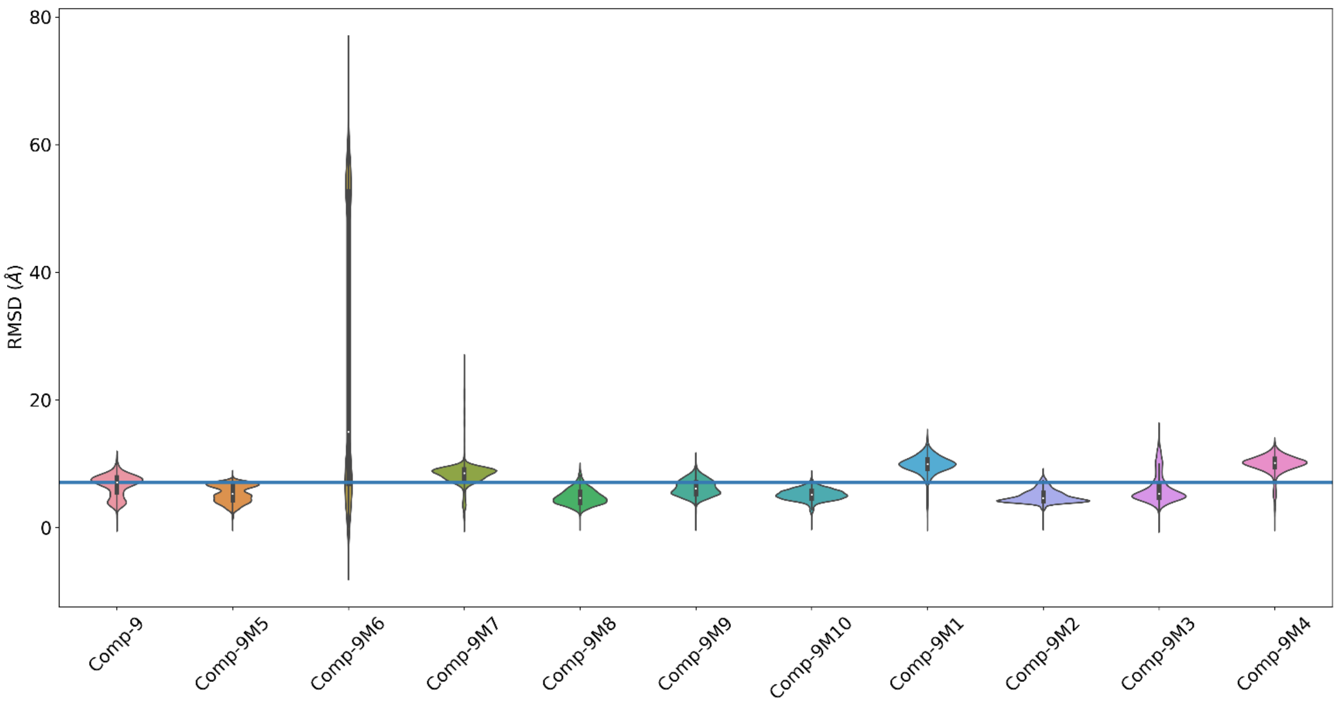


**(C)**

**Figure S12.** Violin plots of the RMSD distribution from the Affilin® protein along the MD simulation of all ProteinMPNN multiple mutants for (**A**) Exp-1, (**B**) Exp-2, and (**C**) Comp-9. The blue line indicates the median RMSD of the original sequence.

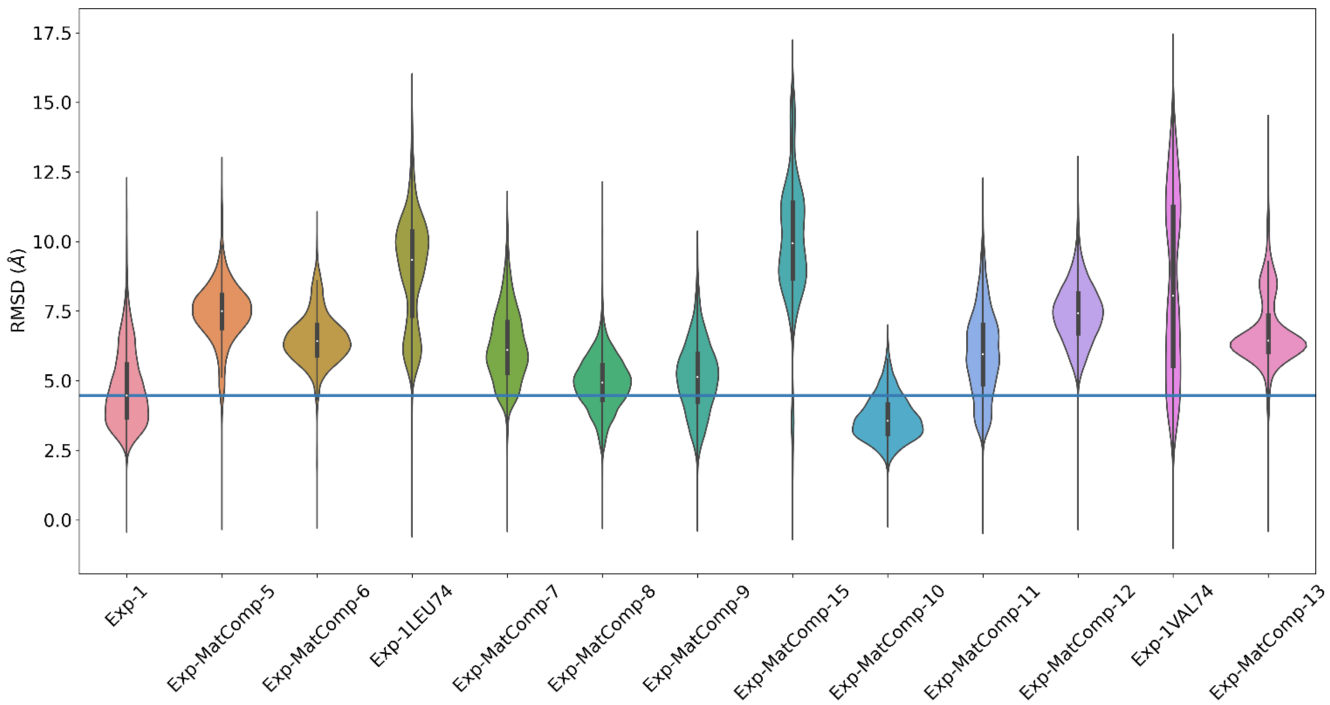


**Figure S13.** Violin plots of the RMSD distribution from the Affilin® protein along the MD simulation of the promising single mutants. The blue line indicates the median RMSD of the original sequence, Exp-1.

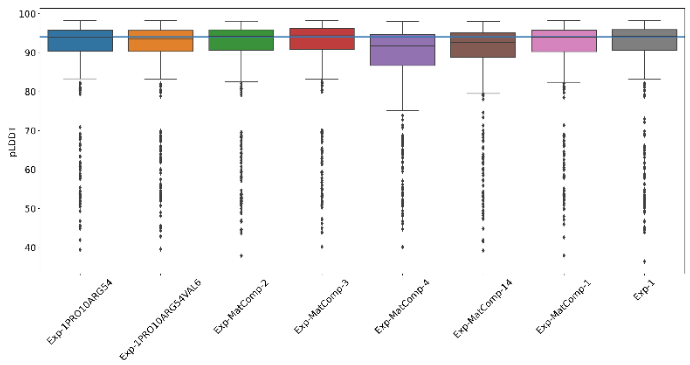


**(A)**

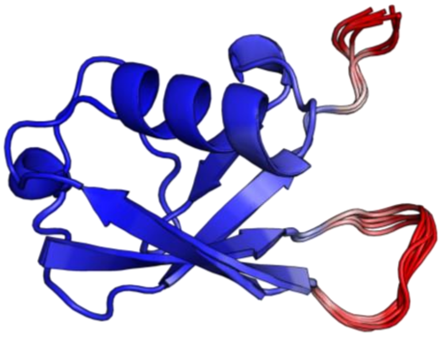

**(B)**

**Figure S14.** AlphaFold2 models confidence scores for the consensus mutants. (**A**) Box plots showing the pLDDT distribution for all models of Exp-1. The blue line indicates the median pLDDT score of the parent Affilin® molecule. (**B**) Superposition of the monomeric Affilin® variants of Exp-1. Regions in blue are highly confident regions, while regions in red indicate low confidence.

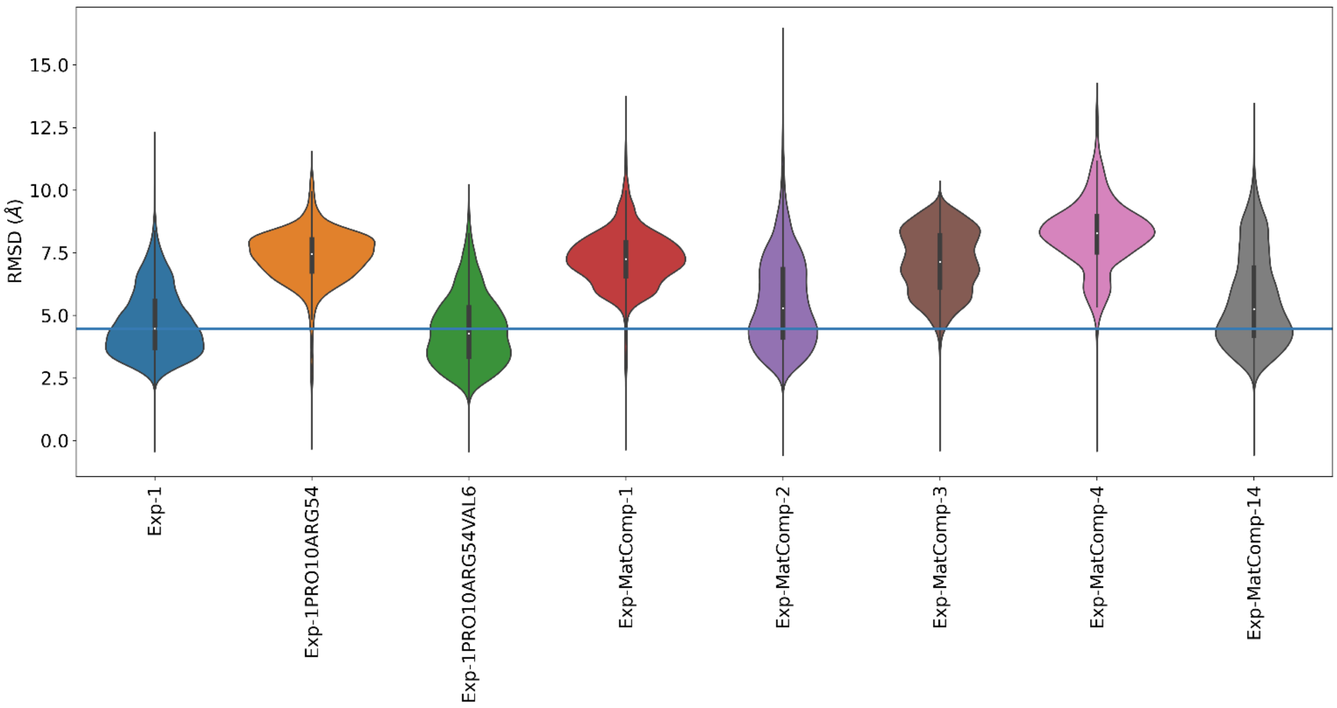


**Figure S15.** Violin plots of the RMSD distribution from the Affilin® protein along the MD simulation of the consensus mutants. The blue line indicates the median RMSD of the original sequence, Exp-1.

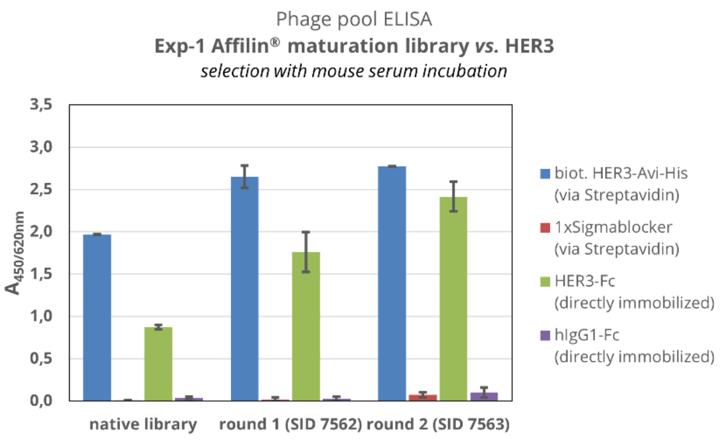


**Figure S16.** Phage pool ELISA for Exp-1 Affilin® maturation library after two rounds of selection using phage display technology. The phage pools of the native library (prior to selection), and round one and two were tested against HER3-Fc or HER3-His-Avi target proteins and controls (Sigmablocker and IgG1-Fc) in a 384-well plate. Phage samples were normalized to $2x10^9$ phages per well. The readout was taken by using a peroxidase-coupled anti-M13 antibody and an absorption at 450 nm and 620 nm.

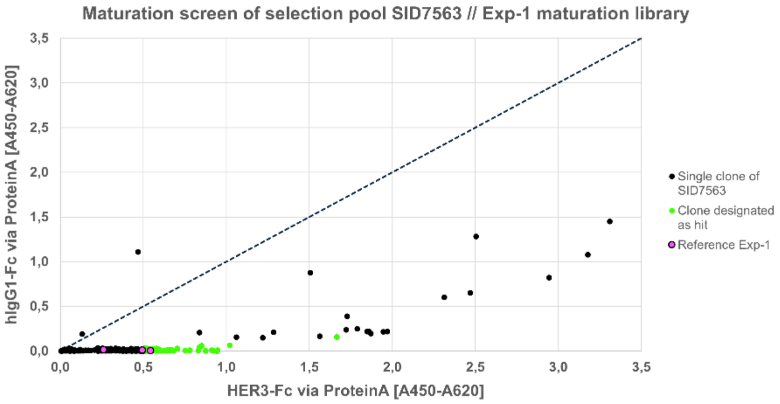


**Figure S17.** Data of ELISA-based maturation screen for the selection pool SID7563 derived from Exp-1 Affilin® maturation library. Expression lysate of *E. coli* BL21 (DE3) single clones containing StrepTag-fused Affilin® proteins was analyzed for binding to either HER3-Fc or hIgG1-Fc, both immobilized on 384-well plates via Protein A. Readout was taken by peroxidase-coupled StrepTactin and an absorption at 450 nm and 620 nm. 100 out of 360 clones with absorption values >0.5 on HER3 and a signal ratio of HER3-Fc:hIgG1-Fc >10 were nominated as hit.

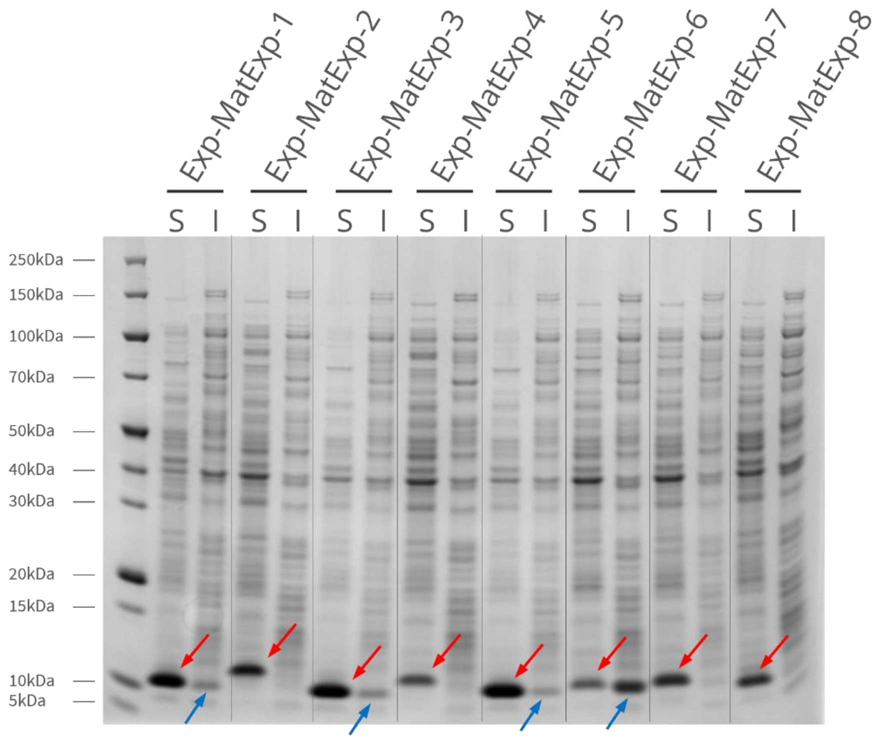


**Figure S18.** Expression of *in vitro* maturated Affilin® variants Exp-MatExp-1 to -8. SDS-PAGE showing soluble (S) and insoluble (I) fractions for each variant. Soluble fractions of Affilin® proteins are indicated by a red arrow. Insoluble fractions are marked by a blue arrow.

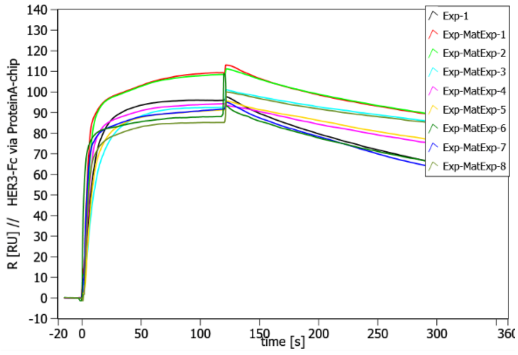


(A)

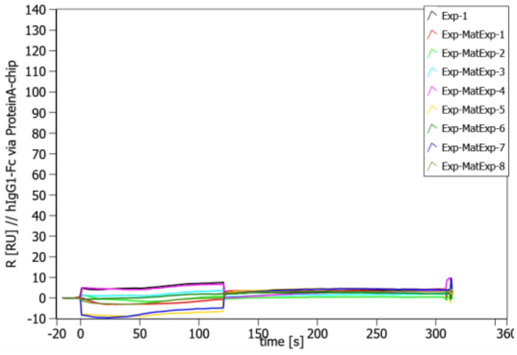


(B)

**Figure S19.** SPR data shows specific binding to HER3-Fc of µ-scale purified Exp-MatExp-1 to Exp-MatExp-8 Affilin® variant. The measurement was performed on a Sierra SPR-32 (Bruker) with 100 nM of Affilin® protein. (**A**) HER3-Fc was used as target protein, immobilized via a Protein A-coated chip. (**B**) Human IgG1-Fc was used as target protein, immobilized via a Protein A-coated chip.

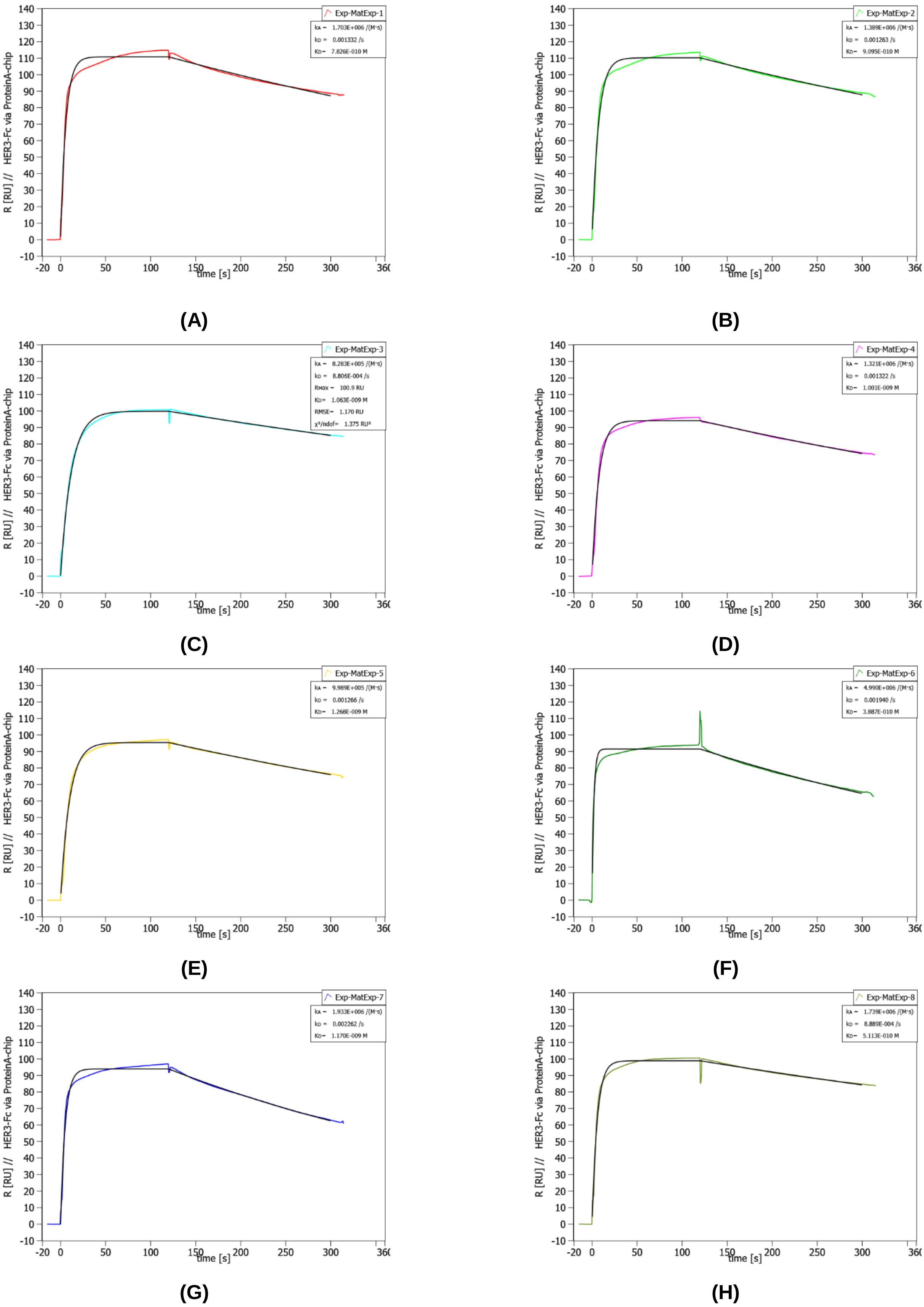


**Figure S20.** SPR data with calculated affinity to HER3-Fc of µ-scale purified Exp-MatExp-1 to Exp-MatExp-8 Affilin® variant (**A**) to (**H**). The measurement was performed on a Sierra SPR-32 (Bruker) with 100 nM of Affilin® protein. HER3-Fc was used as target protein, immobilized via a Protein A-coated chip.

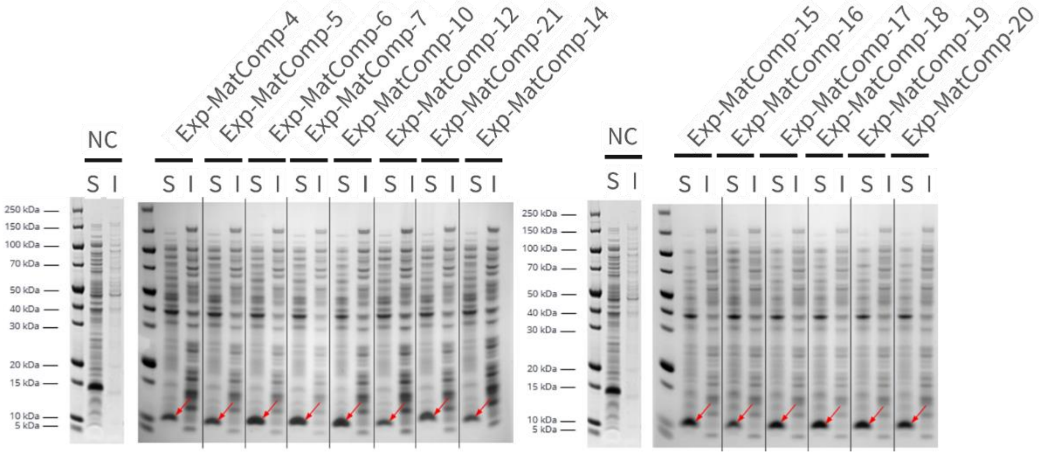


**Figure S21.** Expression of 14 *in silico* maturated Affilin® variants. SDS-PAGE showing soluble (S) and insoluble (I) fractions for each variant. Soluble fractions of Affilin® proteins are marked by a red arrow.

```
         Exp-1: MQIFVKTLTTTIAPVGKTITLEVEPSDTIENVKAKIQDKEGIPPDQQRLIWAGKQLEDGRTLSDYNIEKDSVLALLLNLRAA
  Exp-MatExp-5: MQIFVRTLTTTIAPVGKTITLEVEPSDTIENVKAKIQDKEGIPPDQQRLIWAGKQLEDGRTLSDYNIEKDSVLALLLNLRAA
  Exp-MatExp-8: MQIFVKALTTTIAPVGKTITLEVEPSDTIENVKAKIQDKEGIPPDQQRLIWAGKQLEDGRTLSDYNIEKDSVLALLLNLRAA
 Exp-MatComp-4: MQIFVVTLTTTIAPVGKTITLEVEPSDTIENVKAKIQDKEGIPPDQQRLIWAGRQLEDGRTLSDYNIEKDSVLALLLNLRAA
Exp-MatComp-12: MQIFVVTLTTTIAPVGKTITLEVEPSDTIENVKAKIQDKEGIPPDQQRLIWAGKQLEDGRTLSDYNIEKDSVLALLLNLRAA
```

**Figure S22.** Sequence Alignment of *in silico* and *in vitro* maturated Affilin® sequences.

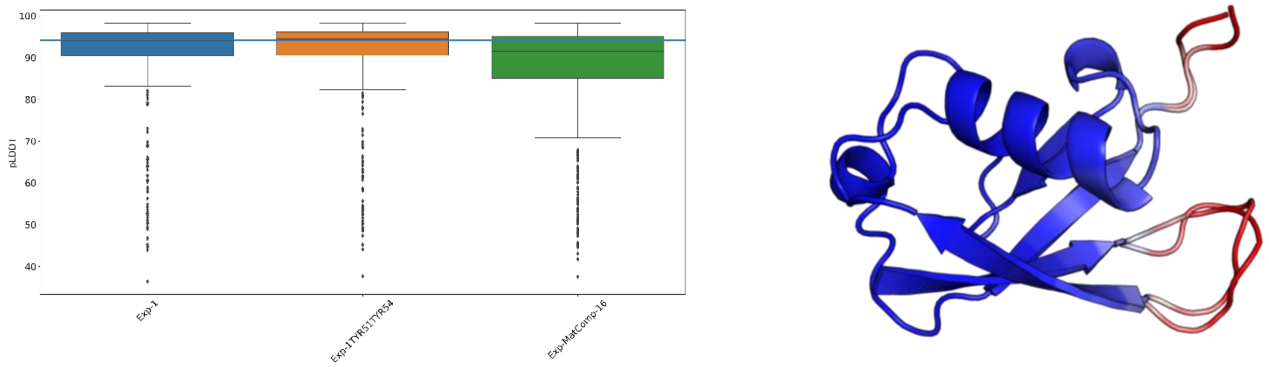


(**A**) (**B**)

**Figure S23.** AlphaFold2 models confidence scores for the double mutants of the second round. (**A**) Box plots showing the pLDDT distribution for all models of Exp-1. The blue line indicates the median pLDDT score of the parent Affilin® molecule. (**B**) Superposition of the monomeric Affilin® variants of Exp-1. Regions in blue are highly confident regions, while regions in red indicate low confidence.

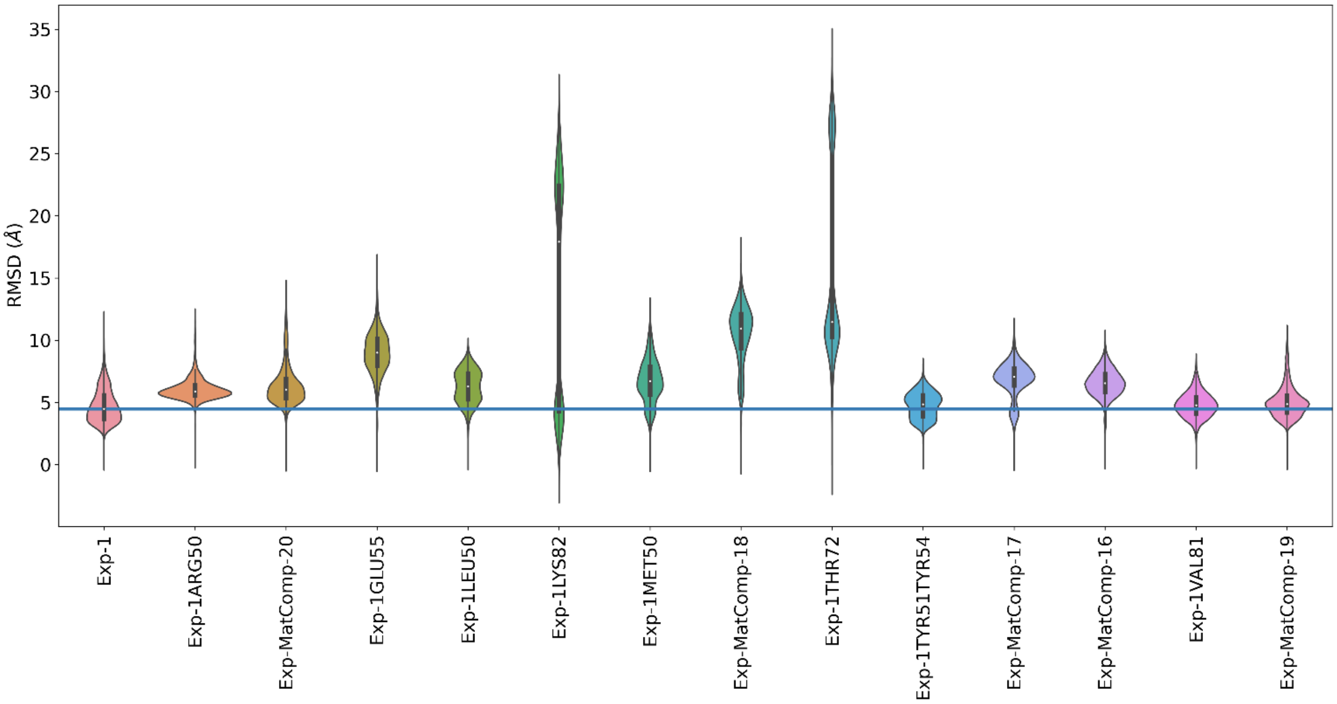


**Figure S24.** Violin plots of the RMSD distribution from the Affilin® protein along the MD simulation of the mutants from the second round. The blue line indicates the median RMSD across all simulations.

### Part II

### Supplementary Information Tables

**Table S1.** Cell binding of lab scale purified Affilin® proteins on HEK293-hHER3-sc87 and SK-BR-3 cells measured by flow cytometry. The measurement was performed on a Guava easyCyte 5HT FACS. StrepTag-fused Affilin® variants were detected using rabbit anti-StrepTag as primary antibody and goat anti-rabbit-IgG-AlexaFluor488 as secondary antibody. Excitation was at 488 nm and emission at 525/30 nm. The concentration of each sample was 1 µM. The given values are x-fold over wildtype ubiquitin control protein (Exp-WT) and empty HEK293-cells.

| Affilin® (lab scale) | Cell binding to HEK293-HER3-sc87 [x-fold over Exp-WT] | Cell binding to SK-BR-3 [x-fold over Exp-WT] |
|---|---|---|
| Exp-1 | 46.03 | 5.16 |
| Exp-2 | 15.82 | 1.91 |
| Exp-3 | 11.32 | 1.67 |
| Exp-4 | 20.68 | 2.23 |
| Exp-WT | 1.0 | 1.0 |

**Table S2.** Cartridges of Affilin® proteins. Cartridges represent the amino acid exchanges in reference to a specific Affilin® library based on the ubiquitin sequence.

| **Affilin®** | **Cartridge** | **Additional side exchanges** |
|---|---|---|
| Exp-1 | [9,6](TTIAPV)EKDSVALN | |
| Exp-2 | [9,6](SNEQYK)SDESIALN | |
| Exp-3 | [9,6](TQVSPP)RAHDQALE | |
| Exp-4 | [9,6](TKYDIP)VQNSMALN | |
| Exp-MatExp-5 | [9,6](TTIAPV)EKDSVALN | (6.K>R) |
| Exp-MatExp-8 | [9,6](TTIAPV)EKDSVALN | (7.T>A) |
| Exp-MatComp-4 | [9,6](TTIAPV)EKDSVALN | (6.K>V),(48.K>R) |
| Exp-MatComp-12 | [9,6](TTIAPV)EKDSVALN | (6.K>V) |

**Table S3.** Change in binding energy of the ProteinMPNN multiple mutants with respect to the parent sequence (ΔΔG) obtained by two methods, MMGBSA (ΔΔGGB, less accurate) and MMPBSA (ΔΔGPB, more accurate). The mutations that improve the binding are highlighted in bold.

| Variant | $\Delta\Delta G_{GB}$ (Kcal/mol) | $\Delta\Delta G_{PB}$ (Kcal/mol) |
|---|---|---|
| Comp-9M1 | 87.380 | 88.312 |
| Comp-9M2 | 20.320 | 32.445 |
| Comp-9M3 | 13.060 | 15.906 |
| Comp-9M4 | 88.090 | 76.575 |
| Comp-9M5 | 63.485 | 79.280 |
| Comp-9M7 | 110.038 | 111.771 |
| Comp-9M8 | 46.665 | 69.074 |
| Comp-9M9 | 27.811 | 22.005 |
| Comp-9M10 | 30.077 | 26.696 |
| **Exp-1M1** | **-33.952** | **-43.799** |
| **Exp-1M2** | **-41.265** | **-46.142** |
| **Exp-1M3** | **-84.385** | **-77.586** |
| **Exp-1M4** | **-67.826** | **-68.861** |
| **Exp-1M6** | **-55.111** | **-59.600** |
| Exp-1M7 | 43.081 | 46.352 |
| Exp-1M8 | 26.792 | 39.806 |
| Exp-1M9 | 14.786 | 18.254 |
| Exp-1M10 | 67.004 | 74.431 |
| Exp-1M11 | 10.346 | 9.001 |
| **Exp-1M12** | **-48.115** | **-55.211** |
| **Exp-2M1** | **-103.964** | **-103.186** |
| **Exp-2M2** | **-8.021** | 19.917 |
| **Exp-2M3** | **-36.389** | **-7.913** |
| **Exp-2M4** | **-4.774** | 18.809 |
| **Exp-2M5** | **-67.311** | **-69.682** |
| **Exp-2M6** | **-88.463** | **-68.624** |
| **Exp-2M7** | **-85.496** | **-62.296** |
| **Exp-2M8** | **-52.565** | **-40.481** |

**Table S4.** Change in binding energy of the single mutants with respect to the parent sequence (ΔΔG) obtained by two methods, MMGBSA (ΔΔGGB, less accurate) and MMPBSA (ΔΔGPB, more accurate). The mutations that improve the binding are highlighted in bold.

| Variant | $\Delta\Delta G_{GB}$ (Kcal/mol) | $\Delta\Delta G_{PB}$ (Kcal/mol) |
|---|---|---|
| **Exp-1ALA6 (Exp-MatComp-5)** | **-25.260** | **-26.789** |
| **Exp-1ARG54** | **-2.654** | 5.480 |
| Exp-1ARG6 | 11.900 | 24.274 |
| Exp-1ASN52 | 0.605 | 20.159 |
| Exp-1GLY10 | 5.510 | 4.007 |
| **Exp-1ILE72 (Exp-MatComp-6)** | **-31.953** | **-21.726** |
| Exp-1ILE74 | 17.870 | 37.947 |
| Exp-1ILE79 | 36.306 | 35.587 |
| **Exp-1ILE9** | **-8.033** | **-6.757** |
| Exp-1LEU6 | 13.051 | 22.532 |
| Exp-1LEU72 | 11.160 | 12.766 |
| **Exp-1LEU74** | **-42.593** | **-20.508** |
| Exp-1LYS79 | 12.526 | 5.713 |
| Exp-1PHE79 | 37.420 | 46.218 |
| Exp-1PRO10 | 11.318 | 17.402 |
| **Exp-1SER10** | **-7.524** | **-3.572** |
| **Exp-1SER52 (Exp-MatComp-7)** | **-85.376** | **-66.204** |
| **Exp-1SER74 (Exp-MatComp-8)** | **-30.209** | **-25.970** |
| **Exp-1THR74 (Exp-MatComp-9)** | **-51.327** | **-53.722** |
| **Exp-1TYR51 (Exp-MatComp-15)** | **-53.778** | **-61.128** |
| **Exp-1TYR54 (Exp-MatComp-10)** | **-109.085** | **-108.305** |
| Exp-1TYR6 | 9.898 | 6.270 |
| **Exp-1VAL10 (Exp-MatComp-11)** | **-57.888** | **-52.519** |
| Exp-1VAL52 | 63.567 | 79.282 |
| **Exp-1VAL6 (Exp-MatComp-12)** | **-18.234** | **-26.731** |
| **Exp-1VAL74** | **-26.861** | **-28.431** |
| **Exp-1VAL9** | **-23.462** | **-12.439** |

**Table S5.** Change in binding energy of the consensus mutants respect to the parent sequence (ΔΔG) obtained by two methods, MMGBSA (ΔΔGGB, less accurate) and MMPBSA (ΔΔGPB, more accurate). The mutations that improve the binding are highlighted in bold.

| Variant | $\Delta\Delta G_{GB}$ (Kcal/mol) | $\Delta\Delta G_{PB}$ (Kcal/mol) |
|---|---|---|
| Exp-1PRO10ARG54 | 8.4792 | 12.2362 |
| Exp-1PRO10ARG54VAL6 | 3.127 | 16.001 |
| **Exp-1PRO10GLN6 (Exp-MatComp-1)** | **-39.912** | **-33.498** |
| **Exp-1PRO10TYR54 (Exp-MatComp-2)** | **-60.720** | **-58.854** |
| **Exp-1PRO10TYR54VAL6 (Exp-MatComp-3)** | **-50.449** | **-41.641** |
| **Exp-1VAL6ARG54 (Exp-MatComp-4)** | **-14.552** | **-18.332** |
| Exp-1VAL6TYR54 (Exp-MatComp-14) | 8.023 | 8.683 |

**Table S6.** Summary of *in vitro* maturated, µ-scale purified Affilin® proteins. SPR data was obtained with 100 nM Affilin® protein and Fc-tagged HER3 (HER3-Fc) immobilized to a Protein A-chip using a Sierra SPR-32 (Bruker). Cell binding data was obtained by flow cytometry using a Guava easyCyte 5HT FACS with StrepTag-fused Affilin® variants at a concentration of 500 nM, which were detected by a rabbit anti-StrepTag as primary antibody and goat anti-rabbit-IgG-AlexaFlour488 as secondary antibody and an excitation at 488 nm and emission at 525/30 nm. The values given are referred to Exp-1 data and are shown as x-fold of Exp-1.

| | $K_D$ – Binding affinity vs HER3-Fc (µ-scale) [nM] | | | |
|---|---|---|---|---|
| **Affilin® (µ-scale)** | **SPR** | **SPR [x-fold over Exp-1]** | **Cell binding to HEK293-HER3-sc87 [x-fold over Exp-1]** | **Cell binding to SK-BR-3 [x-fold over Exp-1]** |
| Exp-MatExp-1 | 0.8 | 2.9 | 1.7 | 2.0 |
| Exp-MatExp-2 | 0.9 | 2.6 | 1.4 | 1.7 |
| Exp-MatExp-3 | 1.1 | 2.1 | 1.6 | 1.9 |
| Exp-MatExp-4 | 1.0 | 2.3 | 1.6 | 2.0 |
| Exp-MatExp-5 | 1.3 | 1.8 | 1.5 | 2.0 |
| Exp-MatExp-6 | 0.4 | 5.8 | 1.0 | 1.2 |
| Exp-MatExp-7 | 1.2 | 1.9 | 1.6 | 1.7 |
| Exp-MatExp-8 | 0.5 | 4.6 | 1.5 | 1.4 |

**Table S7.** Experimental validation of computational maturated Affilin® proteins by using different methods. Either µ-scale or lab scale purified Affilin® variants were prepared in a concentration-dependent manner for SPR or cell binding measurements. SPR measurement was performed on a Sierra SPR-32 (Bruker) with Fc-tagged HER3 (HER3-Fc) immobilized on a Protein A-coated chip. Cell binding was performed by flow cytometry on a Guava easyCyte 5HT FACS with hHER3-overexpressing HEK293 cells (HEK293-HER3-sc87). StrepTag-fused Affilin® variants were detected using rabbit anti-StrepTag as primary antibody and goat anti-rabbit-IgG-AlexaFlour488 as secondary antibody. Excitation at 488 nm and emission at 525/30 nm. Table abbreviations: nd – not determined

| ***In silico* round** | **Affilin®** | **$K_D$ – Binding affinity vs HER3-Fc [nM]** | | **Cell binding to HEK293-HER3-sc87 [nM]** |
|---|---|---|---|---|
| | | **SPR (µ-scale)** | **SPR (lab scale)** | |
| 1 | Exp-MatComp-4 | 0.7 | 0.2 | 0.6 |
| 1 | Exp-MatComp-5 | 14.4 | 7.3 | *nd* |
| 1 | Exp-MatComp-6 | 6.0 | 7.3 | *nd* |
| 1 | Exp-MatComp-7 | 5.8 | 6.2 | *nd* |
| 1 | Exp-MatComp-10 | 4.0 | 10.5 | *nd* |
| 1 | Exp-MatComp-12 | 0.8 | 1.1 | 0.6 |
| 2 | Exp-MatComp-14 | *nd* | 0.7 | 0.9 |
| 2 | Exp-MatComp-15 | *nd* | 19.2 | *nd* |
| 2 | Exp-MatComp-16 | *nd* | 1.1 | 1.0 |
| 2 | Exp-MatComp-17 | *nd* | 16.8 | *nd* |
| 2 | Exp-MatComp-18 | *nd* | 498.0 | *nd* |
| 2 | Exp-MatComp-19 | *nd* | 5.0 | *nd* |
| 2 | Exp-MatComp-20 | *nd* | 4.8 | *nd* |
| - | Exp-1 | 2.5 | 5.4 | *nd* |

**Table S8.** Cell binding by flow cytometry of lab scale purified Affilin® proteins on HEK293-HER3-sc87 and SK-BR-3 cells. The measurement was performed on a Guava easyCyte 5HT FACS. StrepTag-fused Affilin® variants were detected using rabbit anti-StrepTag as primary antibody and goat anti-rabbit-IgG-AlexaFlour488 as secondary antibody. Excitation at 488 nm and emission at 525/30 nm. The concentration of Affilin® proteins was 100 nM. The values given refer to Exp-1 data and are shown as x-fold over Exp-1.

| Affilin® (lab scale) | Cell binding to HEK293-HER3-sc87 [x-fold over Exp-1] | Cell binding to SK-BR-3 [x-fold over Exp-1] |
|---|---|---|
| Exp-MatExp-5 | 1.3 | 1.8 |
| Exp-MatExp-8 | 1.5 | 2.1 |
| Exp-MatComp-4 | 1.5 | 1.9 |
| Exp-MatComp-12 | 1.5 | 1.8 |
| Exp-1 | 1.0 | 1.0 |

**Table S9.** Change in binding energy of the second-round mutants respect to the parent sequence (ΔΔG) obtained by two methods, MMGBSA (ΔΔGGB, less accurate) and MMPBSA (ΔΔGPB, more accurate). The mutations that improve the binding are highlighted in bold.

| Variant | $\Delta\Delta G_{GB}$ (Kcal/mol) | $\Delta\Delta G_{PB}$ (Kcal/mol) |
|---|---|---|
| Exp-1ARG50 | 9.754 | 29.590 |
| **Exp-1GLN82 (Exp-MatComp-20)** | **-99.05** | **-88.904** |
| Exp-1GLU55 | 33.955 | 46.226 |
| Exp-1LEU50 | 82.213 | 84.208 |
| **Exp-1LYS82** | **-56.462** | **-55.383** |
| Exp-1MET50 | 37.225 | 28.753 |
| **Exp-1THR50 (Exp-MatComp-18)** | **-53.991** | **-60.266** |
| **Exp-1THR72** | **-41.285** | **-36.393** |
| Exp-1TYR51TYR54 | 31.276 | 23.456 |
| **Exp-1VAL50 (Exp-MatComp-17)** | **-72.508** | **-77.081** |
| **Exp-1VAL82 (Exp-MatComp-19)** | **-65.814** | **-71.230** |
| **Exp-1VAL6TYR51 (Exp-MatComp-16)** | **-59.472** | **-69.687** |
| Exp-1VAL81 | 85.274 | 75.393 |

**Table S10.** Cloning of Affilin® variants. Exp-MatComp-1 - Exp-MatComp-4 were ordered as String fragments while Exp-MatComp-5 - Exp-MatComp-20 were cloned via PCR. Specific oligonucleotides are given in Table S11.

| Affilin® variant | Cloning method | PCR template | fwd-Primer | rev-Primer |
|---|---|---|---|---|
| Exp-MatComp-1 | String | | | |
| Exp-MatComp-2 | String | | | |
| Exp-MatComp-3 | String | | | |
| Exp-MatComp-4 | String | | | |
| Exp-MatComp-5 | PCR | Exp-1 | 6174_226967_K6A_BsaI_fw | 6173_226967_BsaI_rv |
| Exp-MatComp-6 | PCR | Exp-1 | 1359_SPV-fw2-Bsa | 6179_229730_BsaI_rv |
| Exp-MatComp-7 | PCR1<br>PCR2 | Exp-1<br>Exp-1 | 1359_SPV-fw2-Bsa<br>6182_229731_A52S_BsaI_fw | 6181_229731_BsaI_rv<br>6173_226967_BsaI_rv |
| Exp-MatComp-8 | PCR | Exp-1 | 1359_SPV-fw2-Bsa | 6178_229732_BsaI_rv |
| Exp-MatComp-9 | PCR | Exp-1 | 1359_SPV-fw2-Bsa | 6180_229733_BsaI_rv |
| Exp-MatComp-10 | PCR1<br>PCR2 | Exp-1<br>Exp-1 | 1359_SPV-fw2-Bsa<br>6183_229734_K54Y_BsaI_fw | 6181_229731_BsaI_rv<br>6173_226967_BsaI_rv |
| Exp-MatComp-11 | PCR | Exp-1 | 6177_229735_BsaI_fw | 1413_Wubi-AA-rev-Bsa3 |
| Exp-MatComp-12 | PCR | Exp-1 | 6176_229736_BsaI_fw | 6173_226967_BsaI_rv |
| Exp-MatComp-13 | PCR | Exp-1 | 6175_229737_BsaI_fw | 1413_Wubi-AA-rev-Bsa3 |
| Exp-MatComp-21 | PCR | Exp-2 | 6344_230501_BsaI_fw | 6173_226967_BsaI_rv |
| Exp-MatComp-14 | PCR | Exp-MatComp-2 | 6345_230502_BsaI_fw | 6173_226967_BsaI_rv |
| Exp-MatComp-15 | PCR1<br>PCR2 | Exp-1<br>Exp-1 | 464_SPV-fw-Bsa<br>6384_230626_BsaI_fw | 6385_230626_BsaI_rv<br>1413_Wubi-AA-rev-Bsa3 |
| Exp-MatComp-16 | PCR1<br>PCR2 | Exp-MatComp-12<br>Exp-1 | 464_SPV-fw-Bsa<br>6384_230626_BsaI_fw | 6385_230626_BsaI_rv<br>1413_Wubi-AA-rev-Bsa3 |
| Exp-MatComp-17 | PCR1<br>PCR2 | Exp-1<br>Exp-1 | 464_SPV-fw-Bsa<br>6386_230628_BsaI_fw | 6387_230628_BsaI_rv<br>1413_Wubi-AA-rev-Bsa3 |
| Exp-MatComp-18 | PCR1<br>PCR2 | Exp-1<br>Exp-1 | 464_SPV-fw-Bsa<br>6388_230629_BsaI_fw | 6387_230628_BsaI_rv<br>1413_Wubi-AA-rev-Bsa3 |
| Exp-MatComp-19 | PCR | Exp-1 | 1359_SPV-fw2-Bsa | 6389_230630_BsaI_rv |
| Exp-MatComp-20 | PCR | Exp-1 | 1359_SPV-fw2-Bsa | 6390_230631_BsaI_rv |

**Table S11.** Cloning of Affilin® variants. Specific oligonucleotides used for mutagenesis.

| Name | 5'-3' sequence | Name | 5'-3' sequence |
|---|---|---|---|
| 1359_SPV-fw2-Bsa | AGGAGGGTCTCTAATGCAGATCTTCGTGAAAACC | 6182_229731_A52S_BsaI_fw | TATGGTCTCCTTGGAGCGGTAAACAGCTGGAAGATG |
| 1413_Wubi-AA-rev-Bsa3 | ATATATGGTCTCGGCACTGGCCGCACGCAG | 6183_229734_K54Y_BsaI_fw | TATGGTCTCCTTGGGCGGGTTACCAGCTGGAAGATGGTCG |
| 464_SPV-fw-Bsa | AGGAGGGTCTCTAATGCAGATCTTCGTG | 6344_230501_BsaI_fw | TATGGTCTCTAATGCAGATCTTCGTGGTGACCCTGACCTCTAACG |
| 6173_226967_BsaI_rv | ATATATGGTCTCGGCACTGGCCGCACG | 6345_230502_BsaI_fw | TATGGTCTCAAATGCAGATCTTCGTGGTGACCCTGACAACCACCATTGCACCGGTTG |
| 6174_226967_K6A_BsaI_fw | TATGGTCTCTAATGCAGATCTTCGTGGCAACCCTGACCACTACTATC | 6384_230626_BsaI_fw | TATGGTCTCAGATTTATGCGGGTAAACAGCTGGAAG |
| 6175_229737_BsaI_fw | TATGGTCTCTAATGCAGATCTTCGTGAAAACCCTGGTGACTACTATCGCTCCGG | 6385_230626_BsaI_rv | TATGGTCTCAAATCAGACGCTGC |
| 6176_229736_BsaI_fw | TATGGTCTCTAATGCAGATCTTCGTGGTGACCCTGACCACTACTATC | 6386_230628_BsaI_fw | TATGGTCTCATCTGGTTTGGGCGGGTAAACAG |
| 6177_229735_BsaI_fw | TATGGTCTCTAATGCAGATCTTCGTGAAAACCCTGACCGTGACTATCGCTCCGGTTG | 6387_230628_BsaI_rv | TATGGTCTCACAGACGCTGCTGAT |
| 6178_229732_BsaI_rv | TATGGTCTCGGCACTGGCCGCACGCAGGTTCAGCAGCAGGCTCAGAACACTATCTTTTTCGATG | 6388_230629_BsaI_fw | TATGGTCTCATCTGACCTGGGCGGGTAAACAGCTG |
| 6179_229730_BsaI_rv | TATGGTCTCGGCACTGGCCGCACGCAGGTTCAGCAGCAGTGCCAGGATACTATCTTTTTCGATGTTATAATCG | 6389_230630_BsaI_rv | ATATATGGTCTCGGCACTCACCGCACGCAGGTTC |
| 6180_229733_BsaI_rv | TATGGTCTCGGCACTGGCCGCACGCAGGTTCAGCAGCAGGGTCAGAACACTATCTTTTTCGATG | 6390_230631_BsaI_rv | ATATATGGTCTCGGCACTCTGCGCACGCAGGTTCAG |
| 6181_229731_BsaI_rv | TATGGTCTCCCCAAATCAGACGCTGC | | |

### Part III

### Supplementary Notes

#### NOTE 1. BENCHMARKING THE MOLECULAR MODELING PIPELINE

We conducted retrospective studies to establish a molecular modeling (MM) pipeline (Figure SN1a) capable of distinguishing binders from non-binders (NB) proteins in order to filter AI-generated sequences. Our pipeline integrates state-of-the-art MM software, with advanced predictors of protein-protein interaction regions. By combining these methodologies, we can predict multiple binding modes associated with identified interaction sites, effectively capturing the dynamic nature of protein binding. The resulting data is then used to analyze interactions energetically and classify protein binders.

To assess the pipeline for Affilin® proteins, we used two retrospective targets. One retrospective target was the fibronectin (FINC) extra-domain B (ED-B), for which a crystal structure between the target and a strong binder is available (PDB: 8PEQ). The second retrospective target was HER2, which was selected because of its similarity with the prospective system, HER3.1. Affilin® protein targeting FINC's ED-B domain

##### *1. Affilin® protein targeting FINC's ED-B domain*

##### *1.1. Comparing MODELLER and AlphaFold2 models against the Affilin® crystal*

We began by structurally modeling all available monomeric Affilin® proteins targeting ED-B. These encompassed 5 strong binders (SB), 12 moderate binders (MB), 7 weak binders (WB), and 11 Affilin® proteins specifically designed to target EGFR (comprising 1 SB, 3 MB, and 7 WB), identified as NB. The modeling was performed using both AlphaFold2, simulating a scenario without prior structural knowledge, and MODELLER[1,2], which used the FINC-Affilin® crystal structure (PDB: 8PEQ) as template. Unsurprisingly, since MODELLER used the crystal structure as a template, it produced lower Cα-RMSD values respect to the crystal than AlphaFold2. Nevertheless, AlphaFold2 models exhibited high confidence, with an average predicted local-distance difference test (pLDDT) of 91.83 (Figure SN3). This high confidence is probably due to the structure conservation of ubiquitin-like proteins used as AlphaFold2 templates and the depth of the multiple sequence alignment (3,231 sequences on average, see Table SN1) [3].

Given the high confidence of AlphaFold2, we also generated AlphaFold2 Multimer models and compared them with the reference FINC-Affilin® complex. The best AlphaFold2 Multimer models had an average L-RMSD of 25.13 Å (Figure SN4), and high PAE in the relative orientation between FINC-Affilin® proteins regardless of the binder type (Figure SN5), indicating low confidence in predicting the complex binding mode [4].

Due to this limitation, we opted to extensively sample binding modes using PyDock [5], a widely used rigid body protein-protein docking (PPD) software.

##### *1.2. Differentiating binders and non-binders using PPD energies*

Solely using the top 50 PyDock energies of each system, we consistently distinguished between binders and NB regardless of whether using MODELLER or AlphaFold2 models (Figures SN1b and SN6a; see Figures SN7 and SN8 for all PyDock energy plots). Since our goal was identifying new SBs, we incorporated more precise calculations, such as Monte Carlo (MC) and Molecular Dynamics (MD) simulations, to refine our predictions and distinguish SBs from MBs and WBs. However, to enhance computational efficiency, we needed to reduce the number of docking poses while retaining relevant ones. This led us to introduce a filtering and clustering step.

##### *1.3. Filtering PPD poses*

PPD software exhaustively samples binding modes between the receptor (e.g., FINC) and the ligand (e.g., Affilin® proteins), scoring them by binding energy. While the most relevant binding modes should be chosen according to the score, it is widely accepted that PPD software introduces substantial false positives. To mitigate this, we implemented a prefiltering step that prioritized docking poses based on the proximity of interacting patches.

To accomplish this, we predicted interacting regions of the receptor and ligand proteins using two software: i) PyDockODA [6], which analyzes the protein surface areas with favorable energy change when buried upon protein-protein association, and ii) MaSIF [7], a tool that employs a deep learning method to decipher surface fingerprints, focusing on chemical and geometric features crucial for biomolecular interactions. Figure SN9 shows strong agreement between the interaction patches predicted by both tools. Subsequently, we filtered PPD poses by ensuring that interacting patches were within 10 Å of each other (see Methods in the main manuscript). This approach reduced the number of poses by approximately 19% while retaining the low energy binding modes (Figures SN9e and SN10a). Additionally, filtering reduced the dispersion of Affilin® L-RMSD distributions relative to the crystal structure (Figures SN9f and SN10b), suggesting that binding modes were more consistent with experimental data.

##### *1.4. Discriminating Strong Binders via PELE and MD Simulations*

To refine binder classification, we included MC simulations with PELE [8], which allows a more robust all-atom semi-flexible molecular simulation. Specifically, we performed PELE simulations using the top 50 filtered non-redundant PyDock poses, supplemented with non-redundant AlphaFold2 Multimer models (L-RMSD > 2 Å) to increase diversity. PELE simulations distinguished NB from binders (Figure SN1c) but did not fully separate SBs from MBs and WBs (see Figures SN11 and SN12 for all PELE energy plots).

To address this, we ran 400 ns MD simulations from the lowest energy binding modes identified by PELE. The Affilin® protein's RMSD evolution over time allowed classifying binder types (Figure SN1d), namely: the stronger the binder, the lower RMSD values around which the Affilin® protein's RMSD fluctuates over time. Quantitatively, the strong Affilin® binders maintained RMSD below 5 Å during the entire simulation, indicating that it strongly interacts with its target protein. On the other hand, MB and the WB fluctuated around RMSD values of 5 to 10 Å, while the NB exhibited RMSD above 10 Å. Therefore, the RMSD evolution over time of MD simulations initiated from the best refined PELE pose allows quantitatively identifying SB among MB, WB and NB.

#### *2. HER2's pseudo-neuregulin binding site*

Before proceeding with the prospective HER3 study, we did an additional retrospective study on a similar protein, HER2. In this case, we had 1 MB and 2 WBs targeting HER2, and selected 3 Affilin® proteins targeting FINC (including 2 SBs and 1 MB) as NB.

Both PyDock and PELE effectively differentiated binders from NB (Figure SN13a and SN13b). Figures SN13c-e show the best PELE binding mode for each system, revealing that all but one WB target HER2's pseudo-neuregulin (pseudo-NRG) binding site. MD simulations starting from these refined poses successfully distinguished NB from WB and MB (Figure SN13f), reinforcing the pipeline's effectiveness in predicting binding interactions.

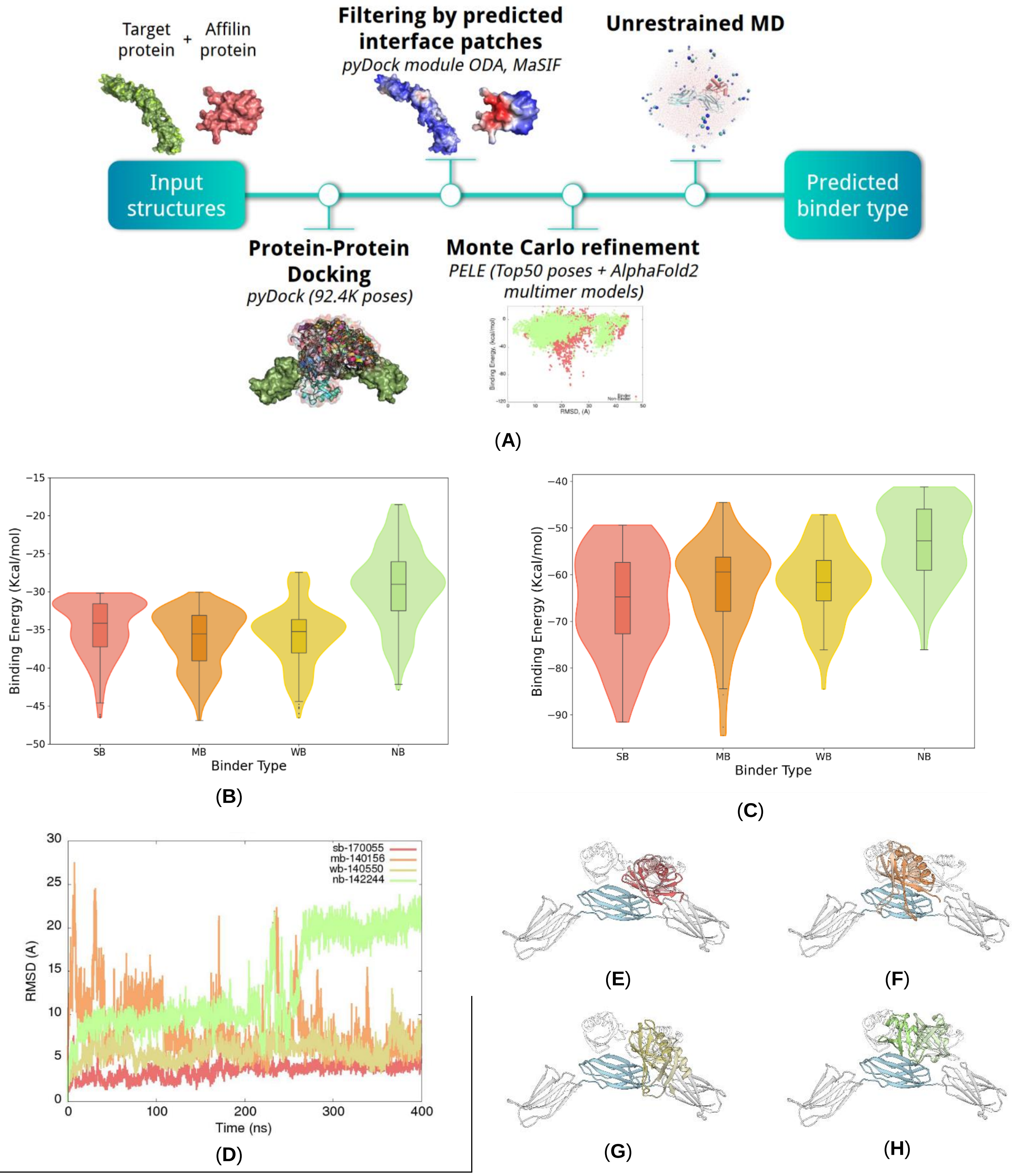


**Figure SN1.** Scheme of the MM pipeline utilized for identifying both SB and NB Affilin® binders targeted towards a specific protein. The pipeline's performance is showcased in the Affilin-FINC retrospective study, employing homology models for the Affilin® proteins. (**A**) The MM pipeline begins with input structures of the target and the Affilin® protein, sourced from AlphaFold2, homology modeling, or crystal structures. However, assessing the model's quality beforehand is crucial to ensure reliable results. Utilizing these input structures, 92,400 PPD poses are generated using PyDock. These poses are then filtered based on distance according to predicted interaction patches obtained by MaSIF and PyDock ODA. The top 50 non-repeated poses (L-RMSD between them greater than 2 Å) with the best PyDock energy are selected as input structures for Monte Carlo simulations using PELE. PELE refines the binding modes by introducing flexibility to the protein's backbone and side chains through soft random translations and rotations to the Affilin® protein. The best binding energy pose obtained from PELE refinement is then subjected to a Molecular Dynamics (MD) simulation to predict Affilin® binder types accurately. (**B**) Top 50 PyDock binding energy per system, grouped by binder type. (**C**) Top 10 PELE binding energy per system, grouped by binder type. (**D**) Affilin® protein's backbone L-RMSD evolution over time in the MD simulations. (**E, F, G, H**) Binding mode of the last frame of the SB, MB, WB, and NB, respectively. In white ribbons is depicted the reference FINC-Affilin® complex (PDB: 8PF0). (Color code in the figure: SB, red; MB, orange; WB, yellow; and NB, green).

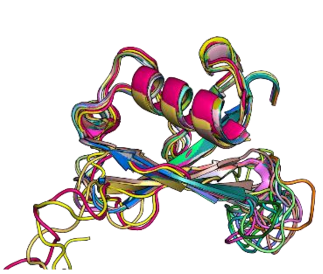

**(A)**

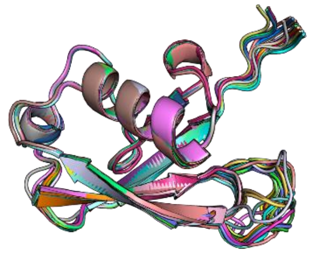

**(B)**

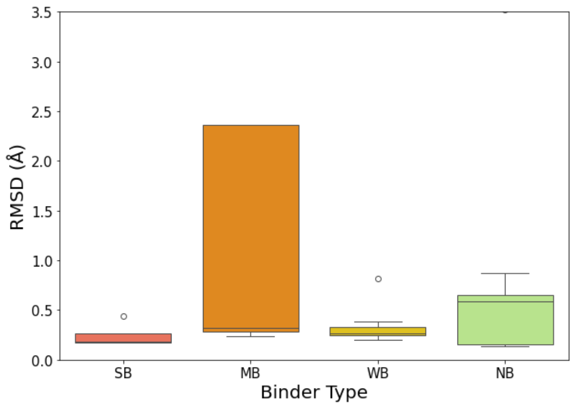


**(C)**

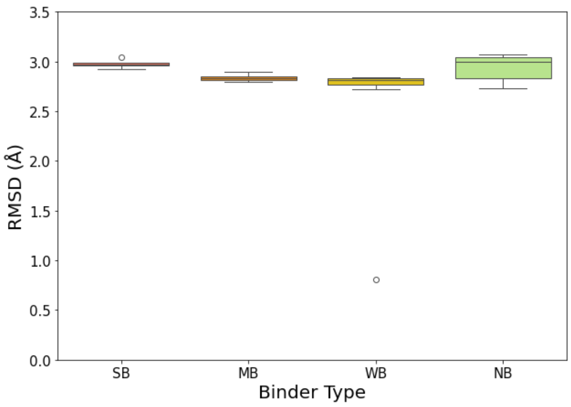


**(D)**

**Figure SN2.** Analysis of MODELLER and AlphaFold2 models. Superposition of models generated by (**A**) MODELLER and (**B**) AlphaFold2 on top of the Affilin® crystal. Cα-RMSD distribution of the (**C**) MODELLER and (**D**) AlphaFold2 best models grouped by binder type. MODELLER models were generated using PDB 8PEQ as template, while AlphaFold2 used default templates found by the MSA analysis.

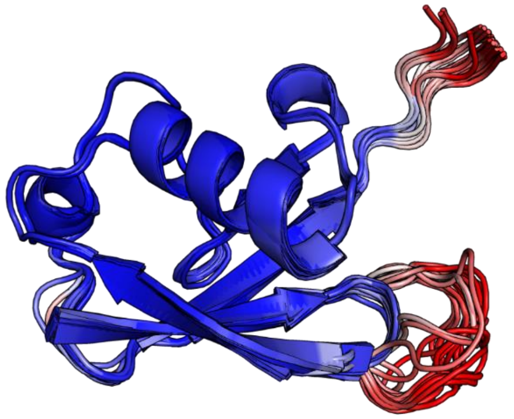

(A)

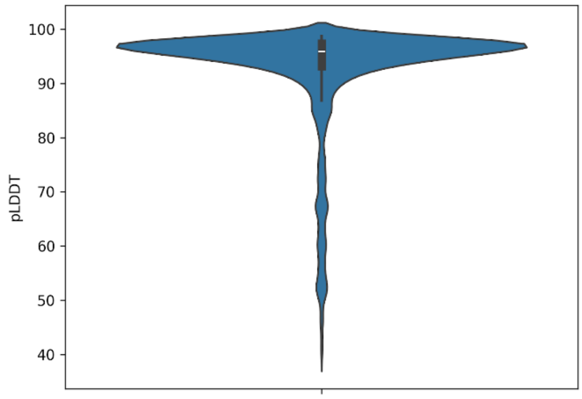


(B)

**Figure SN3.** AlphaFold2 models confidence scores for Affilin® proteins targeting FINC. (**A**) Superposition of the monomeric Affilin® proteins targeting FINC colored by pLDDT. Regions in blue are highly confident regions, while regions in red indicate low confidence. (**B**) Violin plot showing pLDDT distribution for all models.

**Table SN1.** AlphaFold2 templates and MSA depth for each modeled Affilin® protein targeting FINC. In total, only 10 different templates were used to generate all the models, which are: 2AL3_A, 5JNE_E, 5JNE_A, 5C23_B, 3V7O_B, 6EF3_U, 4ZYN_B, 5C1Z_A, 2Y5B_F, 5YCA_A.

| SequenceID | Templates (PDB_Chain) | | | | MSA Depth |
|---|---|---|---|---|---|
| NB_142276 | 2AL3_A | 5JNE_E | 5JNE_A | 6EF3_U | 3275 |
| MB_170059 | 5JNE_E | 5JNE_A | 2AL3_A | 5C23_B | 3206 |
| MB_140553 | 5JNE_E | 5JNE_A | 2AL3_A | 5C23_B | 3205 |
| WB_140552 | 5JNE_E | 5JNE_A | 2AL3_A | 5C23_B | 3224 |
| MB_140156 | 5JNE_E | 5JNE_A | 2AL3_A | 5C23_B | 3214 |
| MB_180095 | 5JNE_E | 5JNE_A | 2AL3_A | 3V7O_B | 3188 |
| NB_183389 | 5JNE_A | 5JNE_E | 2AL3_A | 3V7O_B | 3212 |
| WB_140550 | 5JNE_E | 5JNE_A | 2AL3_A | 3V7O_B | 3188 |
| SB_179931 | 2AL3_A | 5C23_B | 4ZYN_B | 6EF3_U | 3237 |
| MB_170070 | 5JNE_E | 5JNE_A | 2AL3_A | 3V7O_B | 3197 |
| WB_140157 | 5JNE_E | 5JNE_A | 2AL3_A | 5C23_B | 3164 |
| MB_170068 | 5JNE_E | 5JNE_A | 2AL3_A | 5C23_B | 3213 |
| NB_142244 | 5JNE_A | 5JNE_E | 2AL3_A | 3V7O_B | 3285 |
| NB_140127 | 5JNE_A | 5JNE_E | 2AL3_A | 3V7O_B | 3296 |
| SB_170058 | 5JNE_E | 5JNE_A | 2AL3_A | 6EF3_U | 3322 |
| NB_142308 | 5JNE_E | 5JNE_A | 2AL3_A | 3V7O_B | 3234 |
| MB_179959 | 6EF3_U | 4ZYN_B | 5C23_B | 5YCA_A | 3220 |
| MB_170061 | 5JNE_E | 5JNE_A | 2AL3_A | 5C23_B | 3200 |
| MB_170067 | 2AL3_A | 5JNE_A | 5JNE_E | 5C23_B | 3240 |
| WB_140554 | 5JNE_E | 5JNE_A | 2AL3_A | 5C23_B | 3246 |
| SB_170056 | 5JNE_E | 5JNE_A | 2AL3_A | 3V7O_B | 3278 |
| NB_183167 | 2AL3_A | 5JNE_E | 5JNE_A | 5C23_B | 3314 |
| NB_140128 | 2AL3_A | 5JNE_E | 5JNE_A | 5C23_B | 3284 |
| NB_142238 | 2AL3_A | 5JNE_E | 5JNE_A | 5C23_B | 3306 |
| NB_140129 | 5JNE_E | 5JNE_A | 2AL3_A | 3V7O_B | 3292 |
| NB_142375 | 2AL3_A | 5JNE_A | 5JNE_E | 5C23_B | 3247 |
| NB_142285 | 5JNE_E | 5JNE_A | 2AL3_A | 3V7O_B | 3306 |
| WB_44013 | 5C23_B | 5C1Z_A | 2Y5B_F | 2AL3_A | 3034 |
| MB_180017 | 5JNE_E | 5JNE_A | 2AL3_A | 3V7O_B | 3200 |
| MB_140551 | 5JNE_E | 5JNE_A | 2AL3_A | 5C23_B | 3220 |
| WB_140549 | 2AL3_A | 5JNE_E | 5JNE_A | 5C23_B | 3198 |
| MB_170069 | 5JNE_E | 5JNE_A | 2AL3_A | 5C23_B | 3219 |
| WB_170064 | 5JNE_E | 5JNE_A | 2AL3_A | 5C23_B | 3230 |
| SB_170055 | 5JNE_A | 5JNE_E | 2AL3_A | 3V7O_B | 3176 |
| SB_170062 | 5JNE_A | 5JNE_E | 2AL3_A | 3V7O_B | 3221 |

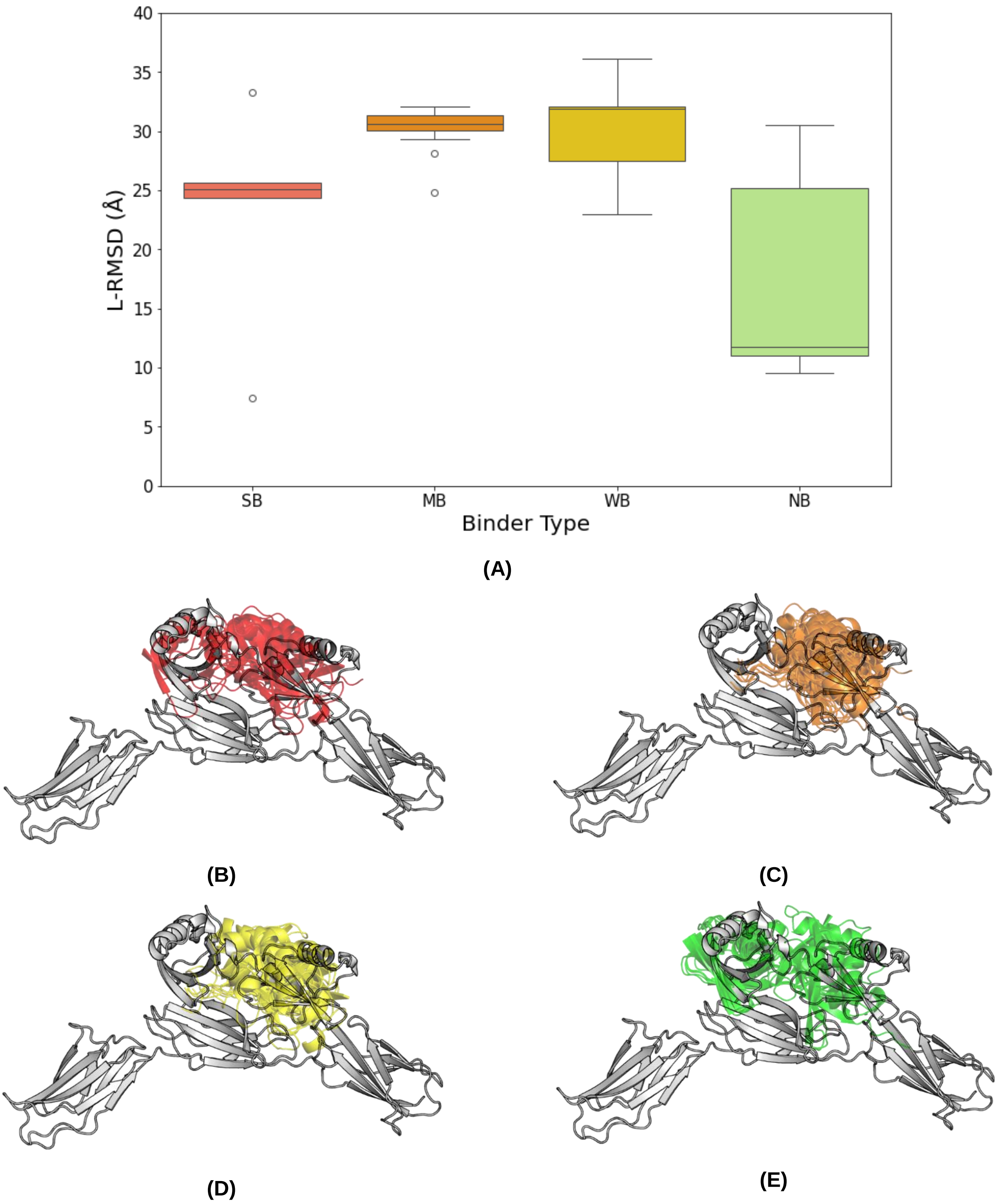


**Figure SN4.** Superposition results of the AlphaFold2 Multimer models of monomeric Affilin® protein targeting FINC. (**A**) L-RMSD distribution of AlphaFold2 Multimer best model with respect to the reference crystal (PDB: 8PEQ). (**B, C, D, E**) Visualization of the AlphaFold2 multimer predicted binding modes grouped by binder type: SB (red), MB (orange), WB (yellow) NB (green). The reference structure is displayed in white ribbons.

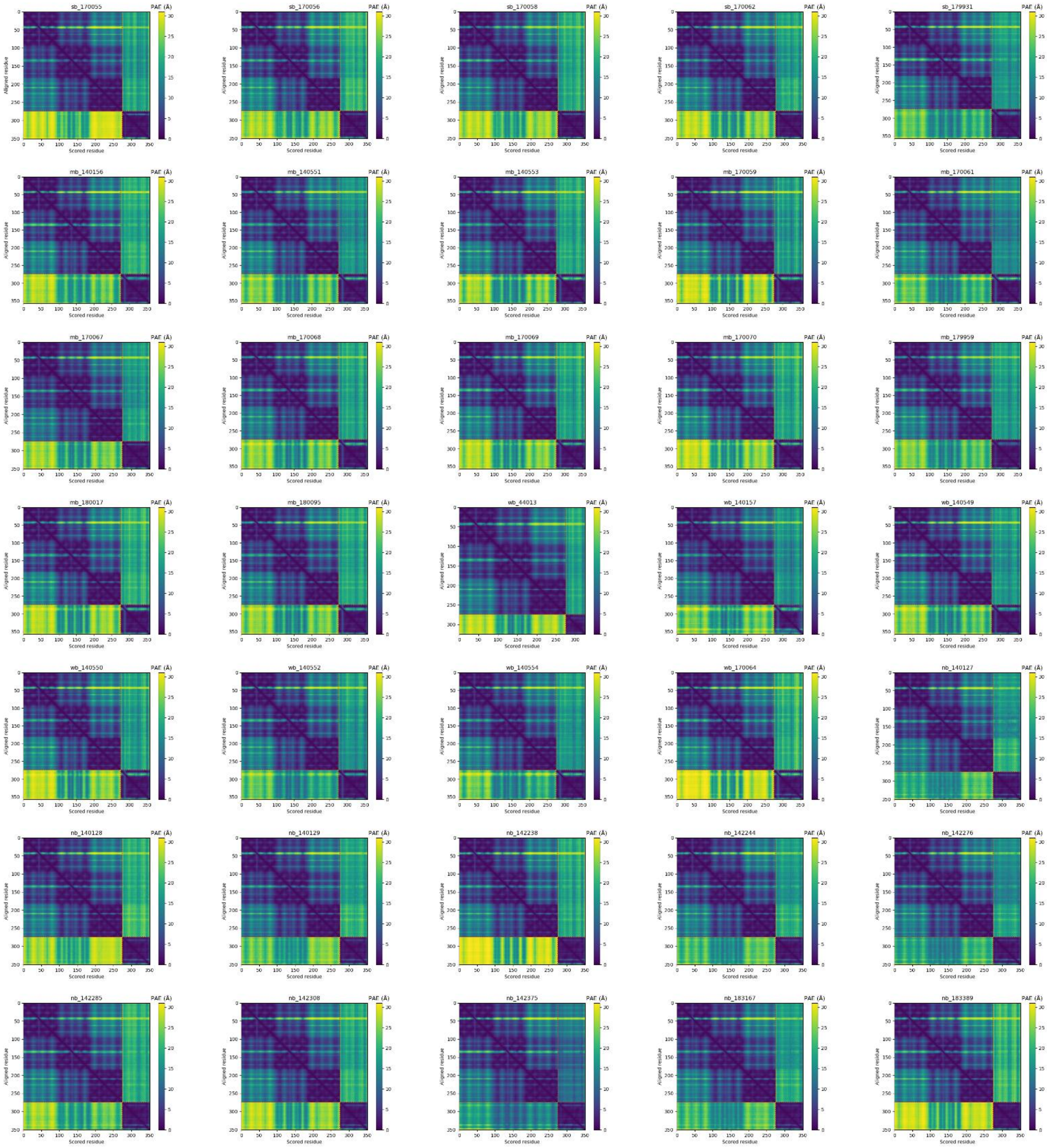


**Figure SN5.** Predicted Aligned Error (PAE) plots for the best AlphaFold2 Multimer model for each Affilin-FINC complex. The colors estimate the expected distance error in Å on the scored residue's position X (x-axis) when the predicted and the true structures are aligned on residue Y (y-axis). Yellow colors indicate higher expected distance while blue lower expected distances. Red dashed lines indicate FINC last residue (residue 274). Therefore, the upper left block has the PAE values within the FINC protein, while the bottom right within the Affilin® protein. The upper right block estimates the distance error of the Affilin® protein residues when aligning FINC, while the bottom left block is the estimated error of FINC residues when aligning the Affilin® protein.

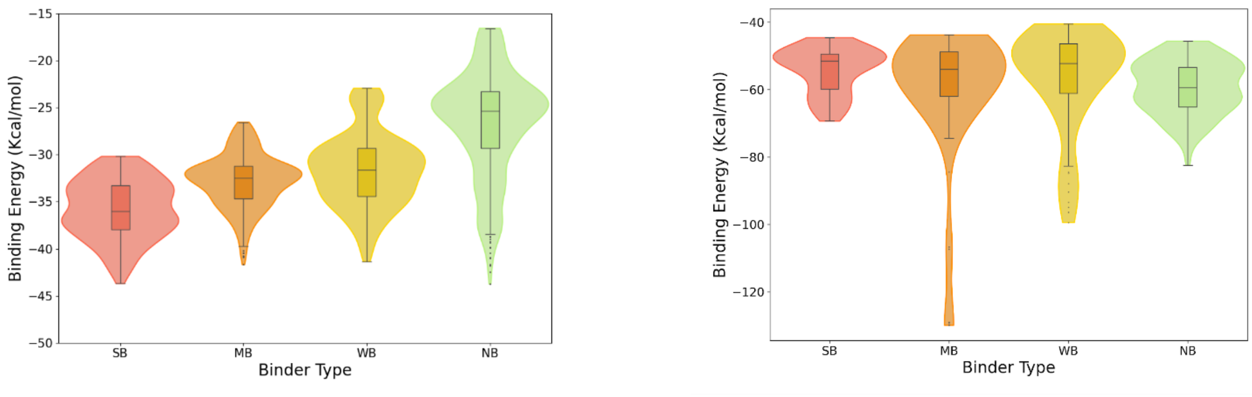


(A) (B)

**Figure SN6.** Protein-Protein Docking and refinement binding energy distributions when using AlphaFold2 for Affilin® proteins targeting FINC (**A**) Top 50 PyDock binding energies per binder grouped by binder type. (**B**) Top 10 PELE binding energies per binder grouped by binder type.

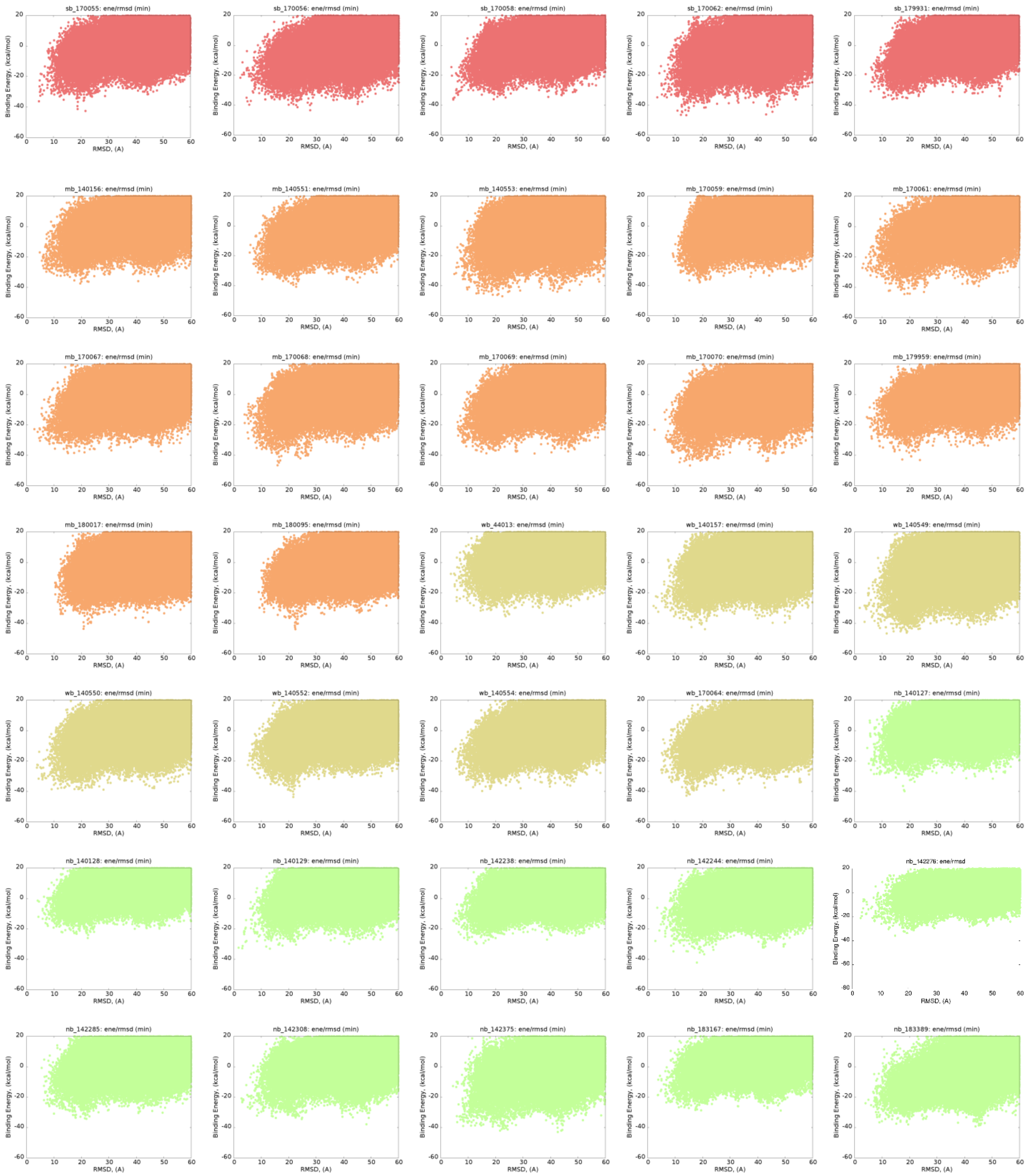


**Figure SN7.** PyDock energy plots for all tested complexes FINC-Affilin® using MODELLER models. PyDock energy in function of Affilin® L-RMSD with respect to the crystal when docking MODELLER models. Red are SBs, orange MBs, yellow WBs, and green NBs.

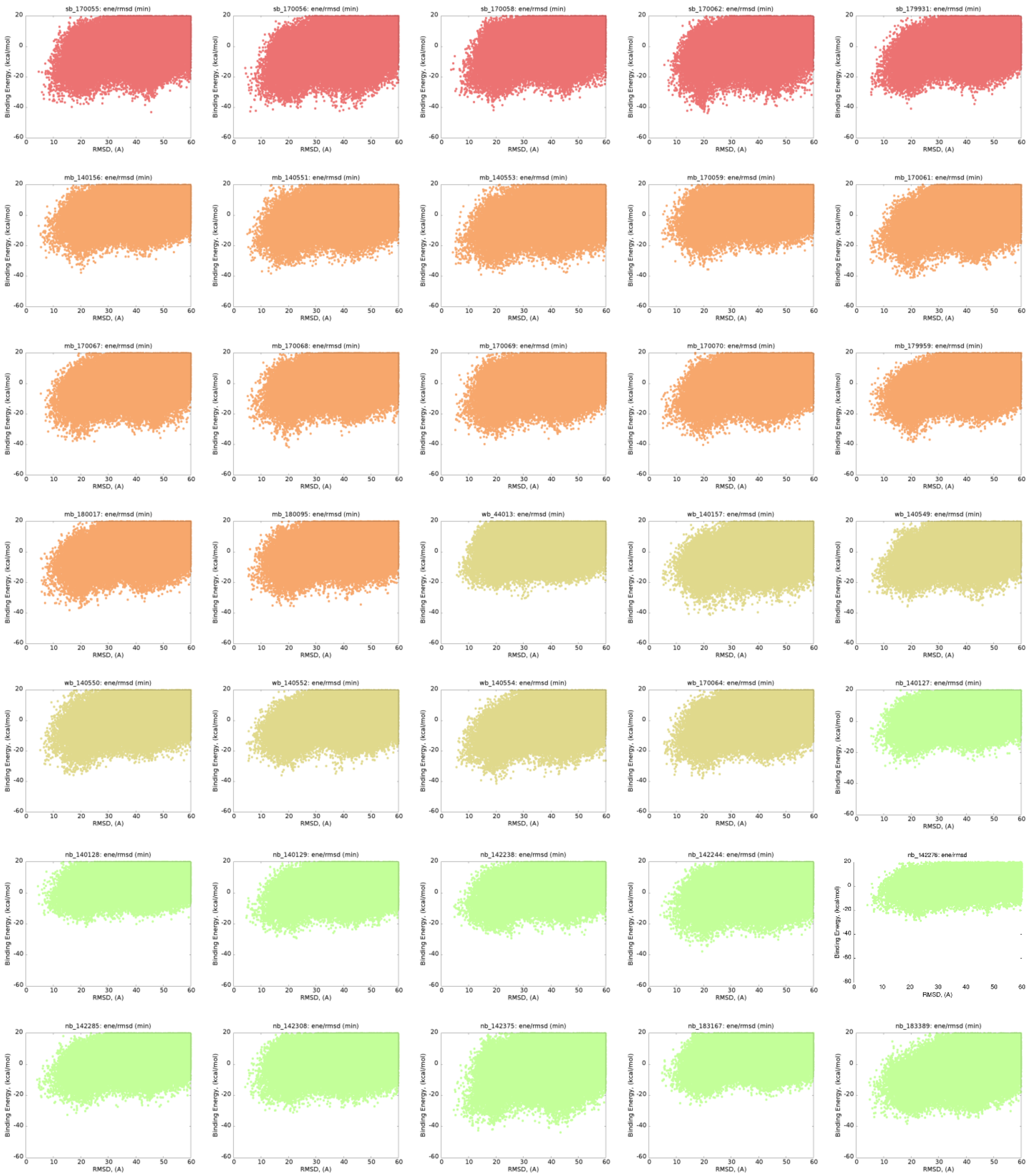


**Figure SN8.** PyDock energy plots for all tested complexes FINC-Affilin® using AlphaFold2 models. PyDock energy in function of Affilin® L-RMSD with respect to the crystal when docking AlphaFold2 models. Red are SBs, orange MBs, yellow WBs, and green NBs.

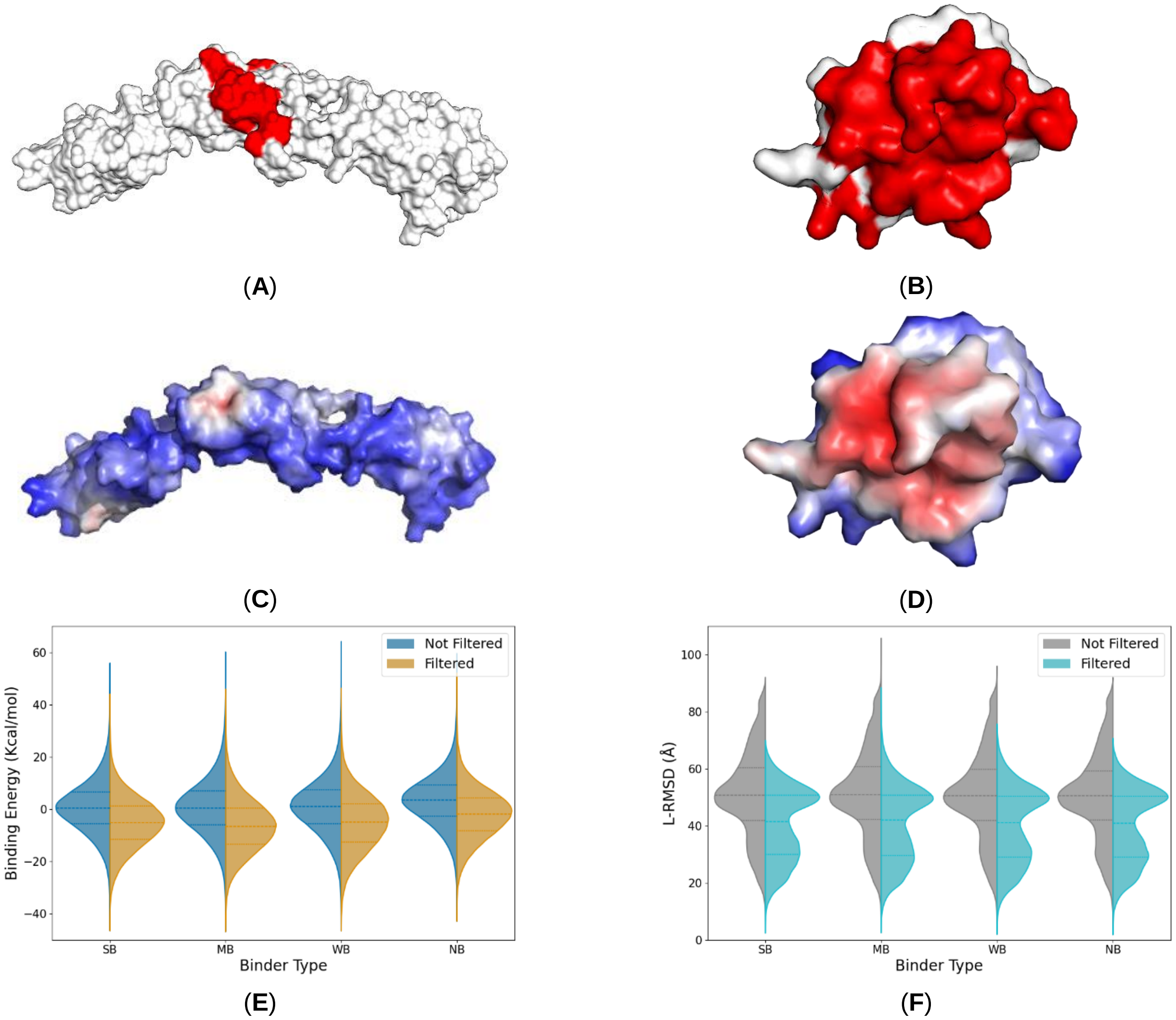


**Figure SN9.** Comparison of predicted interacting patches and resulting energy and L-RMSD distributions after applying the interacting patch filter. (**A, B**) PyDockODA predictions in red for the FINC and an Affilin® protein, respectively. (**C, D**) MaSIF-site predictions for the FINC and the same Affilin® protein, respectively. Red regions indicate likely interacting regions, while blue regions suggest a lower interacting likelihood. (**E**) PyDock energy and (**F**) L-RSMD (respect the reference FINC-Affilin® crystal (PDB: 8PF0)) distribution of all systems poses before (left distributions) and after applying the interacting patch filter (right) when using Affilin® homology models.

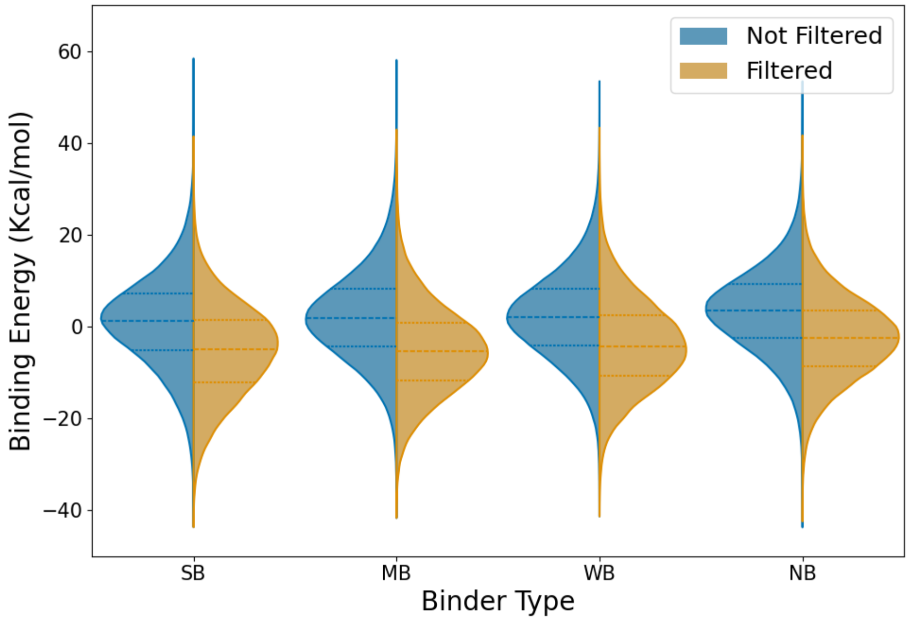


**(A)**

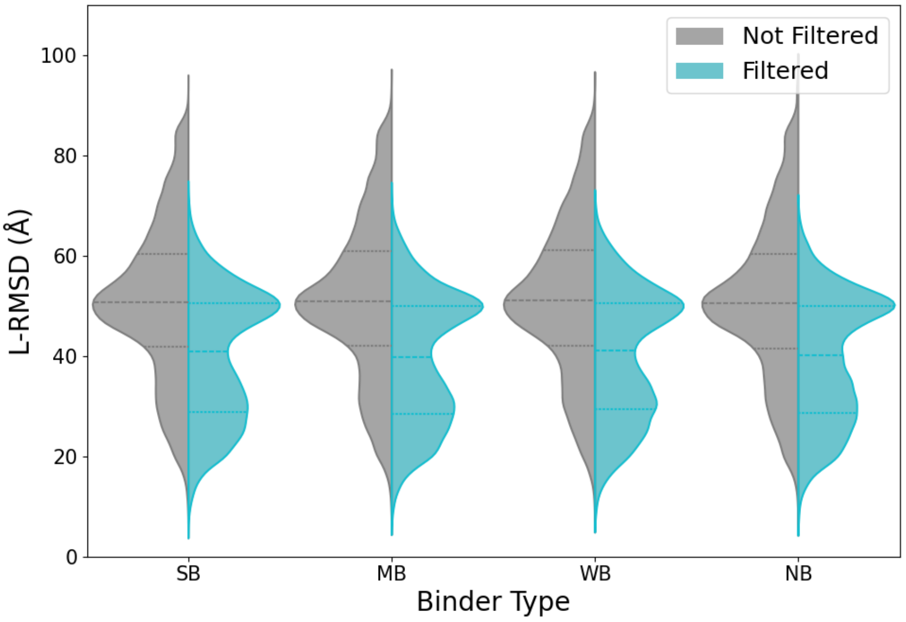


**(B)**

**Figure SN10.** Resulting energy and L-RMSD distributions before (left distributions) and after (right distributions) applying the interacting patch filter to PPD poses obtained using AlphaFold2 Affilin® models. (**A**) PyDock energy distributions. (**B**) L-RMSD distributions compared with the reference FINC-Affilin® crystal (PDB: 8PEQ) for all the poses per each system.

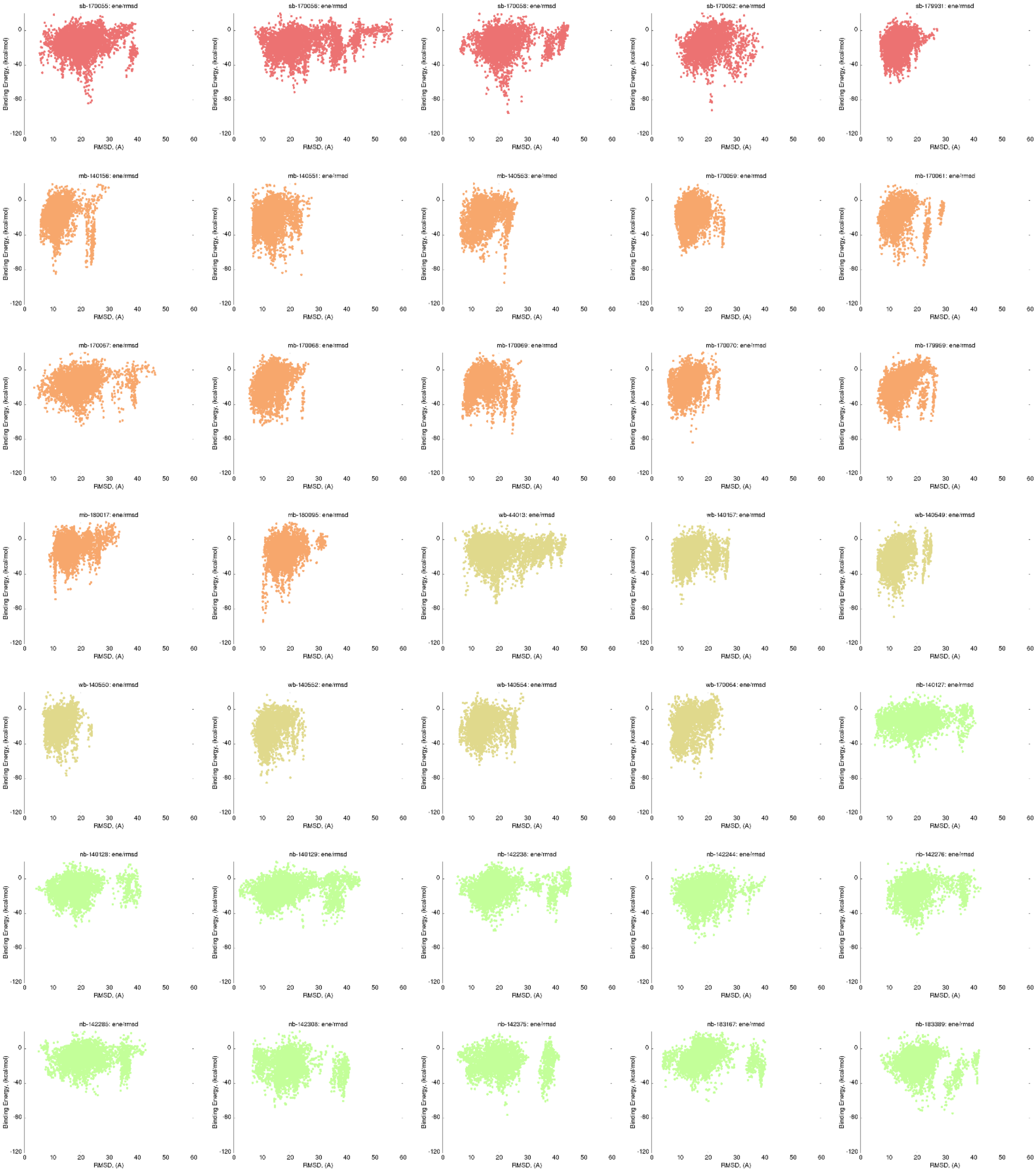


**Figure SN11.** PELE binding energy plots for all tested complexes FINC-Affilin® using MODELLER models. PELE binding energy in function of Affilin® L-RMSD with respect to the crystal when using Affilin® MODELLER models plus the 5 AlphaFold Multimeter models. Red are SBs, orange MBs, yellow WBs, and green NBs.

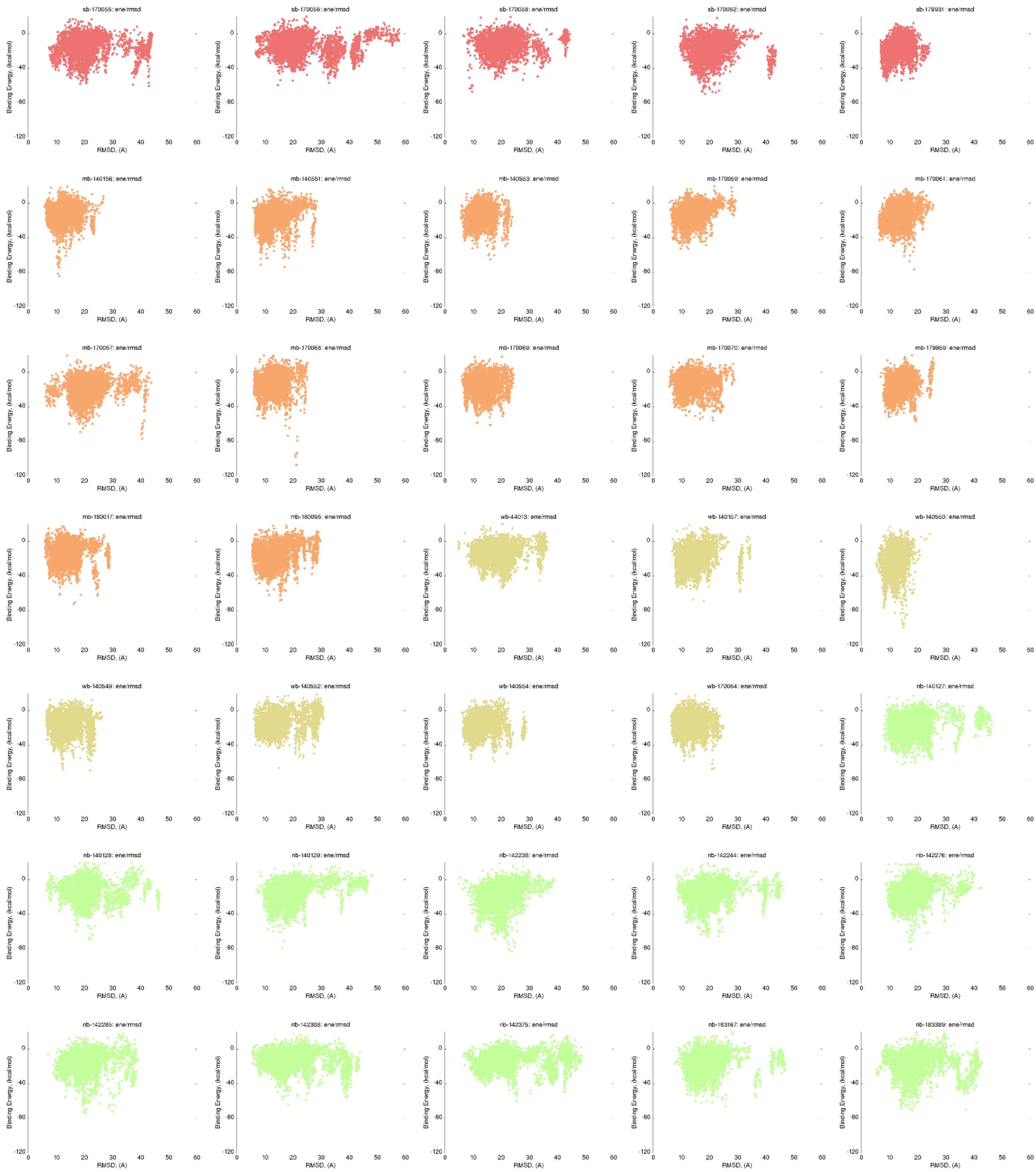


**Figure SN12.** PELE binding energy plots for all tested complexes FINC-Affilin® using AlphaFold2 models. PELE binding energy in function of Affilin® L-RMSD with respect to the crystal when using Affilin® AlphaFold2 models plus the 5 AlphaFold Multimeter models. Red are SBs, orange MBs, yellow WBs, and green NBs.

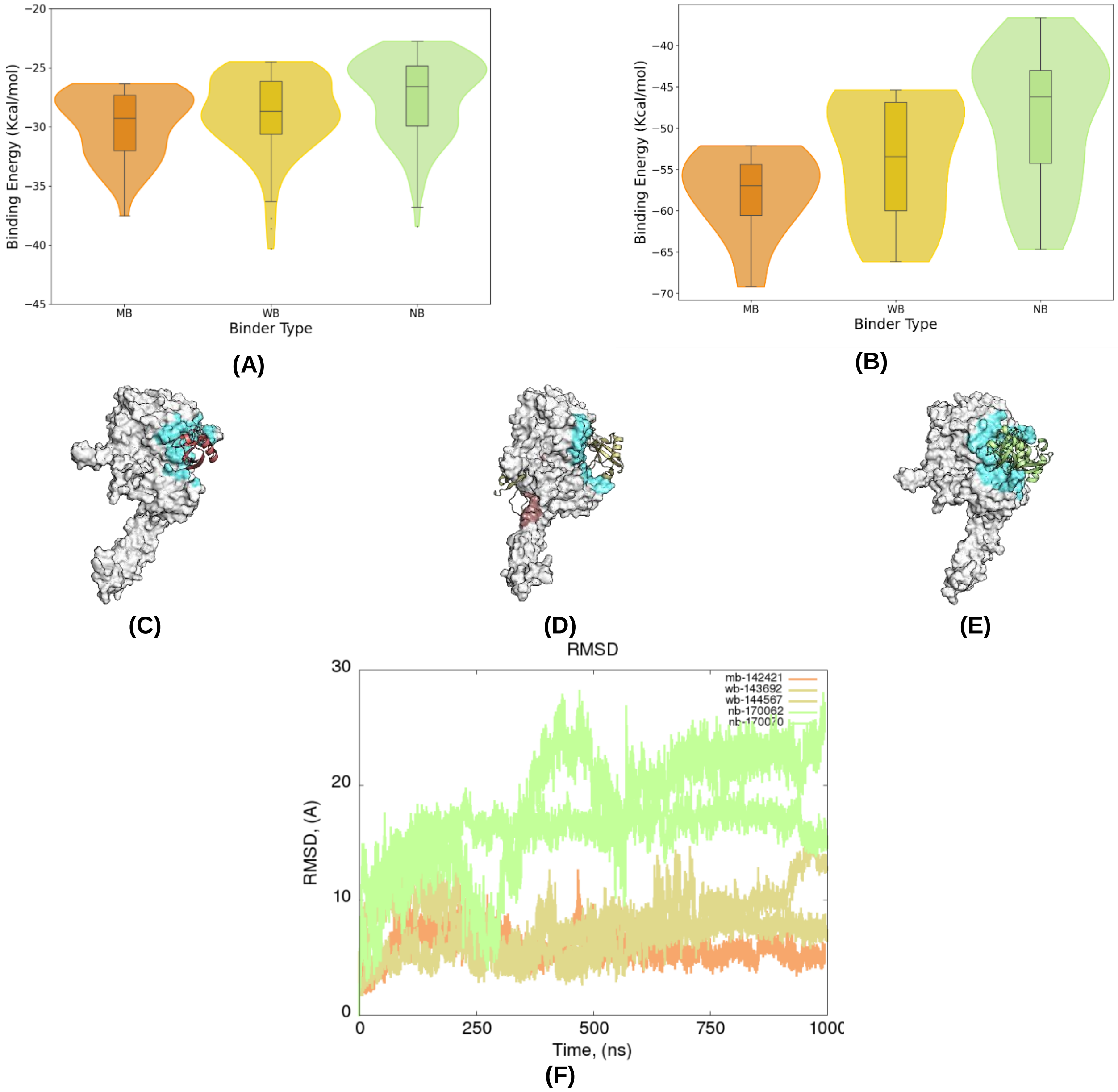


**Figure SN13.** Results of our MM pipeline on HER2 retrospective study. (**A**) Binding energy distribution of the top 50 pyDock filtered poses per system and grouped by binder type. (**B**) Binding energy distribution of the top 10 PELE poses per system and grouped by binder type. (**C, D, E**) Best PELE binding modes for each Affilin® protein for MB, WB and NB, respectively. WB view has been rotated 90º to the right to show all the binding modes found. HER2 is displayed as a white surface and its pseudo-neuregulin (pseudo-NRG) binding site is highlighted in blue. (**F**) Affilin® protein's backbone L-RMSD evolution over time along the MD simulations (MB, orange; WBs, yellow; and NBs, green).
